# Sequence-dependent conformational and mechanical landscapes of double-stranded nucleic acids

**DOI:** 10.64898/2026.08.09.740023

**Authors:** Rahul Sharma, Alessandro S. Patelli, Raushan Singh, Daiva Petkevičiūtė-Gerlach, Oscar Gonzalez, John H. Maddocks

**Affiliations:** Institute of Mathematics, École Polytechnique Fédérale de Lausanne, Lausanne, Switzerland; Department of Mechanical Engineering, IIT Madras, Chennai, India; Department of Applied Mathematics, Kaunas University of Technology, Kaunas, Lithuania; Department of Mathematics, University of Texas, Austin, USA

**Keywords:** Sequence-dependence, DNA mechanics, RNA, DNA:RNA hybrids, coarse-grain models, epigenetic modifications, Groove width, CTCF binding sites, methylation, hydroxymethylation, persistence length

## Abstract

The sequence-dependent mechanical landscapes of double-stranded nucleic acids (dsNAs) remain largely unexplored beyond canonical dsDNA. We describe cgNA+, a coarse-grained predictive model of the mechanics of dsRNA, DNA:RNA hybrids, and epigenetically modified dsDNA, all parameterised from 1.26 milliseconds of atomistic simulations. cgNA+ predicts non-local sequence-dependent equilibrium shape and stiffness with errors an order of magnitude smaller than sequence-variability, while enabling exploration of numbers/lengths of sequences inaccessible to atomistic simulation. We show that dsNA equilibrium shape is strongly influenced by flanking sequence up to octamer context, with flexible dimer-steps more context-sensitive. C_p_G-modification alters equilibrium shape comparable to changes caused by single-nucleotide polymorphisms. Groove-width analysis across dsNA decamers reveals strong sequence dependence, reflecting the differing characteristic helical-geometry of dsDNA and dsRNA, whereas DRHs exhibit mixed behaviour depending on DNA-strand pyrimidine content. CTCF binding sites exhibit a distinct groove-width signature. Persistence-length spectra from ∼ 9 million sequences indicate that dsRNA is stiffer than dsDNA, whereas DRH exhibits intermediate stiffness modulated by DNA-strand pyrimidine content. Persistence length increases upon C_p_G-modification, but decreases on hypermodification. Overall, the cgNA+ model enables a first, highly accurate (compared to atomistic simulations), very large-scale, comparative study of sequence-dependent mechanics both within and across dsNA classes, demonstrating previously hidden regulatory layers.

**Graphical Abstract:** 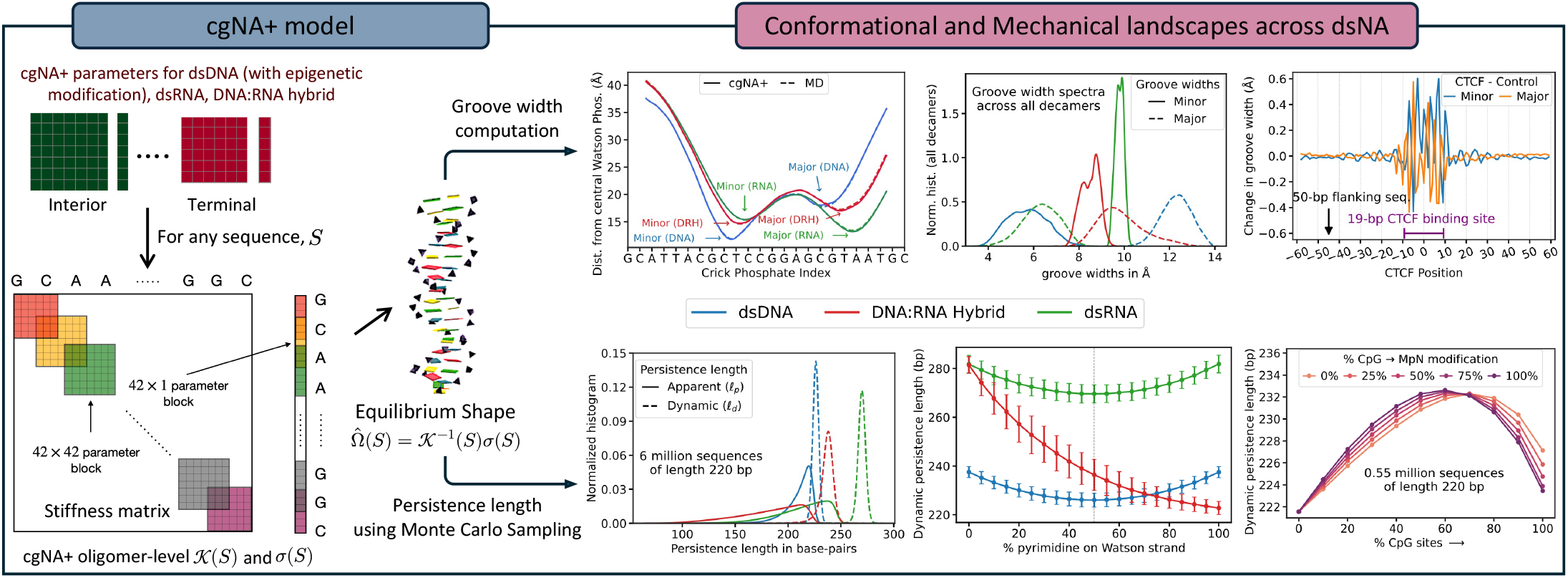

## Introduction

Double-stranded nucleic acids (dsNA) not only encode genetic information but also possess intrinsic structural and mechanical properties that are strongly modulated by sequence [1–14]. These properties, including intrinsic shape, flexibility, bending propensity, torsional response, and susceptibility to local deformations, are key determinants of dsNA biological functions. The functional importance of sequence-dependent mechanics is particularly well established for double-stranded DNA (dsDNA). Variations in dsDNA curvature and flexibility influence nucleosome positioning and chromatin organisation [7, 8], regulate DNA looping involved in transcriptional control [3, 15], and underlie indirect readout mechanisms in protein– DNA recognition [16]. Structural and mechanical features such as groove geometry, bending propensity, and stiffness further modulate the protein-binding specificity and affinity [17, 18]. Furthermore, epigenetic modifications, particularly cytosine methylation and hydroxymethylation at C_p_G sites, introduce an additional layer of regulatory complexity by altering both the chemical and mechanical properties of DNA [4, 19, 20], thereby influencing gene regulation, chromatin accessibility, genomic imprinting, development, and disease [4, 8, 19, 21–23].

Beyond dsDNA, the broader dsNA family also plays important biological roles. Double-stranded RNA (dsRNA) is central to RNA interference, constitutes the genome of many viruses, and forms structured regulatory RNA elements [13, 24, 25]. DNA:RNA hybrids (DRH) arise in a wide range of biological contexts, including reverse transcription, DNA replication, transcription-associated R-loop formation, DNA repair, and CRISPR–Cas target recognition [14, 26, 27]. RNase H selectively degrades the RNA strand of DRH, possibly through recognition of its distinct intermediate helical structure and characteristic minor groove width [28, 29].

Experimental studies, including X-ray crystallography [9, 10, 30], cryogenic electron microscopy [11, 12], optical tweezers [31, 32], and magnetic tweezers [33–35] have revealed sequence-specific duplex flexibility and deformation. More recently, high-throughput cyclization experiments demonstrated that DNA looping depends strongly on sequence [3, 4]. Complementary insights have emerged from molecular dynamics (MD) simulations, which provide atomic-resolution descriptions of sequence-dependent structure and fluctuations [36–48]. Furthermore, numerous studies have shown that the structure and flexibility of a base-pair step can depend strongly on flanking sequence context extending well beyond the immediate dimer [5, 6, 39– 41, 49, 50]. Collectively, these studies establish that dsNA mechanics is inherently sequence-dependent and reflects interactions that extend beyond local nearest-neighbour effects.

Despite this progress, a comprehensive understanding of sequence-dependent mechanical landscapes remains elusive. Experimental approaches are generally limited to relatively small subsets of sequence space, e.g., X-ray crystallography [9, 10, 30], or do not provide detailed structural characterization, e.g., high-throughput cyclization experiments [3, 4]. Computational methods such as MD simulations can reveal the molecular origins of sequence-dependent behaviour but remain computationally demanding, restricting the length and number of sequences that can be explored. As a result, systematic investigation of the vast sequence space associated with dsNA remains beyond the reach of current experimental and atomistic MD approaches.

Coarse-grained models offer a promising route to bridging this gap by retaining only the degrees of freedom most relevant to nucleic-acid mechanics, thereby enabling efficient exploration of large sequence ensembles. Several models have been developed to investigate sequence-dependent nucleic acid structure and mechanics, ranging from coarse-grained simulations to machine-learning-based potentials [51–54]. A comprehensive overview can be found in [42]. Classical worm-like chain models successfully describe the average behaviour of homo-polymers [55, 56], whereas rigid-base-pair coarse-grained frameworks capture local, dimer-level sequence-dependent structure and elasticity [10, 57]. However, these models are inherently limited in their ability to represent accurately both local and non-local couplings and sequence-dependent effects when benchmarked against direct MD simulation.

Many rigid-base-pair coarse grain models assume, either implicitly or explicitly, that the inverse covariance matrix of the equilibrium distribution for their rigid-base pair coordinates is 6 × 6 block diagonal. Such a sparsity pattern is associated with there being no direct energetic coupling between degrees of freedom in one base-pair junction and any other, even the two nearest neighbour junctions. As pointed out in [58], this assumption is rather inaccurate when compared with data extracted directly from MD simulations. In contrast, it was also observed in [58] that the inverse covariance matrix for the degrees of freedom associated with nearest-neighbour interaction in a slightly finer grain model, namely a rigid base description was much closer to the actual sparsity patterns observed directly from MD. This observation led to the development of the predictive cgDNA rigid-base model [59–62]. The cgNA+ model (as originally developed for dsDNA in Patelli’s thesis [63] and refined and extended to other dsNA in Sharma’s thesis [64]) adopts the even finer grain description of rigid bases plus rigid phosphate groups. An interactive web implementation cgNA+web [65] is available at https://cgdnaweb.epfl.ch. This cgNA family of models (including cgNA+) has already been applied to studies of persistence length [66], nucleosome wrapping/unwrapping [67–70], a genome-wide search for mechanically exceptional sequences [71], and sequence-dependent DNA minicircle shapes and energies [72].

This article first comprehensively describes the cgNA+ model. The model is parameterised from extensive all-atom MD simulation libraries of total duration 580 *µ*s for dsDNA, including epigenetically modified base-pair steps, and 340 *µ*s each for dsRNA and DRH. We benchmark the cgNA+ model against independent test MD simulations, revealing how remarkably accurate it is in predicting both sequence-dependent equilibrium shape and stiffness across dsDNA, dsRNA, DRH, and epigenetically modified dsDNA. In particular, the model accurately captures non-local sequence dependence in the equilibrium shape of dsNA, often extending to the hexamer context or beyond, with prediction errors that are an order of magnitude or more lower than the observed sequence-variability. It should be noted that the accuracy of cgNA+ has been verified against MD simulations. The question of how closely MD simulations represent reality is a separate issue.

Buoyed by these favourable verification tests of accuracy compared to MD, we then leverage the computational efficiency of cgNA+ to scan large ensembles of long dsNA sequences of all types, in order to investigate and contrast their sequence-dependent mechanical properties. We construct comparative maps of equilibrium shape, groove geometry, and persistence length, for all dsNA classes, and over ensembles of millions of different sequences of lengths up to and beyond 200 bp, with the goal of demonstrating that the cgNA+ model is well capable of investigating biologically pertinent questions.

## Materials and Methods

### The cgNA+ model

cgNA+ is a sequence-dependent, coarse-grained model that efficiently and accurately predicts first and second moments of the configuration space equilibrium distribution (or equivalently the groundstate, or average shape, and stiffness) for arbitrary-sequence fragments of double-stranded nucleic acids, including dsDNA in an epigenetically modified alphabet, dsRNA, and DRH (DNA:RNA Hybrid). The explicit level of coarse-grain description in the model is that each base and each phosphate group are approximated as rigid bodies. The equilibrium distribution on the conformational space of this coarse grain description of a dsNA molecule is then approximated as a multivariate Gaussian distribution expressed in a set of relative coordinates between this system of rigid bodies. The model parameter sets that allow prediction of the equilibrium distribution for arbitrary sequence fragments of each of the three types of dsNA is dimer-sequence-dependent, but nevertheless captures much longer sequence-dependence of the structure. (Examples are provided below.) Each parameter set is estimated from an extensive library of all-atom molecular dynamics (MD) simulations of a carefully chosen set of sequence fragments. The logical structure of the cgNA+ model is highlighted in Fig. 1. In the main text and SI of this presentation we provide a condensed, but self-contained, description of each of these modelling steps, but our focus here is primarily on predictions arising from the cgNA+ model that we believe to be biologically interesting. A more comprehensive discussion, including the complete underlying computational foundations, and estimation of errors in the various approximations, can be found in the two PhD theses [63, 64].

**Figure 1.**
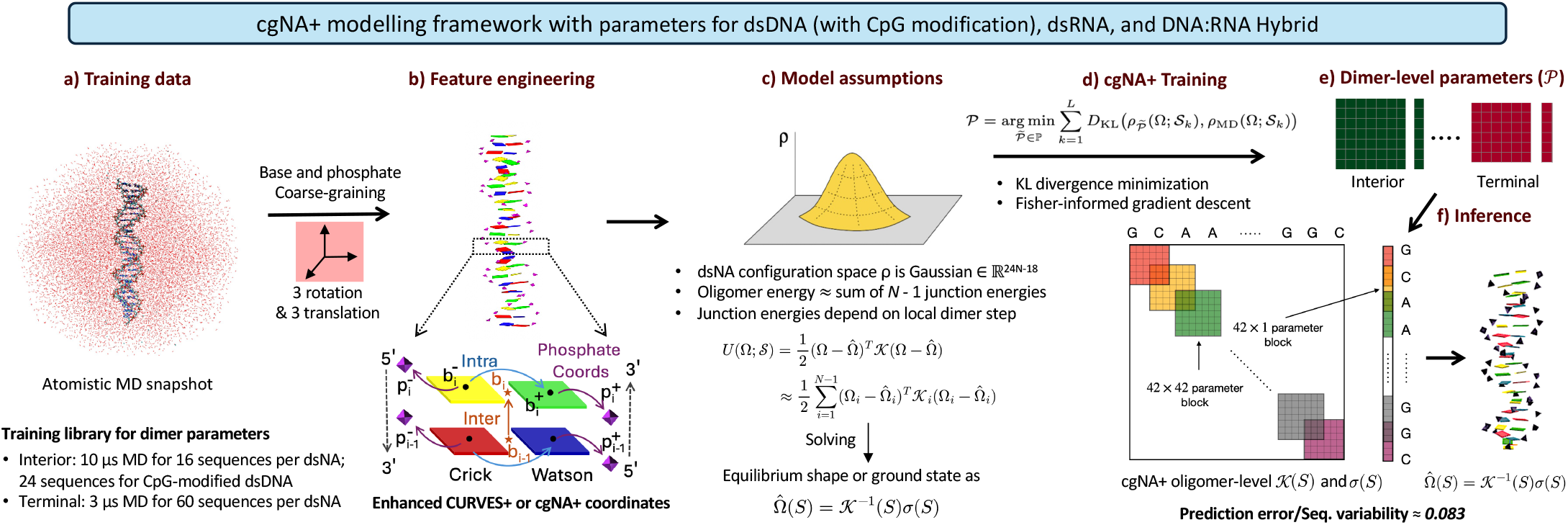
Schematic overview of the cgNA+ framework. **(a)** cgNA+ parameters are derived from extensive atomistic MD simulations of diverse sequences (training libraries provided in Tables S1–S3) and are independently trained for dsDNA, epigenetically modified dsDNA, dsRNA, and DRH. **(b)** Atomistic MD trajectories are coarse-grained by fitting rigid-body reference frames to nucleobases and phosphate groups and expressing the resulting configurations in the enhanced Curves+ (or cgNA+) internal coordinates. 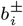 and 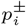 denote the base and phosphate frames on the Crick (denoted by superscript −) and Watson (denoted by superscript +) strands at the *i*^th^ base-pair level, respectively, while *b*_*i*_ denotes the corresponding averaged base-pair frame. **(c)** The conformational space of a dsNA is represented as a multivariate Gaussian distribution in cgNA+ internal coordinate space. The corresponding oligomer-level quadratic free-energy is approximated as sequence-dependent local junction contributions (Section S3). **(d)** Model parameters are determined by minimising the total Kullback–Leibler (KL) divergence between cgNA+ predicted and MD-derived Gaussian estimates over all training sequences using a Fisher-informed gradient-descent optimisation procedure. **(e)** Schematically shows the parameters consisting of dimer-dependent stiffness blocks and stress-like vectors. The terminal parameter blocks 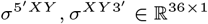 and 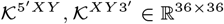 and the interior parameter blocks *σ*^*XY*^ ∈ R^42×1^ and K^*XY*^ ∈ R^42×42^ . **(f)** For any input sequence, these dimer-dependent parameters are assembled to construct the oligomer-level stress-like vector and stiffness matrix (Section S3) which is then used to compute the equilibrium shape. Although the parameterization is local, cgNA+ naturally captures non-local sequence-dependence through mechanical coupling between neighbouring junctions with average prediction error in equilibrium shape and stiffness an order in magnitude smaller than sequence-variability for test sequences (Table S6) in terms of KL divergence (Eq. 8).

#### Atomistic MD Simulations for Training cgNA+ parameter sets

The cgNA+ model includes independently trained parameter sets (described below) for canonical dsDNA, dsRNA, DRH, and epigenetically modified dsDNA, with each parameter set being derived from distinct all-atom MD simulations. Furthermore, the model contains distinct parameter blocks for interior and terminal dimer steps. For many biological applications it is only the interior parameter blocks that are of most interest, but the introduction of independent end parameter blocks allows the interior blocks to be estimated more accurately, with the end fraying of the dsNA structure being properly, and separately, accounted for. In fact, the end blocks can be estimated quite straightforwardly from MD simulations of a separate library of very short, and therefore not very computationally intensive, dsNA sequences. For estimating interior dimer parameters along with specifically (the most stable) GpC dimer step end parameters, we used a set of palindromic (i.e., reading strand invariant) sequences embedded in GpC end dimers (16 sequences of length 24 base pairs each for dsDNA, dsRNA, and DRH and 24 additional sequences for training parameters for methylated and hydroxymethylated C_p_G steps) designed to compactly pack all possible tetramers, and to have balanced statistics for all dimers and trimers (Tables S1-S For training the remaining fifteen end-dimer parameter blocks, we used 60 sequences of length 12 base pairs (Table S3). We performed additional MD simulations with identical protocols for several other dsNA sequences (Tables S1-S2) to create a diverse testing library for benchmarking cgNA+ predictions. And no test sequence was used in parameter estimation.

The cgNA+ framework (but definitely not the parameter set) is designed to be agnostic to MD protocol. Differences in MD protocol can arise for either physical reasons, e.g., different species and concentration of counter ions, or solvent temperature, or due to modelling differences in the underlying MD potentials. In fact the cgNA+ parameter estimation procedure is sufficiently accurate that differences due solely to differences between MD inputs (differing only in their associated protocols) can be clearly identified [65]. Nevertheless, here we focus on predictions for differing dsNA, all arising from one particular cgNA+ parameter set that we name PS2 (naming convention follows the cgNA+web interactive interface, https://cgdnaweb.epfl.ch). The specific MD simulations leading to the PS2 cgNA+ parameter set were performed using the parmbsc1 [73] and the OL3 [74] corrections to the parmbsc0 and parm99 force fields [75, 76] for dsDNA and dsRNA, respectively, while DRH was described using a combination of both. Additional parameters were included for methylated and hydroxymethylated cytosine [19, 20]. The dsNA molecule was solvated in explicit TIP3P water [77] using a truncated octahedral box with a 10 Å solvent buffer and neutralised with K^+^ ions. KCl was further added to obtain an ionic strength of approximately 150 mM using Joung–Cheatham ion parameters [78]. The MD production runs were 10 *µ*s for each dsNA sequence in Tables S1-S2 and 3 *µ*s for each dsNA sequence in Table S3 using AMBER [79]. We refer to Section S1 and the thesis [64] for more detail on the MD simulations.

#### Coarse-Graining: Atomistic MD snapshots to cgNA+ Coordinates

Atomistic MD snapshots were transformed into cgNA+ coarse-grained rigid body representation snapshots through a least-squares fitting procedure (Table S4), in which rigid-body reference frames were assigned to the nucleobases and phosphate groups (Figure 1b; Figure S1). Sugars and solvent are included only implicitly via their effect on the statistics of these ensembles of snapshots. The reference frames for bases are taken from the Tsukuba convention [80], and an analogous reference frame is adopted for phosphate groups [63], as described in Figure S1 and Table S5. Additionally, base-pair frames (average of two complementary base frames) and junction frames (average of two adjacent base-pair frames) were computed to allow the definition of the full internal relative coordinates of any coarse-grain duplex configuration.

We refer to Section S2 and [63, 64, 81] for full detail on the cgNA+, or enhanced CURVES+, coordinates that we adopt. But their main features are as follows. The cgNA+ internal coordinates parametrize relative rigid-body transformations by three rotational and three translational degrees of freedom (Figure 1b). For an *N* base-pair linear blunt-ended duplex configuration (without the two dangling 5^′^ terminal phosphates), the internal coordinate vector Ω is of the form

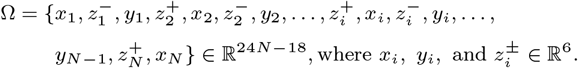

Each of the four 6-vectors 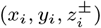 comprises three rotational and three translational degrees of freedom which are grouped into three categories (Figure 1b), each admitting a structural interpretation:

- Intra-base-pair coordinates *x*_*i*_: six standard base-pair parameters (buckle, propeller, opening, shear, stretch, and stagger) describing
- the relative position and orientation of the two bases, with the relative translations expressed in the associated base-pair frame [81];
- Inter-base-pair step coordinates *y*_*i*_: six standard base-pair-step parameters (tilt, roll, twist, shift, slide, and rise) describing the relative position and orientation of adjacent base pairs (i and i+1) with the relative translations expressed in the associated junction frame [81];
- Phosphate coordinates 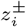: three relative translational and three rotational coordinates describing the position and orientation of the 5^′^ phosphate on the Watson (denoted by superscript +) and Crick (denoted by superscript −) strands relative to their associated bases.

The translational degrees of freedom in the cgNA+ model are expressed in Angstroms. Rotational degrees of freedom are represented using Cayley parameters (see Section S2) and are scaled to units of one-fifth of a radian (approximately 11.46 degrees). This unconventional rotational unit is adopted to ensure balanced numerical scaling between translations and rotations within the model [59, 60]. (Essentially the 1*/*5 factor arises because 5 Angstroms is a good characteristic length scale for all the rigid units and relative translations arising in dsNAs.)

#### Multivariate Gaussian approximation of dsNA in conformational space

For any dsNA sequence S, the first and second moments of the equilibrium distribution on conformational space, expressed in internal coordinates, can be approximated by a multivariate Gaussian probability density function (or pdf) of the form

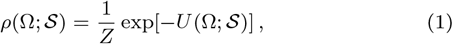

where *Z* is the partition function and *U* is a quadratic approximation to the effective free energy (in units of *k*_B_*T* with *k*_B_ Boltzmann’s constant and *T* the absolute temperature) given as

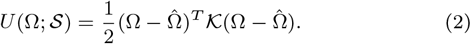

Here, the equilibrium (groundstate) configuration 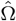and the positive-definite stiffness matrix K (or covariance matrix K^−1^) fully determine the Gaussian pdf, but their values can depend on the entire sequence S of all of the fragment, and *a priori* K need have no particular bandedness or sparsity structure.

For each sequence S_*j*_ in the training library (Tables S1-S3), an oligomer based Gaussian pdf, *ρ*_MD_(Ω; S_*j*_ ) with groundstate 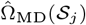and stiffness K_MD_(S_*j*_ ) for that complete sequence fragment S_*j*_ can be estimated from the ensemble of configuration coordinate snapshots generated by the MD simulations of that fragment using standard summation formulas approximating the first and centered second moments of the distribution (cf. Section S3). Those are the data from which the cgNA+ model parameters can be estimated, as described in the next section.

#### cgNA+ parameters

The first major assumption in the cgNA+ model (locality of physical interaction) is that the total model energy of a dsNA conformation (Eq. 2) is approximated as a sum of (shifted, quadratic) local dimer-step junction contributions

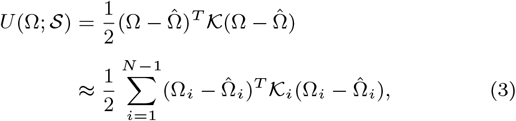

where 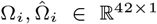and K_*i*_ ∈ ℝ^42×42^ for an interior dimer junction, and 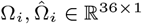and K_*i*_ ∈ ℝ^36×36^ for a terminal dimer junction (with the lower dimension at the two ends accounting for the two omitted 5^′^ phosphates). Here Ω_*i*_ denotes a local 42 (= 6 × 7) dimensional coordinate vector associated with the *i*th (interior) dimer junction, comprising two sets of intra-base-pair coordinates, one set of inter-base-pair junction coordinates, and four sets of base-to-5^′^- phosphate coordinates, each in ℝ^6^. And K_*i*_ ∈ ℝ^42×42^ is an associated localised stiffness matrix, which captures all nearest neighbour physical interactions between the eight rigid bodies associated with the four nucleotides making up the i^th^ junction (cf. Figure 1b).^1^

A crucial feature of the assumed form (3) of the cgNA+ energy is that the local coordinate vectors Ω_*i*_ are not independent. Rather Ω_*i*_ and Ω_*i*+1_ share 18 components in common corresponding to the 6 intra base pair coordinates and 12 phosphate coordinates arising at the common base-pair level shared by their two respective junction steps. The linear algebra inherent to Eq. 3 then implies that the oligomer-level stiffness matrix K(S) can be constructed by overlaying the local 42 × 42 stiffness blocks K_*i*_, to arrive at a banded symmetric matrix with overlapping blocks on the diagonal, where each overlap is 18 × 18. Similarly an oligomer-level stress-like vector *σ*(S), can be constructed by overlaying the local stress-like vectors 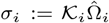. (These constructions are illustrated schematically in Figure 1f with more mathematical detail provided in Section S3.) Then the oligomer-level shape, or groundstate, configuration 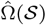 can be found from solving the banded linear system

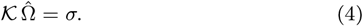

(And it is typically the case that solving the banded linear system numerically for each 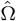is more efficient than actually computing the inverse matrix K^−1^ and carrying out the dense matrix-vector multiply on the right-hand side.)

The missing step to make cgNA+ a fully predictive model of a Gaussian pdf of the form (3) for any sequence S is a procedure that provides for each *i* the values of {K_*i*_, *σ*_*i*_} (or equivalently 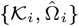. That procedure is provided by the second major assumption in the cgNA+ model, which is that these local nearest-neighbour parameter blocks in fact depend only on their local dimer-step sequence, and not on the location *i* of the dimer along the oligomer (except for the two terminal dimer steps).

A cgNA+ parameter set comprises all possible dimer-dependent stress-like vectors *σ* and local (positive definite, symmetric) stiffness matrices K

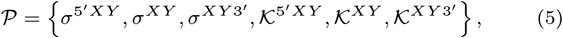

where 5^′^*XY* and *XY* 3^′^ denote, respectively, all possible terminal dimer steps, and *XY* denotes all possible interior dimer steps, with *X* and *Y* taking any possible value in the sequence alphabet appropriate for the particular dsNA. The dimensions of the terminal parameter blocks are 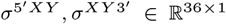 and 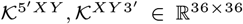 while for the interior parameter blocks *σ*^*XY*^ ∈ R^42×1^ and K^*XY*^ ∈ ℝ^42×42^.

The simplest case in which to determine the actual size of a cgNA+ parameter set is DRH in the unmodified alphabet {*A, T, C, G*} (where we adopt the convention that the sequence is read 5^′^ to 3^′^ along the DNA backbone). Then there are 16 possible parameter blocks of each type, and all blocks are independent of one another. A simple count then reveals that a single cgNA+ DRH parameter set corresponding to a given training library of MD simulations (of a given protocol) contains just under 27*K* independent scalars. Thus the cgNA+ model certainly has a Machine Learning flavour.

For dsDNA and dsRNA in the standard alphabets {*A, T/U, C, G*} the count of independent parameters is more intricate because there is the symmetry of interchanging the roles of Crick and Watson reading strands (see Section S2 for further detail). This means that all 16 5^′^ end blocks can be taken as independent, but that they then also imply all 3^′^ end blocks. For interior steps, one of the blocks for each of the six pairs of non-palindromic dimer steps AA/TT, AG/CT, AC/TG, GG/CC, GA/TC, GT/AC can be taken as independent and it then implies all the entries in its partner block, while in the four palindromic dimer steps AT, TA, CG, and GC not all entries in the blocks are independent. Thus there are fewer independent scalars in a full cgNA+ parameter set for dsDNA or dsRNA, but it is still more than 21*K*.

The parameter sets for the three different types of dsNA in their standard alphabets were all estimated independently. Parameters for methylated and hydroxymethylated C_p_G steps in dsDNA, both symmetric (both Cs modified) and asymmetric (only one C modified), were incorporated by extending the dsDNA alphabet, and corresponding parameter sets, to include 14 additional possible independent dimer steps: MN, NM, MG, AM, TM, CM, GM and HK, KH, HG, AH, TH, CH, GH, where M and H denote methylated and hydroxymethylated C, respectively, and N and K denote G paired with methylated and hydroxymethylated C, respectively. While training C_p_G modification-specific parameters, the previously estimated canonical sequence dsDNA parameter blocks were kept fixed.

#### cgNA+ parameter training

A detailed description of parameter estimation is beyond the scope of the current presentation, which is instead focused on how the model can be used given one of the existing parameter sets [65]. Implementation details for parameter estimation are described in Refs. [63, 64] and associated code is provided at https://github.com/rahul2512/cgNA_plus_training. We remark that the first step in estimating a parameter set *de novo* require computationally intensive MD simulations to construct suitable training data. Then there is a high-dimensional non-linear optimisation involving delicate computational choices (for example, the choice of initial conditions in the iterative parameter fit optimisation), but it is nevertheless rather efficient. Given the MD training data, the best fit cgNA+ parameter set P is obtained from the variational principle of minimizing over all admissible parameter sets P, the total Kullback-Leibler divergence (Eq. 8 in the next section) between the training MD-estimated and the cgNA+ predicted Gaussian pdfs over all the sequences in the training library

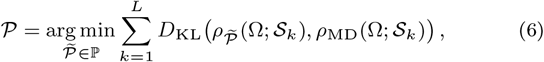

where *L* is the number of training sequences, *ρ*_MD_(Ω; S_*k*_) is the Gaussian distribution inferred from the MD simulation of sequence S_*k*_, and 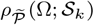 is the corresponding distribution predicted by the cgNA+ model with parameter set 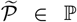. All symmetry and independence conditions on the parameter blocks defining the admissible parameter set P are explicitly enforced during the minimisation. The iterative optimisation was performed using Fisher-informed gradient descent as

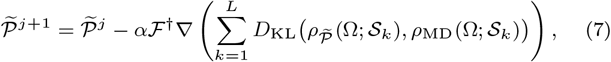

where *α* ∈ [0, 1] is step-size, *j* is iteration index, ∇ is gradient with respect to 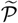, and F^†^ is pseudo-inverse of the Fisher matrix used as pre-conditioner. The above is equivalent to a quasi-Newton scheme in which the Jacobian matrix, which is precisely the Fisher information matrix in this context, is held fixed over all iterations. More details are provided in Section S3.4 and ref. [63].

#### cgNA+ predictions

In contrast to the intensive computations necessary to estimate a cgNA+ parameter set, once a specific parameter set P is in hand, the computation to use cgNA+ to predict a Gaussian equilibrium probability distribution for any dsNA sequence S of length *N* base pairs is essentially trivial. Assembling the dimer sequence-dependent local stiffness parameter blocks (cf. Figure 1f and Section S3) yields the oligomer-level stiffness matrix K(S) and stress-like vector *σ*(S), and then the groundstate configuration 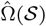 is obtained by solving the banded linear system Eq. 4.

We emphasize that a key feature in the cgNA+ model is non-local sequence dependence of the groundstate 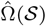. Even though the parameter set blocks are only dimer-sequence-dependent, so that the blocks in the stiffness matrix K(S) and stress-like vector *σ*(S) have only localised sequence dependence, it is nevertheless the case that the predicted groundstate configurations 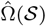 exhibit non-local sequence-dependence, which is known to occur in dsNA groundstates. Mathematically, these non-local sequence dependencies arise due to the non-local operation of back solving (or implicitly inverting the oligomer-level stiffness matrix K) in Eq. 4 to find the groundstate 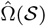. The non-locality arises because while the stiffness matrix K is block banded, it is not block diagonal, so that all the configuration coordinates are coupled one to another. The physical interpretation of the non-local sequence dependence of 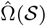 is that the two non-rigid nucleotides at each base-pair level cannot simultaneously achieve their ideal minimal energy configuration for all nearest neighbour interactions with the two nucleotides in the upstream base pair level, and at the same time achieve their ideal minimal energy configuration for all nearest neighbour interactions with the two nucleotides in the downstream base pair level. Consequently, the equilibrium structure 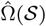of the complete molecule is determined by a global compromise along the fragment between competing local conformational preferences. The phenomenon of incompatible nearest-neighbour interactions having non-local consequences is well-known in various contexts throughout materials science, where it is generally referred to as the phenomenon of *frustration*. To the best of our knowledge its importance in the context of dsNAs was first remarked upon in [60]. Frustration is a characteristic feature of any non-rigid base pair coarse grain model of dsNAs. It cannot, however, arise with only nearest neighbour interactions in the more standard rigid-base pair coarse grain descriptions of dsNA, where non-locality in the groundstate can only be captured by allowing the model parameter sets themselves to have non-local sequence dependence. This is particularly relevant given recent work suggesting that off-site, non-local elastic couplings between a distal set of base pair coordinates play a key role in length-scale-dependent elasticity in dsNA [82].

#### cgNA+web and cgNA+ scripts

All of the steps in running cgNA+ for a particular sequence fragment are numerically sufficiently straightforward and highly efficient as to be able to make the cgNA+ model readily accessible through an interactive web implementation https://cgdnaweb.epfl.ch [65]. This interface provides a convenient way to run the model on a small number of input sequences ranging from ten or so base pairs in length (with sequences entered by hand) up to sequences of length 3K bp (which can be entered with copy-paste). The web-interface provides a selection of parameter sets, of which PS2 is recommended. All of the examples presented here use this PS2 parameter set, which is trained on the MD data generated with the protocol described above and in Section S1.

For applying cgNA+ to more than a few sequences, it is more convenient to run it in batch mode with scripts. Matlab and Python implementations are available at https://github.com/rahul2512/cgNA_plus_Matlab and https://github.com/rahul2512/cgNA_plus, with a C++ version integrated in the associated Monte Carlo code https://github.com/rahul2512/cgNA_plus_mc to be able to treat larger ensembles of longer sequences faster.

With these scripts, it is straightforward to generate ensembles of millions of Gaussians each approximating the equilibrium distribution for sequence fragments of several hundred base pairs in length (cf. Figure 10). And it is perfectly possible to scan entire genomes, for example, the *S. cerevisiae* genome treated in ref. [71]. More specifically, in this study, estimating the 510-dimensional Gaussian pdfs for each of all of the 1,048,576 decamers (embedded in 22-bp sequences) took only two hours, whereas scanning 2 million sequences of length 220 bp, followed by Monte Carlo sampling of 10K conformations per sequence and persistence-length calculations, required approximately 14K single-core CPU hours on an Intel Xeon processor.

### Quantitative comparison of two Gaussian pdfs

Both in our parameter estimation, and in our assessment of the accuracy of cgNA+ predictions, we have need of a quantitative measure of the difference between two pdfs *ρ*_1_, *ρ*_2_ ∈ ℝ^*N*^ . We will adopt one of the standard ways of doing this, namely the Kullback-Leibler (or KL) divergence [83]. In the case that both of the pdfs are Gaussian, the KL divergence takes the simple explicit form

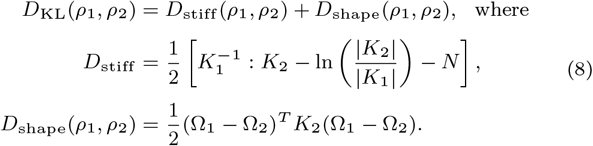

Here Ω_1_ and Ω_2_ denote the two mean vectors, *K*_1_ and *K*_2_ are the two inverse covariance (or stiffness) matrices, : denotes the standard Euclidean inner product on square matrices, and |· | is the determinant. The terms S and *D*_shape_ represent, respectively, the stiffness and shape contributions to the KL divergence *D*_KL_ with 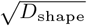 corresponding to the Mahalanobis distance [84]. Because the KL divergence and its shape and stiffness components are non-symmetric, i.e., *D*_KL_(*ρ*_1_, *ρ*_2_)/= *D*_KL_(*ρ*_2_, *ρ*_1_), we use the symmetrised form for error estimation, which is obtained by averaging over the two orderings. Throughout this article, we refer to *D*_shape_ (not 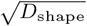) as Mahalanobis distance. Because of the presence of a stiffness matrix in the weighted inner product between the differences of the means, the units of *D*_shape_ are energy, irrespective of the different dimensions and scalings of individual components of *µ* in the specific case of cgNA+ pdfs.

There are other standard choices of divergences. We adopt KL divergence here for a variety of reasons. It has a physical interpretation as the relative entropy of one pdf with respect to the other, and in fact for bounded domains reduces to the entropy of *ρ*_1_ when *ρ*_2_ is taken as the uniform pdf. The decoupling *D*_KL_(*ρ*_1_, *ρ*_2_) = *D*_stiff_ + *D*_shape_(*ρ*_1_, *ρ*_2_) that arises for the particular case of Gaussians allows an independent discussion of error in reconstruction of mean and of stiffness. And, uniquely among standard divergences, KL has the property that in the special case where *ρ*_1_ and *ρ*_2_ are independent product measures on each component, then the KL divergence between the product measures is the sum of the KL divergences between the component measures. That apparently esoteric property means that it is sensible to scale (i.e., divide by the dimension *N* ) the KL divergence to report values per degree of freedom. In the context of cgNA+ using scaled KL divergences allows us to compare, for example, reconstruction error for a sequence of length 12 bp with reconstruction error for a sequence of length 20 bp in a meaningful way.

### Averaging over Gaussian pdfs

For certain applications discussed below, we will compare groundstate and stiffness of subsequences of a dsNA fragment within an average flanking context. To achieve this requires averaging over the Gaussian pdfs predicted by cgNA+, which can be achieved in a known, but perhaps not widely known, way (refer to chapter 6 in ref. [85]).

For a given sequence S_*j*_, cgNA+ predicts a corresponding oligomer-level Gaussian pdf, *ρ*_*j*_ (S_*j*_, P), described by its groundstate 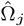 and stiffness matrix K_*j*_ . The associated covariance matrix is given by 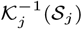, centered on 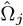 .

To compute the ensemble-averaged Gaussian pdf *ρ*_avg_ across *J* sequences S_*j*_ (*j* = 1, …, *J* ), we first evaluate the uncentered second moment matrix H_*j*_ for each sequence:

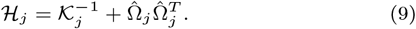

Then, the average groundstate 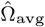 and average uncentered second moment H_avg_ across the ensemble are then given by:

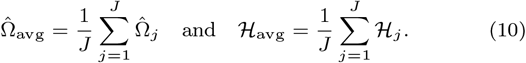

Finally, the average covariance matrix centered on 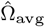 is obtained as:

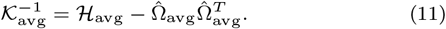

Thus, the ensemble-averaged distribution *ρ*_avg_ is fully described by the mean groundstate 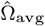 and the average stiffness matrix K_avg_.

### Configurational Volume

cgNA+ predicts a full stiffness matrix for each sequence fragment. Such a full stiffness matrix contains too much detailed information to be able to make simple comparisons between different fragments. Rather, we need a simplified proxy to be able to say that one sequence is stiffer than another. The first such proxy that we will use is configurational volume [10]. For a Gaussian pdf ∈ ℝ ^*N*^, the configurational volume is defined as

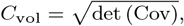

where Cov ∈ ℝ^*N* ×*N*^ is the covariance matrix. In the context of cgNA+, Cov ∈ ℝ^*N* ×*N*^ and half of the degrees of freedom are translational while the other half are rotational, therefore, *C*_vol_ has units Å^*N/*2^(rad*/*5)^*N/*2^.

In this work, we consider the configurational volume of the dimer junction. The dimer junction consists of four bases and two central phosphates, and its configuration is accordingly described by five sets of cgNA+ relative coordinates, each in ℝ^6^: two sets of intra-base-pair coordinates, two sets of base-to-phosphate coordinates (one associated with the Watson strand and the other with the Crick strand), and one set of inter-base-pair coordinates. Collectively, these coordinates define the dimer-junction fluctuations in ℝ^30^.

### Dynamic Persistence Length

The second simplified proxy for stiffness that we will adopt is sequence-dependent dynamic persistence length *l*_*d*_(S) as introduced in [66] and extensively discussed in [86] from a theoretical polymer physics point of view. The more standard and familiar tangent-tangent apparent persistence length *l*_*p*_(S) can be defined as follows. For some approximation to the unit tangent vector **t**_*i*_ at the *i*th mer along a polymer the expectations

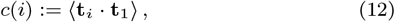

are computed, or estimated in some way, where ⟨·⟩ denotes the average over an ensemble of different configurations of the same polymer (for us, a dsNA fragment with fixed sequence S). Then *l*_*p*_ (expressed in number of mers or bp) is estimated as the characteristic length scale in the exponential fit

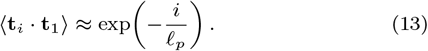

For simple polymer models, for example the standard Worm Like Chain (or WLC), it can be proven that the exponential decay assumed on the right-hand side of Eq. 13 is either a very good approximation to, or even an equality with, the data appearing on the left hand side (and for example in the WLC the length scale is simply related to the single bending stiffness constant appearing in the model). However, whenever the polymer is intrinsically curved, i.e., for us the groundstate described by 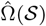is not intrinsically straight, then the exponential fit assumed in Eq. 13 is typically an extremely bad approximation to the data *c*(*i*). For dsNA, sequence examples are shown in Figure 9 panels a), b) and c), and Fig S25, where the poorness of the exponential fit is demonstrated by a lack of linearity in a semi-log plot of the data *c*(*i*) versus *i*.

In contrast, the dynamic persistence length *l*_*d*_(S) is found as the characteristic decay length scale in the exponential fit

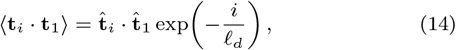

where 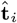 denotes the tangent vector to the groundstate configuration of the polymer. The pre-factor 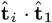 on the right-hand side (which is deterministic, there is no average) can be regarded as a non-exponential correction to the data *c*(*i*) on the left hand side. All corrected data *c*(*i*) generated within the cgNA+ model always decays rather closer to exponentially. The examples of the almost linear semi-log plots presented in panels d)-f) in Figure 9 are typical.

We report spectra of dynamic persistence lengths *l*_*d*_(S) computed in this way over ensembles of millions of sequences S of each of the three types of dsNA, both with and without epigenetic base modifications in dsDNA. Sequence-dependent dynamic persistence lengths were computed using a cgNA+mc code written in C++, which is a straightforward extension of the approach described in [66] that was originally developed within the context of the slightly simpler cgDNA model [60]. For dsNAs we take **t**_*i*_ to be the third vector of the *i*th base pair frame, which can be regarded as the unit normal to the best fit plane to the *i*th base-pair. The vectors **t**_*i*_ can be computed for each configuration coordinate snapshot Ω (Section S2), and ensembles of coordinates Ω for each sequence can be drawn directly from their equilibrium Gaussian pdf for each particular sequence S. In fact for some sequences we compute ensemble averages over two distinct Gaussian pdfs: the cgNA+ predicted Gaussian, and also the MD observed Gaussian for the same sequence. This yields another verification of the accuracy of the predictions of the cgNA+ model and its parameter sets. To minimise end effects such as fraying, terminal base pairs are excluded from the analysis. All averages were estimated from ensembles of 10^5^ Monte Carlo draws, which provides sufficiently converged statistics [63] to accurately estimate *l*_*d*_ for each S.

To avoid any confusion, and as is fully discussed in [66], we remark that the names apparent and dynamic persistence length for dsDNA were introduced some decades before the publication [66], but only in the specific context of sequence-averaged values, in which case there is a third quantity named static persistence length associated with decorrelation due to randomness in sequence. The quantities of interest here are particularly the sequence specific decorrelation data *c*(*i*), the sequence specific apparent persistence length *l*_*p*_(S), and the sequence specific dynamic persistence length *l*_*d*_(S), as introduced in [66].

### Groove widths

Groove widths of a cgNA+ configuration were computed following the method proposed in [81, 87]. For a given sequence, we chose the central phosphate on the Watson strand as the reference phosphate. A cubic spline was then fitted through the phosphate positions of the Crick strand backbone, with tangents defined by neighbouring phosphate groups. Nine equally spaced points were sampled along each spline segment and their distances to the chosen reference Watson phosphate were computed. The minor groove width was defined as the minimum distance measured in the upstream (3^′^ → 5^′^) direction along the Crick strand, whereas the major groove width was defined as the minimum distance measured in the downstream (5^′^ → 3^′^) direction. An offset of 5.8 Å was subtracted from the measured distances to account for the van der Waals radius of the phosphate groups.

## Results

### Sequence-dependent conformational landscapes

This section examines the conformational landscapes of dsDNA, dsRNA, DRH, and epigenetically modified dsDNA. We first rigorously benchmark cgNA+ predictions against atomistic MD simulations and then use the model to characterize how sequence, flanking context, and chemical modifications shape dsNA conformational landscapes.

#### cgNA+ accurately captures the non-local sequence dependence in dsNA equilibrium shape

We benchmarked cgNA+ predictions against atomistic MD simulations (using the same protocol as the training runs) for a diverse set of test sequences spanning all three dsNA classes (Tables S1–S2); none of these sequences were used during model training. Each colour plot in each panel of Figure 2 displays the groundstate values of a particular coordinate along a particular oligomer. In fact there are two line styles for same colour in each plot, solid coming from cgNA+ model predictions, and dashed coming directly from MD verification simulations. The two line styles are in many places indistinguishable, and where they can be distinguished their difference is always much, much smaller than the variation in the signal along the oligomer. These examples are absolutely typical. They, and analogous plots in the Figures S3-S5, are the justification for our claim that the accuracy of cgNA+ predictions of sequence groundstates is quite remarkable.

**Figure 2.**
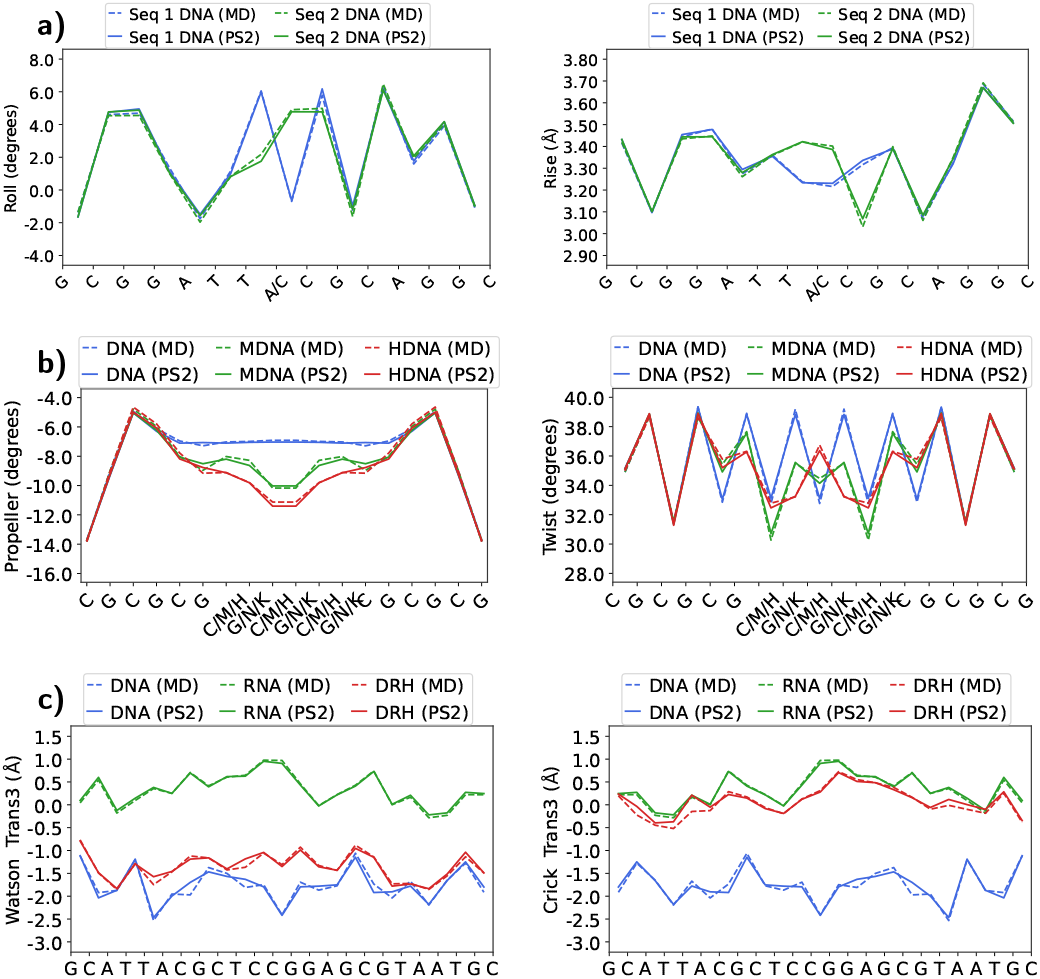
Benchmarking cgNA+ (solid line; referred to as PS2 as in the https://cgdnaweb.epfl.ch website) predictions against atomistic MD simulations (dashed line) for test sequences in predicting the equilibrium shape. **(a)** Roll and rise coordinates for two dsDNA sequences differing at only one position. **(b)** Propeller and twist coordinates for (CG)_9_ and its epigenetically methylated variant, (CG)_3_(MN)_3_(CG)_3_, and hydroxymethylated variant, (CG)_3_(HK)_3_(CG)_3_. **(c)** Third translational base-to-5^′^-phosphate coordinate on both Watson and Crick strands for the same sequence in dsDNA, dsRNA (U instead of T), and DRH versions. M and H denote the methylated C and hydroxymethylated C, while N and K denote the G paired with methylated C and hydroxymethylated C. Results are shown for a few representative coordinates, while the complete set of results for all coordinates is provided in Figures S3-S5.

The particular sequences chosen for Figure 2, were selected to illustrate various characteristic features of dsNA predicted by cgNA+ (and then verified by independent MD simulation). Figures 2a and S3 illustrate the cgNA+ accuracy in predicting the sequence-dependent equilibrium shape for two sequences that differ at only a single base-pair position. The point mutation induces a substantial non-local change in equilibrium shape that propagates several base-pairs away from the site of substitution as is evident in both the roll and rise coordinates. Figure 2b shows that cgNA+ captures the changes in propeller and twist angles upon epigenetic methylation ((CG)_3_(MN)_3_(CG)_3_) and hydroxymethylation ((CG)_3_(HK)_3_(CG)_3_) of poly(CG); see Figure S4 for the full set of coordinates. Figure 2c shows the cgNA+ accuracy in predicting the phosphate coordinates for dsDNA, dsRNA (U instead of T), and DRH version of a test sequence; see Figure S5 for the full set of coordinates. The phosphate coordinates on the Watson strand (DNA strand) are closer to the dsDNA coordinates, whereas the phosphate coordinates on the Crick strand (RNA strand) are closer to the dsRNA coordinates. This asymmetry reflects the hybrid chemical nature of DRH, in which the two backbones retain distinct structural signatures in the equilibrium shape.

To quantify prediction accuracy, we define the Sequence-Variability, or SV, as the mean pairwise symmetric Mahalanobis distance (Eq. 8) per degree of freedom between the MD-inferred Gaussian distributions of training sequences. This number then reflects a characteristic variability attributable to the sequence. The SV values are 0.0223 (dsDNA), 0.0177 (dsRNA), and 0.0209 (DRH). We then compared this SV with the prediction error, defined similarly as the symmetric Mahalanobis distance per degree of freedom between Gaussian pdfs inferred from MD and cgNA+. The average prediction error, computed over 28 dsDNA (including C_p_G modified sequences) test sequences, 8 dsRNA test sequences, and 7 DRH test sequences, is approximately one order of magnitude smaller than the corresponding SV: 0.0026 for dsDNA, 0.0015 for dsRNA, and 0.0023 for DRH (Table S6). Furthermore, prediction errors for training sequences are of comparable magnitude, confirming that the model generalises well without overfitting (Table S6).

#### Average shape of dsDNA, dsRNA, and DNA:RNA hybrid dimers in all tetramer contexts

Here we present the variation in equilibrium shape of the central dimer junction across all tetramer flanking contexts in dsDNA (including C_p_G modifications), dsRNA, and DRH, as observed in MD simulations of the training libraries, and as predicted by cgNA+. The dimer junction comprises four bases and two central phosphates, described by five sets of cgNA+ relative coordinates in R^6^: two sets of intra-base-pair coordinates, two sets of base-to-phosphate coordinates (one on the Watson strand and the other on the Crick strand), and one set of inter-base-pair coordinates. Figure 3 focuses on selected coordinates of a representative dimer junction from the purine–purine, purine– pyrimidine, pyrimidine–purine, pyrimidine–pyrimidine, and modified C_p_G classes. The complete set of results is provided in Figures S6–S15.

**Figure 3.**
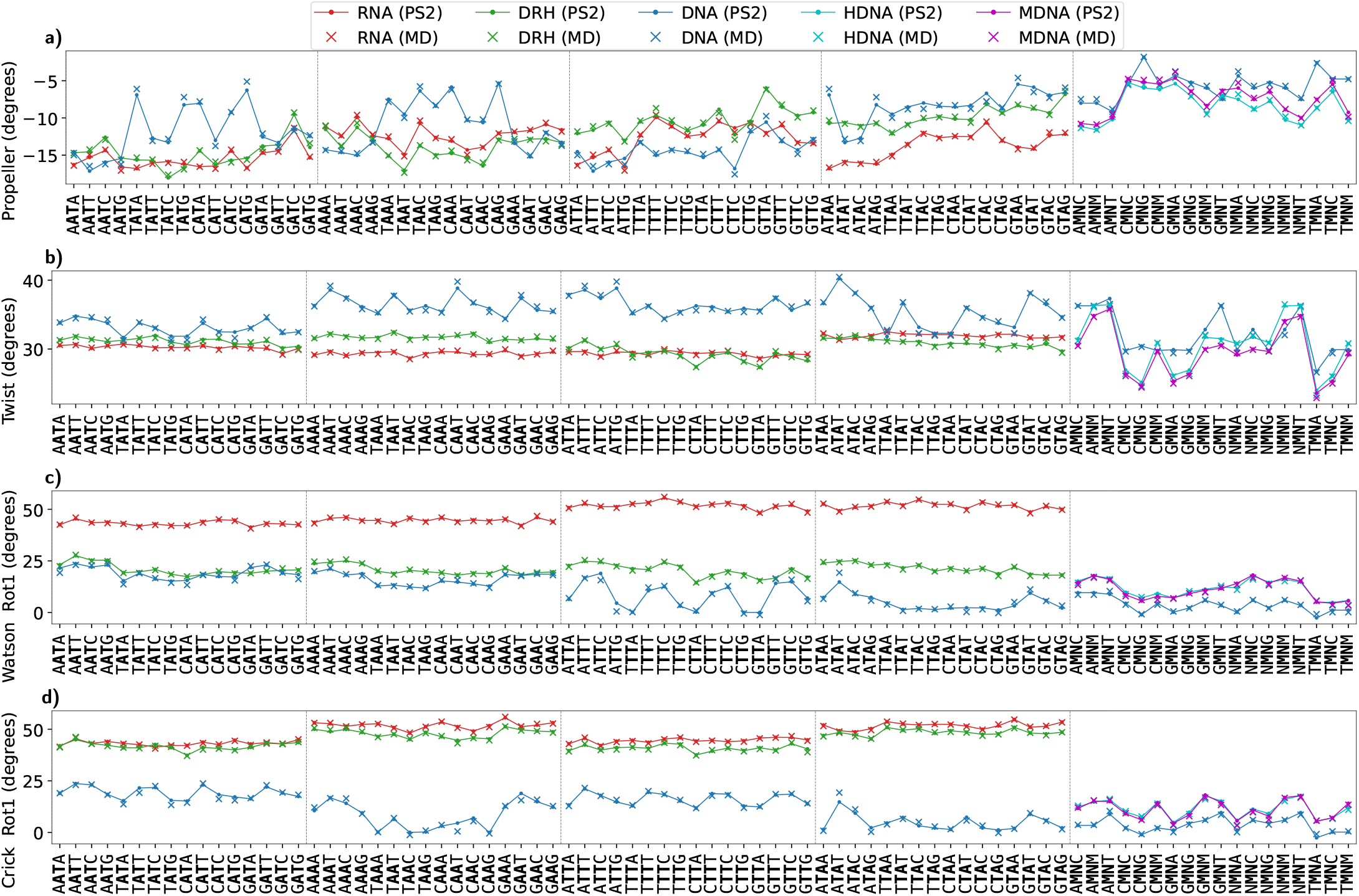
Comparison of representative base-pair, base-pair-step, and phosphate coordinates for selected central dimer steps in all tetramer flanking contexts across dsNA. **(a-b)** Propeller (intra base-pair) corresponding to the first base-pair in the central dimer and Twist (inter base-pair step). **(c-d)** First rotational base-to-5^′^-phosphate coordinate on both Watson and Crick strands across dsDNA, dsRNA, and DRH. The results shown here correspond to one representative central base-pair step from each sequence class–AT (purine–pyrimidine), AA (purine–purine), TT (pyrimidine–pyrimidine), and TA (pyrimidine–purine)–compared across dsDNA, dsRNA, and DRH, in addition to the C_p_G step in all independent tetramer contexts along with its methylated and hydroxymethylated version. M denotes the modified C and N the guanine paired modified C. Their specific identities follow the legend: in unmodified DNA, M = C and N = G; in methylated DNA (MDNA), M denotes methylated C and N denotes G paired with methylated C; and in hydroxymethylated DNA (HDNA), M denotes hydroxymethylated C and N denotes G paired with hydroxymethylated C. The results for all dimers and coordinates are provided in Figures S6–S15. Note that the statistics were derived from the training library; consequently, the sequence context beyond the central tetramer is not averaged but corresponds to specific flanking sequences. Coordinates observed in MD simulations and cgNA+ predictions are plotted as × and •, respectively. For better visualization, a line plot is overlaid along the • symbols.

Figure 3a shows that propeller values (corresponding to the first base pair of the central dimer) are generally more negative in dsRNA than in dsDNA, with DRH typically adopting intermediate values. An exception is the TT step (pyrimidine–pyrimidine), for which propeller in dsDNA is more negative than, or comparable to, that in dsRNA. For the C_p_G step, the propeller value at the cytosine base pair is closer to zero, likely reflecting the greater rigidity conferred by the three hydrogen bonds of the C:G pair relative to the two bonds of A:T. This is a general trend for intra-base-pair coordinates: their equilibrium shape and fluctuations are strongly governed by the hydrogen-bonding pattern (Figures S6, S10, S11, and S15). Both methylation and hydroxymethylation of the C_p_G step reduce propeller further toward more negative values. Across all dsNA systems, the spread in propeller values arising from flanking context is often larger than the variation between different base-pair types, underscoring that flanking context is as informative as base-pair identity for this coordinate. Note that the cgNA+ predictions are nearly indistinguishable from the MD observations throughout.

Figure 3b plots an inter-base-pair coordinate, twist, which follows the ordering dsDNA *>* DRH *>* dsRNA across the majority of dimer steps. For the C_p_G step, both methylation and hydroxymethylation decrease twist, with the reduction being slightly larger for hydroxymethylation. The higher average twist in dsDNA relative to dsRNA reflects their characteristic B-form and A-form geometries, respectively, while DRH, with its predominantly A-form character, generally exhibits twist values closer to those of dsRNA. Noy et al. [43] similarly reported that twist and slide in DRH are closer to dsRNA values, while the remaining inter-base-pair step coordinates tend to lie between the dsDNA and dsRNA values. More generally, for inter-base-pair translational coordinates, dsDNA dimers are substantially more sensitive to flanking tetramer context than dsRNA, with DRH again showing intermediate behaviour (Figure S8). The same qualitative trend is observed for the rotational coordinates, although the differences are less pronounced.

The phosphate coordinates in dsDNA and dsRNA are strikingly different, a direct consequence of their B-form and A-form helical geometries, respectively (Figures S7, S9). Furthermore, in contrast to inter base-pair coordinates, the observed phosphate coordinates in DRH are not in between dsDNA and dsRNA; instead, Watson phosphate coordinates are closer to those observed in dsDNA, whereas Crick phosphates are closer to those observed in dsRNA. This implies that the backbone behaviour of the DNA strand in DRH is closer to pure dsDNA, and the RNA strand is closer to pure dsRNA. Similar findings have been reported in [43, 44, 88, 89] but are characterized in different variables. More specifically, in Figure 3c–d, the Watson phosphate is attached to the first base pair of the central dimer step, whereas the Crick phosphate is attached to the second base pair. This explains the difference observed in phosphate Rot1 between the AT/AA and TT/TA steps. Furthermore, epigenetic modification of the C_p_G step has a considerable effect on phosphate Rot1, although the difference between methylation and hydroxymethylation is negligible.

Notably, dimer-step-dependent fluctuations in phosphate coordinates for dsDNA are considerably higher than those in dsRNA (Figures 3c-d, S7, S9). Moreover, for a given dimer step, the differences in average phosphate coordinates for various flanking tetramer contexts are also much higher in dsDNA than in dsRNA. This trend is consistent with the broader conformational flexibility of the dsDNA backbone, which can sample a wider range of geometries, including sequence-dependent shifts between B-form and A-form structures, whereas dsRNA remains predominantly constrained to A-form geometry. In the case of DRH, once again, one can observe that the RNA strand behaves similarly to pure dsRNA (with less variation due to the dimer step sequence and the flanking tetramer context), and the DNA strand behaves similarly to pure dsDNA (i.e., sensitive to dimer step sequence and flanking tetramer context).

We note that pairwise comparisons of dimer-step geometry across dsDNA, dsRNA, and DRH or C_p_G modification in dsDNA have been reported previously, often for a limited number of sequences [19, 20, 43–45, 88, 89]. The results presented here highlight the sequence-dependent variability in dimer equilibrium shape induced by tetramer flanking context across all duplex classes and dimer equilibrium shape is a function of both its identity and its local sequence context. More importantly, cgNA+ captures this non-local sequence-dependence in equilibrium shape using only local dimer dependent parameter blocks across dsNA classes.

#### Dimer-step flexibility drives flanking-context sensitivity in its equilibrium shape

Having established that cgNA+ accurately captures the MD-inferred conformational landscapes for independent test sequences across dsNA classes and epigenetic states, we next use the model to extend the analysis to regimes that would be impractical to study systematically by atomistic simulation.

In the previous subsection, we observed that the variability in structural parameters arising from tetramer flanking context is often larger than the variability associated with the central dimer itself. For each dimer-step, we quantified this by computing the symmetric Mahalanobis distance per degree of freedom (Eq. 8) between its equilibrium shape in a given octamer context and that in the average context (see Materials and Methods). Figure 4a plots this statistic for each central dimer step, showing that the equilibrium shape of a dsDNA dimer depends strongly on its flanking sequence context. This dependence is strongly dimer-step-specific: pyrimidine–purine steps are generally the most context-sensitive, with TA being the most context-sensitive and AT the least. A similar analysis by Battistini et al. [5] quantified context effects in hexamer space via the deformation energy required to transfer a dimer between equilibrium shapes in different hexamer contexts.

**Figure 4.**
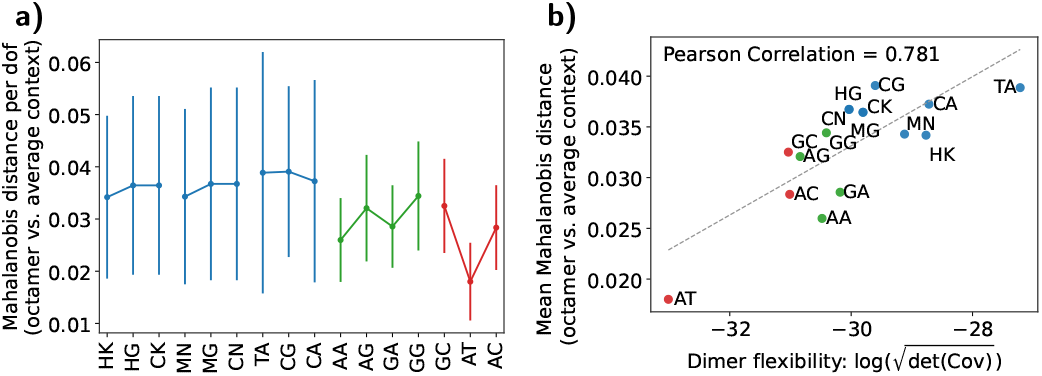
**(a)** Symmetric Mahalanobis distance between dimer’s equilibrium shape in a specific octamer context and its equilibrium shape in the average context. The bars denote the standard deviation across all octamer contexts. **(b)** Pearson correlation between the mean Mahalanobis distance of the dimer equilibrium shape in a specific octamer context relative to the average context and the logarithm of the configurational volume of the dimer in the average context. For fluctuations in ℝ^*N*^, the configurational volume is defined as 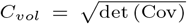 [10] (see Materials and Methods). Here the dimer junction fluctuations are in R^30^ and the unit of *C*_*vol*_ is Å^15^· (rad*/*5)^15^. Dimer level configurational volume and analogous correlation plots for dsRNA and DRH are provided in Figure S16.

One possible physical interpretation is that a mechanically soft dimer step can be more easily deformed by its neighbouring bases, whereas rigid steps are less easily modulated by the flanking sequence environment. To examine this hypothesis, we quantified the flexibility of each dimer step in the average context through its configurational volume, defined as 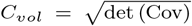 where for physical dimer steps, or bp junctions, the covariance matrix Cov ∈ ℝ^30×30^ [10] (see Materials and Methods). Specifically, for each central dimer, we computed the configurational volume averaged over every octameric sequence context embedded within fixed terminal GpC ends. The mean context-dependent deviation in equilibrium shape is positively correlated with the logarithm of this configurational volume (Pearson correlation = 0.78), supporting this interpretation (Figure 4b). Thus, dimers with larger configurational volume, and hence greater intrinsic flexibility, also tend to exhibit stronger modulation of their equilibrium shape by the surrounding sequence environment.

Similar trends are observed for the equilibrium shapes of dsRNA dimers, although their sensitivity to flanking sequence context is lower than in dsDNA, consistent with the greater stiffness of dsRNA dimers (Figure S16). Within dsRNA, pyrimidine–purine steps are again the most sensitive to flanking context, in line with their comparatively high flexibility. By contrast, DRHs exhibit a more complex mixed pattern: pyrimidine–pyrimidine steps are less sensitive to flanking context, yet at the same time are among the most flexible in terms of configurational volume (Figure S16).

As a remark, the variation in equilibrium shape induced by changes in flanking sequence, measured as the Mahalanobis distance per degree of freedom, provides an alternative SV for assessing *cgNA*+ accuracy. By this measure, the context-induced variation (0.0304 ± 0.0159) is again approximately one order of magnitude larger than the *cgNA*+ prediction error (average Mahalanobis distance per degree of freedom for 28 dsDNA test sequences: 0.0026).

#### CpG modification alters the equilibrium shape comparable to single-nucleotide polymorphisms

Single nucleotide polymorphisms (SNPs) are the most common genetic variations in the human genome. Out of 12 possible SNPs, only six are independent (A↔T, A↔C, A↔G, T↔C, T↔G, and G↔C). Furthermore, in a systematic study of all flanking contexts, A↔G and T↔C are dependent; similarly, A↔C and T↔G, thus leaving only four independent SNPs. We quantified SNP influence on dsDNA mechanics by computing the change in groundstate, in terms of symmetric Mahalanobis distance (Eq. 8), for each independent SNP embedded in all tetramers on both sides plus a fixed flanking sequence to avoid end-effects.

Figure 5a shows the change in groundstate associated with each SNP, with bars indicating the influence of the flanking sequence context. As expected, substitutions within the purine class induce smaller changes in the groundstate than substitutions between purines and pyrimidines. The A↔G substitution produces the smallest change in groundstate, whereas A↔T produces the largest change, approximately twice that observed for A↔G. However, the relative ordering among purine↔pyrimidine SNPs, A↔T *>* A↔C *>* G↔C, is less straightforward to interpret.

**Figure 5.**
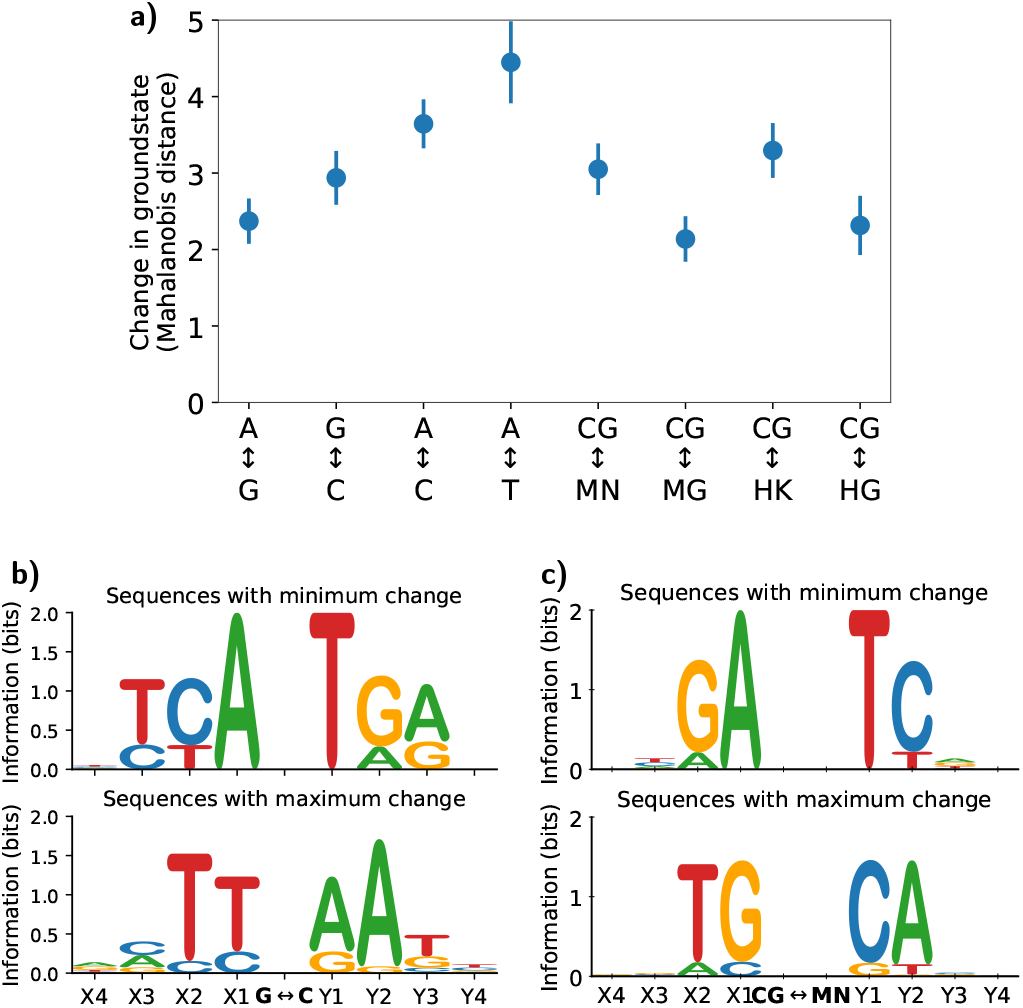
**(a)** Change in groundstate in terms of symmetric Mahalanobis distance due to single nucleotide polymorphisms (SNPs) at the central base-pair and epigenetic modifications at the central C_p_G steps, with bars indicating the influence of the flanking context. Statistics are derived from all tetramer flanking contexts on both sides of the central SNP position or central C_p_G steps embedded in a 21/22mer: GCGTCGX_4_X_3_X_2_X_1_—central position — Y_1_Y_2_Y_3_Y_4_GTCGGC. Sequence logos [90] for flanking contexts associated with the smallest and largest changes (beyond three standard deviations from the mean) in groundstate upon G↔C substitution at the 5^th^ (central) position **(b)** and epigenetic modification of the central C_p_G step **(c)**. In sequence logos [90], the x-axis is the base index of the sequences, and the y-axis is information content with the maximum possible value of two for dsNA. Analogous sequence logos for other SNPs and C_p_G modifications are provided in Figure S17.

Furthermore, the flanking sequence strongly influences the change in groundstate upon point mutation, as indicated by the bars in Figure 5a. Figures 5b and S17a-c show sequence logos [90] for flanking contexts in which a given point mutation produces the smallest or largest change in groundstate obtained from outlier sequences (three standard deviations from the mean). Notably, in addition to the immediate flanking context, the next- and next-nearest-neighbour contexts also play an important role. To visualize the effect of a point mutation on the groundstate of a given sequence, we plotted the groundstates of two sequences in Figure S18; one shows a slight change in groundstate upon A↔G substitution, whereas the other shows a highly non-local change in groundstate upon A↔T substitution. The non-local change is up to four base pairs on either side of the A↔T substitution.

Furthermore, Figure 5a compares SNP-induced changes in groundstate with those caused by epigenetic modification of the central C_p_G step. The change in groundstate for asymmetric methylation or hydroxymethylation is comparable to that for A↔G SNPs, whereas symmetric C_p_G modifications produce larger changes comparable to G↔C SNPs.

The effect of the flanking sequence context is similar to that observed for SNPs. Sequence logos for the sequences exhibiting the smallest and largest changes in equilibrium shape upon symmetric C_p_G methylation (Figure 5c) show that the least affected sequences are characterised by a GA step upstream of C/M, whereas the most affected sequences are characterised by a TG step upstream of C/M, with negligible additional information beyond the hexamer context. Analogous analysis (Figure S17d-f) for asymmetric methylation (C_p_G to M_p_G) and hydroxymethylation (C_p_G to H_p_G/H_p_K) also leads to similar conclusions. Figure S19 compares the groundstate for two sequences that only differ in the immediate flanking context (underlined) to C_p_G step, GCGTCGGTGA***CG***TCTTGTCGGC and GCGTCGGTTG-***CG***CATTGTCGGC to visualize the change in groundstate on the symmetric methylation of the central C_p_G step (in bold). The change in groundstate for the latter sequence on C_p_G methylation is much larger and non-local compared to the former, particularly for the phosphate coordinates.

Epigenetic modifications are traditionally thought to silence genes primarily by increasing dsDNA stiffness [91, 92]. Here, we show that they also significantly alter the dsDNA groundstate. More broadly, both SNPs and epigenetic modifications can induce substantial changes in equilibrium shape, with effects that are often highly non-local, extending several base pairs away from the site of chemical change. Together, these findings highlight that both genetic and epigenetic variation can have significant, context-dependent structural consequences for dsDNA structure.

#### Comparison of groove widths of dsDNA, dsRNA, and DNA:RNA hybrid

Protein–DNA recognition is strongly influenced by sequence-dependent DNA groove geometry, as observed in both experiments and simulations [17, 87, 93]. Groove geometry is therefore a particularly relevant structural feature to characterise within the cgNA+ framework.

Figure 6a plots the distance between the central Watson phosphate and a spline interpolating the Crick phosphate positions, computed from cgNA+ and MD groundstate configurations for a representative dsDNA, dsRNA, and DRH test sequence. The cgNA+ and MD profiles for all three dsNA sequences are nearly indistinguishable, demonstrating the accuracy of cgNA+ in predicting the backbone geometry relevant to groove width calculations. The distance profiles also capture the distinct geometric features of dsDNA, dsRNA, and DRH: dsDNA is typically B-form, with a major groove wider than the minor groove; dsRNA is typically A-form, with a comparatively narrow major groove and a wider minor groove; and DRH exhibits intermediate groove widths between dsDNA and dsRNA.

**Figure 6.**
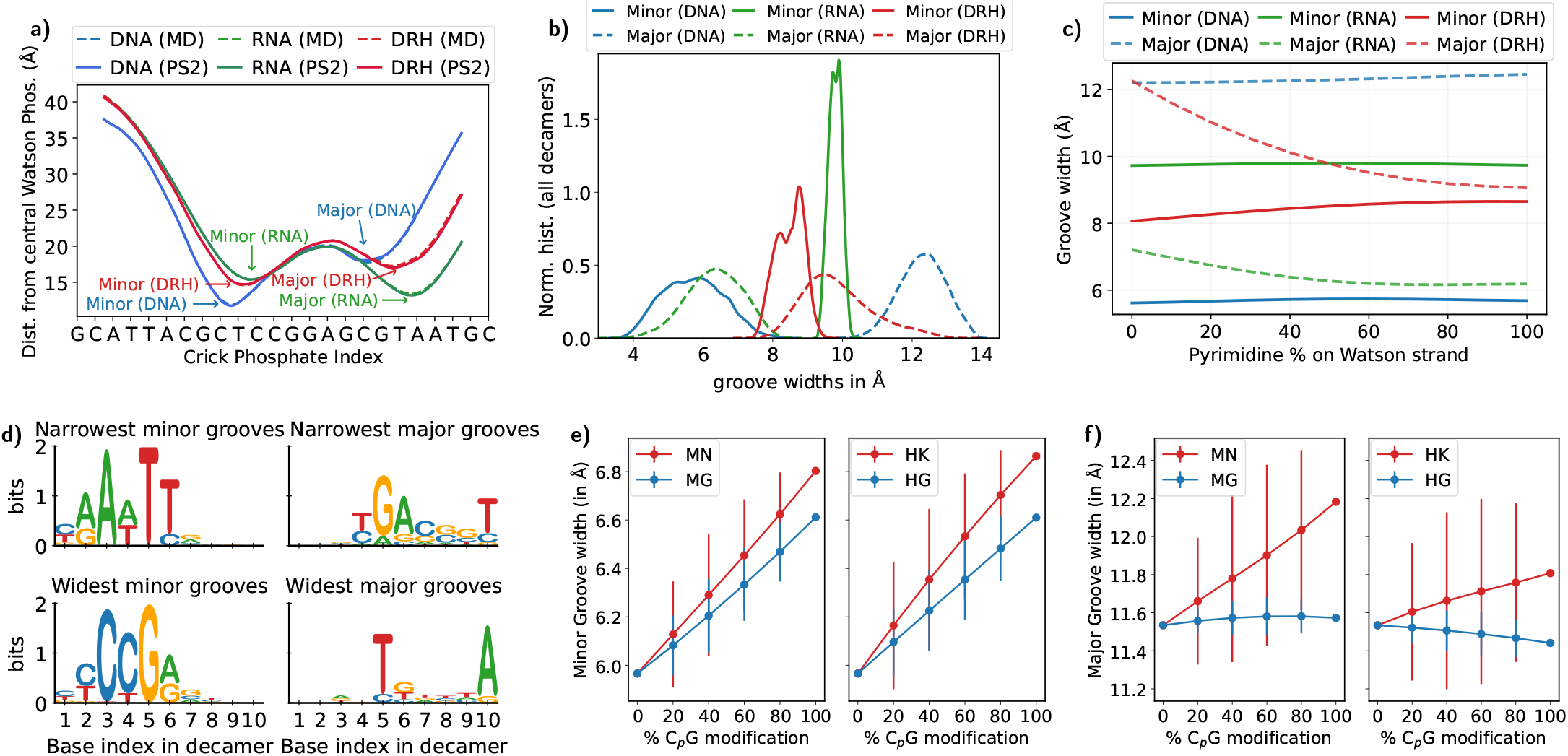
Distribution of major and minor groove widths across dsNA. **(a)** Distance between the central Watson phosphate position and a cubic spline interpolating the Crick phosphate positions, computed from the cgNA+ predicted (referred to as PS2) and the MD groundstate for a test sequence shown in its dsDNA, dsRNA (U in place of T), and DRH versions. **(b)** Histograms for major and minor groove widths in dsDNA, dsRNA, and DRH. The groove widths are computed for all decamers (approximately one million sequences) embedded in fixed flanking contexts. The Watson phosphate between 5^th^ and 6^th^ base-pair of decamer is taken as the reference to compute the groove widths. **(c)** Dependence of major and minor groove widths on Watson strand pyrimidine content (C or T) for dsDNA, dsRNA, and DRH. Detailed results are provided in Figure S20. **(d)** Sequence logos for sequences with outlier major and minor groove widths (beyond three standard deviations from the mean) in dsDNA. Corresponding plots for dsRNA and DRH are provided in Figure S21. In sequence logos [90], the x-axis is the base index of the sequences, and the y-axis is information content with the maximum possible value of two for dsNA. The total height of the stack of A/T/C/G alphabets tells the information present in that index, and the relative height of the alphabets represents their frequency at that index. Change in minor **(e)** and major **(f)** groove widths due to extent of C_p_G modifications in the highlighted sub-sequence of GCTGTG**CGCGCGCGCG**CATGGC.

Figure 6b shows the minor and major groove width spectra for dsDNA, dsRNA, and DRH computed from all decamers (approximately one million sequences, each embedded in fixed six base-pair flanking sequence on both sides). For dsDNA, the minor groove width (5.72 ± 0.86 Å) is about 6.5 Å narrower than major groove width (12.29 ± 0.66 Å) and the major groove width varies less than the minor groove width across sequence space. For dsRNA, the minor groove width (9.79 ± 0.19 Å) is about 3.5 Å wider than the major groove width (6.32 ± 0.80 Å) and major groove width varies more than the minor groove width across sequence space. Lastly, DRH exhibits intermediate behaviour in both groove dimensions, with minor and major groove widths of 8.50 ± 0.40 Å and 9.87 ± 1.01 Å, respectively, and a less pronounced separation between the two. These trends are consistent with previous observations from experiments [17, 93] and simulations [43, 44, 46], which were, however, limited to a few sequences.

Figures 6c and S20 show how groove widths depend on pyrimidine content (on Watson strand) for dsDNA, dsRNA, and DRH. Both canonical duplexes showed limited dependence on Watson strand pyrimidine content: dsDNA varied only weakly, while in dsRNA the minor groove remained nearly unchanged and the major groove narrowed modestly from approximately 7.2 Å to 6.2 Å. In contrast, DRH exhibited pronounced compositional sensitivity, with the minor groove widening from approximately 8.1 Å to 8.7 Å and the major groove narrowing from 12.3 Å to 9.1 Å as Watson strand pyrimidine content increased. The largest changes occurred between 0% and 60% pyrimidine, after which both groove dimensions approached a plateau. These findings demonstrate that DRH groove widths are substantially more sensitive to sequence composition than canonical DNA or RNA duplexes. A related MD study reported similar trends in minor groove width, together with backbone structural analysis showing that DRH adopts a mixed A/B-form conformation, with increasing pyrimidine content in the DNA strand shifting the structure towards a more A-like geometry [94].

Figure 6b shows that groove widths depend strongly on sequence; for example, the dsDNA minor groove width spans a range of approximately six Å. To better understand the sequence determinants of groove widths, we examined the sequences associated with extreme groove widths (outside three standard deviations from the mean) by plotting sequence logos [90] for the central decamer. For minor grooves, the information is only present at positions 2 to 6 of the decamer as the minor groove is upstream of the reference phosphate (between indices 5 and 6). Figure 6(d) shows that A/T-rich sequences are associated with narrow minor grooves, whereas C/G-rich sequences are associated with wider minor grooves. The general trends observed here are qualitatively consistent with available crystallographic and NMR data [17, 93]. Moreover, we observed that most sequences with narrow minor grooves are A-tracts and surprisingly do not have a single TA step (in position indices 2 to 6 of the decamer). A similar conclusion has been reached in protein-DNA X-ray crystal data where TA steps are found to be correlated with minor groove widening [17]. Furthermore, the major groove is narrower when positions 5 and 10 are occupied by G and T, respectively, and wider when these positions are occupied by T and A.

Analogous analysis for dsRNA (Figure S21) shows that narrower major grooves correlate with A/U-rich sequences, whereas wider major grooves correlate with C/G-rich sequences. Similar observations have been made from MD simulations for six dsRNA sequences [46] with poly(AU) and poly(CG) exhibiting the narrowest and widest major grooves. The dsRNA minor groove width varies over a restricted range of less than 2 Å, with G/C-rich sequences favouring narrower grooves and A/U-rich sequences favouring wider grooves.

Finally, sequence logos for DRH in Figure S21 show that A/G-rich sequences prefer narrower minor grooves and wider major grooves, whereas C/T-rich sequences (less clear signal) prefer wider minor grooves and narrower major grooves. Note, RNase H recognition of DRH has been linked to an intermediate minor groove width, which is thought to facilitate selective degradation of the RNA strand [28, 29].

At the same time, the strong sequence dependence of DRH groove widths, evident in Figures 6b and S21, suggests that local groove geometry may contribute importantly to RNase H recognition. This possibility is supported by prior studies showing reduced RNase H activity upon minor groove widening as pyrimidine content in the DNA strand increases [29, 94].

Although C_p_G methylation is generally thought to narrow minor grooves and widen major grooves [97, 98], structural analyses revealed groove width depends on sequence context and modification position [99]. Figure 6e shows a positive correlation between the minor groove widths and the extent of C_p_G modifications on the central decamer of GCTGTG(CG)_5_CATGGC. A fully symmetrically methylated or hydroxymethylated sequence (C_p_G to M_p_N/H_p_K) has a minor groove roughly 1 Å wider than its unmodified version. Moreover, the widening of the minor groove upon C_p_G modification is greater for symmetric modifications than for asymmetric ones, and hydroxymethylation of C_p_G step widens the minor groove more than methylation. The bars highlight the importance of the modified C_p_G step’s position, indicating that sequences with the same modification percentage can exhibit different minor groove widths. Lastly, C_p_G modification slightly widens the major grooves (Figure 6f). It must also be noted that along with structural changes, C_p_G modification significantly changes the chemical environment inside the grooves (with the methyl group being hydrophobic while hydroxymethyl is hydrophilic) and therefore, has implications in protein-DNA interactions [99].

#### CTCF binding sites exhibit distinctive DNA groove architecture

Next, we investigated the groove geometry of 35,765 CTCF binding sites identified using FIMO [95] with a *p*-value threshold of 10^−6^. For this analysis, we considered a 119-bp window centred on each predicted 19 bp CTCF binding site. We also generated a genomic control set by sampling one random 119-bp sequence from the human genome for each CTCF site, matched individually to the same GC content. This produced a GC-matched control set with the same number of sequences as the CTCF dataset. Groove widths at each dimer step, for both CTCF motifs and genomic controls, were computed using dimer’s central Watson-strand phosphate as the reference.

Figures 7a-b show the distributions of major and minor groove widths across the central 19-bp CTCF binding sites. Both differ significantly from those of the GC-matched genomic controls at most positions (Mann–Whitney U test, *p*-value *<* 10^−50^). To assess whether these differences extend beyond the motif itself, Figure 7c shows the difference in groove width between CTCF sites and the GC-matched controls across the 119-bp window. The results reveal a localized groove width signature associated with the 19-bp CTCF motif, with deviations of up to approximately 0.5 Å relative to the genomic control. These deviations are confined to the motif region and are not observed in the flanking sequences. Together, these results reveal that CTCF-binding sequences preponderantly display a distinctive local groove geometry, although orthogonal validation will be required to establish whether groove geometry directly contributes to CTCF recognition.

**Figure 7.**
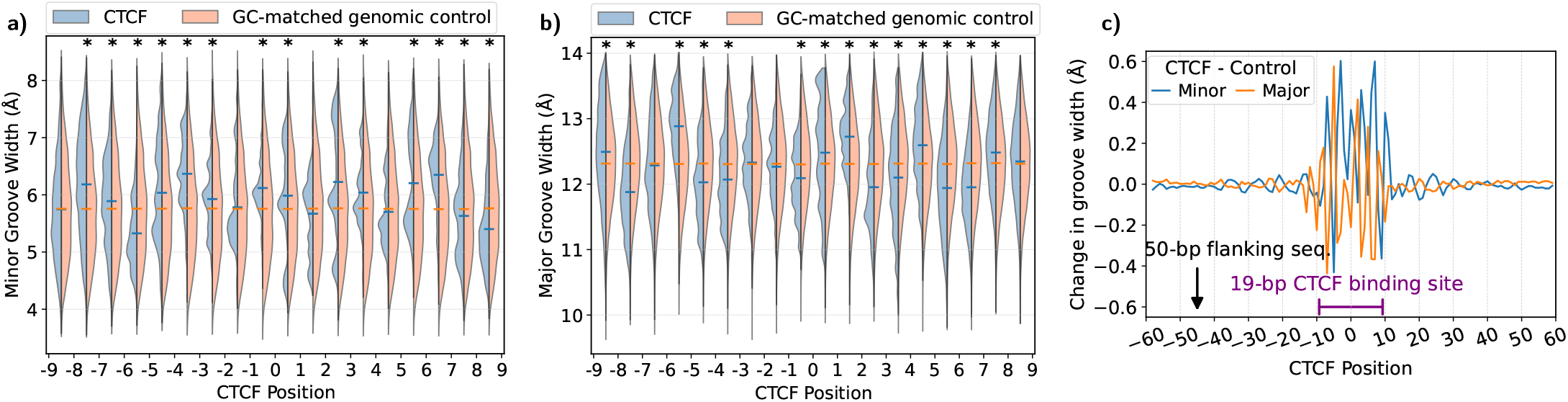
Groove geometry of predicted CTCF binding sites. CTCF motifs were identified using FIMO [95] with a *p*-value threshold of 10^−6^, yielding 35,765 sites of length 19 bp. For each CTCF site, a 119-bp window centred on the 19 bp motif was analysed, together with a GC-matched genomic control sequence of the same length. Groove widths at each dimer step were computed relative to the central Watson-strand phosphate. Violin plots [96] compare the distributions of major **(a)** and minor **(b)** groove widths at motif positions between CTCF sites and GC-matched controls. The left and right lobes represent CTCF sites and GC-matched controls, respectively. Asterisks mark positions with significant differences by a two-sided Mann–Whitney U test (*p*-value *<* 10^−50^). **(c)** Mean groove width differences between CTCF sites and GC-matched controls across the 119-bp window show a localized structural signature centred on the 19 bp CTCF binding motif.

### Sequence-dependent mechanical landscapes

We next examine sequence-dependent mechanical properties beyond groundstate configuration predicted by cgNA+, including local stiffness matrices and long-range bending behaviour. We first benchmark cgNA+ against estimates from atomistic MD simulations, and then leverage its computational efficiency to investigate how sequence composition and epigenetic modifications influence dsNA flexibility, enabling statistical analysis across sequence space.

#### cgNA+ accurately captures second moments

Figure 8 compares the stiffness matrix for a representative dsDNA test sequence obtained directly from an ensemble of MD simulation snapshots against the one predicted by cgNA+, together with their element-wise differences. Analogous plots for dsRNA and DRH stiffness are shown in Figures S22-S23. The MD stiffness matrix exhibits a banded diagonal structure, with only a few small entries beyond nearest-neighbour interactions (highlighted by the green stencils), thereby providing strong support for the nearest-neighbour assumption underlying the cgNA+ model. Visually, the cgNA+ predictions are very close to the MD reference, with small discrepancies limited to a few matrix elements. Quantitatively, the agreement is assessed using the symmetrized Kullback–Leibler (KL) divergence defined in Eq. 8. The average symmetrized KL divergence values over test libraries (Tables S1, S2, S6) are 0.0312 for dsDNA including C_p_G-modified sequences (SV: 0.3806), 0.0085 for dsRNA (SV: 0.2185), and 0.0262 for DRH (SV: 0.3273). SV is defined analogously to the groundstate, as the average pairwise KL divergence per degree of freedom between MD Gaussian estimates across all training sequences. These errors are one order smaller than the corresponding SV, demonstrating that cgNA+ accurately reproduces the sequence-dependent stiffness matrices across dsNA.

**Figure 8.**
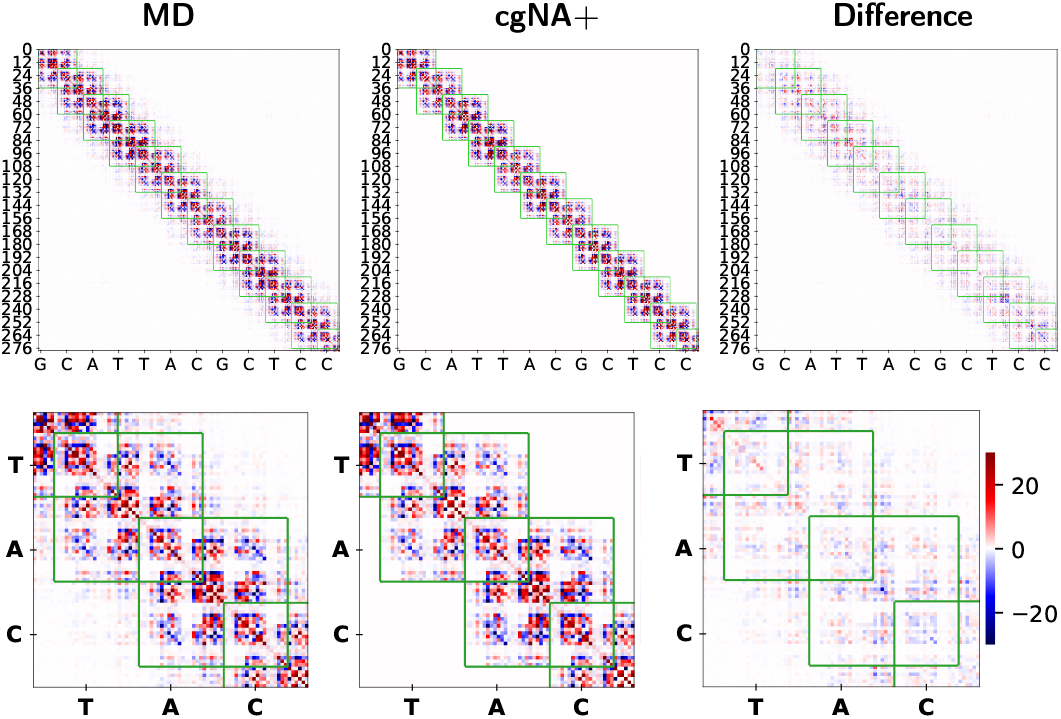
Comparison of stiffness matrices for a test sequence GCATTACGCTCCGGAGCGTAATGC (dsDNA). Top panel: the stiffness matrix estimated from MD simulations is shown on the left, the cgNA+ prediction in the center, and the element-wise difference between these two matrices on the right. Because the sequence is palindromic, only half of each matrix contains independent entries; accordingly, only this portion is displayed. The green outlines indicate the (42×42) blocks corresponding to the nearest-neighbour nucleotide interactions, so that for the cgNA+ predictions all entries outside the stencil vanish. For MD data, some entries outside the stencil are non-zero, but they are pointwise small in absolute value when compared with the large entries inside the stencil. The lower panel shows magnified views of the corresponding matrices. All matrices in the figure share a common color bar spanning -30 to 30. The discrepancy between the MD-derived and cgNA+ matrices is quantified by the symmetrized KL divergence (Eq. 8) per dof, which is 0.0338 for this test sequence. Analogous plots for the dsRNA and DRH are presented in Figures S22–S23.

As an independent validation of duplex stiffness, we examined the tangent–tangent correlation (Eq. 12), a classical measure of nucleic-acid flexibility. Figure 9a compares the logarithm of the tangent–tangent correlation for a test sequence in dsDNA, dsRNA, and DRH, as computed from samples drawn from the Gaussian pdfs inferred from MD simulations and cgNA+ predictions. The dsDNA exhibits a gradual, monotonic decay, whereas the dsRNA and DRH display oscillations arising from their intrinsic A-form helical geometry, which has a substantially wider diameter than the B-form geometry of DNA. Figure S24 shows top views of the 3D equilibrium shapes for the same sequences, illustrating the contrasting helical diameters in the order dsRNA *>* DRH *>* dsDNA. Note that the uneven oscillation amplitudes along the sequence in the tangent–tangent correlation decay for DRH indicate strong sequence-dependent variations in local helical diameter, reflecting fluctuations between A- and B-form geometries (see Figure S24). The cgNA+ model reproduces the MD results closely for all three systems, accurately capturing both the decay rate, which reflects bending persistence, and the oscillation period associated with the underlying helical structure.

**Figure 9.**
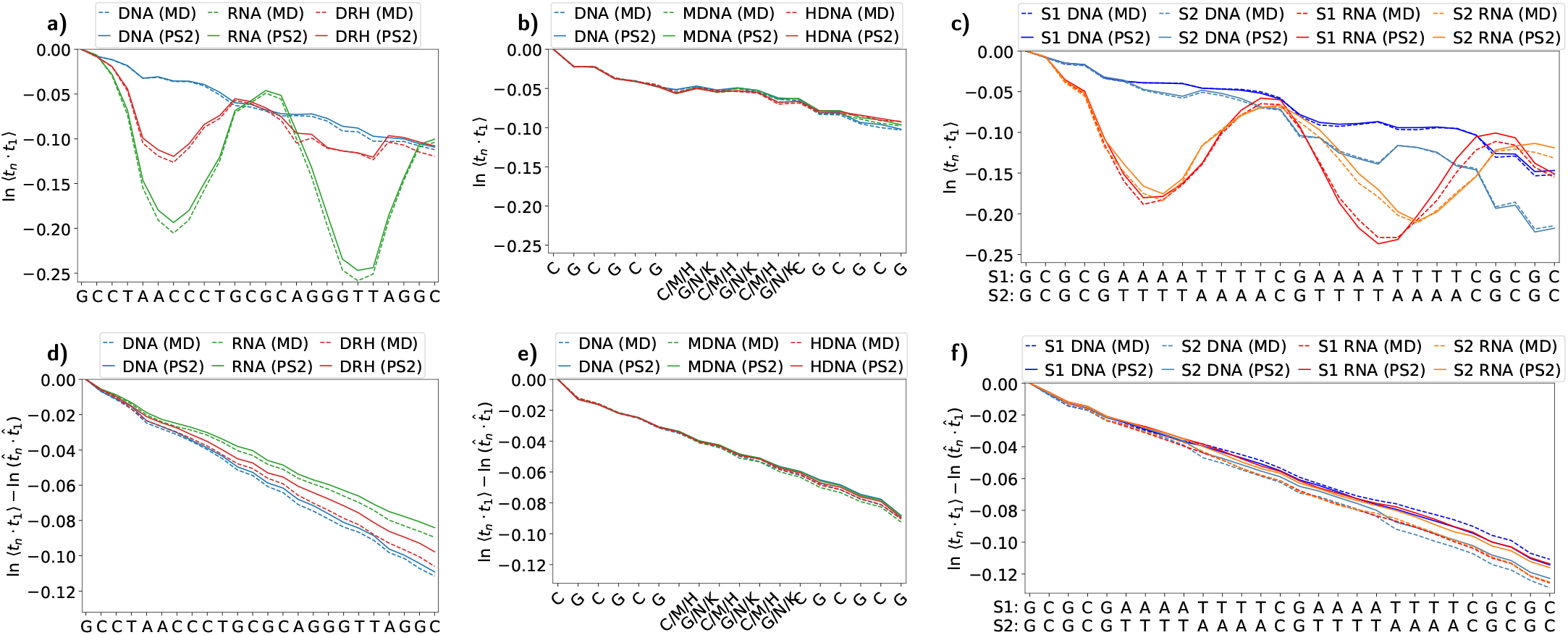
Comparison of the logarithm of the tangent–tangent correlation (Eq. 12) obtained from Monte Carlo sampling of the Gaussian pdfs inferred from MD and cgNA+ for **(a)** the same 24-mer sequence in dsDNA, DRH, and dsRNA (U in place of T) versions, **(b)** poly(CG) sequence along with its methylated and hydroxymethylated versions, and **(c)** two different A-tract sequences (S1 and S2) in dsDNA and dsRNA (U in place of T) versions. **t**_*i*_ denotes the unit tangent vector associated with base-pair frame 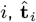 denotes the unit tangent vector associated with base-pair frame *i* in the groundstate, and ⟨·⟩ denotes an average over the Monte Carlo ensemble. In **(a–c)**, linear fits to the tangent–tangent correlation decay yield the inverse apparent persistence length (*l*_*p*_). **(d–f)** show the corresponding tangent–tangent correlation plots after factoring out the contribution of the equilibrium shape, yielding the inverse dynamic persistence length (*l*_*d*_).

Figure 9b presents the analogous comparison for a poly(CG) sequence and its methylated and hydroxymethylated variants, again revealing an excellent agreement between the tangent–tangent correlation decay in cgNA+ and MD. Finally, Figure 9c compares the tangent–tangent correlation for two distinct A-tract motifs in dsDNA and dsRNA forms. The characteristic differences due to distinct helical geometry in dsDNA and dsRNA are evident and are accurately reproduced by cgNA+. Furthermore, the two A-tract motifs, despite identical GC content, exhibit markedly different decay profiles, underscoring the strong sequence context dependence. Figure 9d-e present the tangent–tangent correlation decay after removal of the equilibrium-shape contribution. As discussed in Materials and Methods the linear fit to this data provides the dynamic persistence length *l*_*d*_ for the sequence fragments.

Finally, we note that Laeremans et al. [100] compared cgDNA+ stiffness with all-atom MD simulations in Fourier space, showing that cgDNA+ closely reproduces the q-dependent stiffnesses of the all-atom model and provides a suitable description of local elastic behaviour as well as over longer length scales.

#### dsRNA is stiffer than dsDNA, intermediate stiffness of DRH is driven by DNA-strand pyrimidine-content

Motivated by the close agreement between MD observations and cgNA+ predictions for various test sequences described in the previous Section, we next present spectra of cgNA+ predicted persistence lengths over millions of sequences for both apparent *l*_*p*_ and dynamic *l*_*d*_ persistence lengths, as introduced in Materials and Methods. We reiterate that it is known that the classic apparent persistence length *l*_*p*_ is affected by both the stiffness of the sequence fragment and by intrinsic bends in the groundstate of the fragment, with exceptionally low values typically meaning that the fragment is exceptionally bent. In contrast the dynamic persistence length is more directly tied to only the overall stiffness of the sequence fragment.

Figure 10a compares the persistence-length spectra obtained from two million random sequences of length 220 bp for each of dsDNA, dsRNA, and DRH. The ranges of *l*_*p*_ found are 138, 175, and 166 bp for dsDNA, dsRNA, and DRH, respectively, which are substantial ranges relative to their corresponding mean values of 210.9, 213.9, and 182.4 bp. The distribution is highly asymmetric with the long tails to the left arising for sequences with high intrinsic bends, and therefore exceptionally low apparent persistence length. In contrast, the distributions of *l*_*d*_ are considerably narrower, with ranges of 50, 76, and 106 bp and mean values of 226.1, 269.7, and 236.5 bp for dsDNA, dsRNA, and DRH, respectively. Notably, the ranges of *l*_*p*_ are broadly comparable among dsDNA, dsRNA, and DRH, suggesting that the sequence-dependent variation in intrinsic shape is similar across the three dsNAs, while the range of *l*_*d*_ is markedly larger for DRH than that for either dsDNA or dsRNA, suggesting that the coexistence of DNA and RNA strands renders hybrid duplexes particularly sensitive to changes in overall stiffness with sequence.

**Figure 10.**
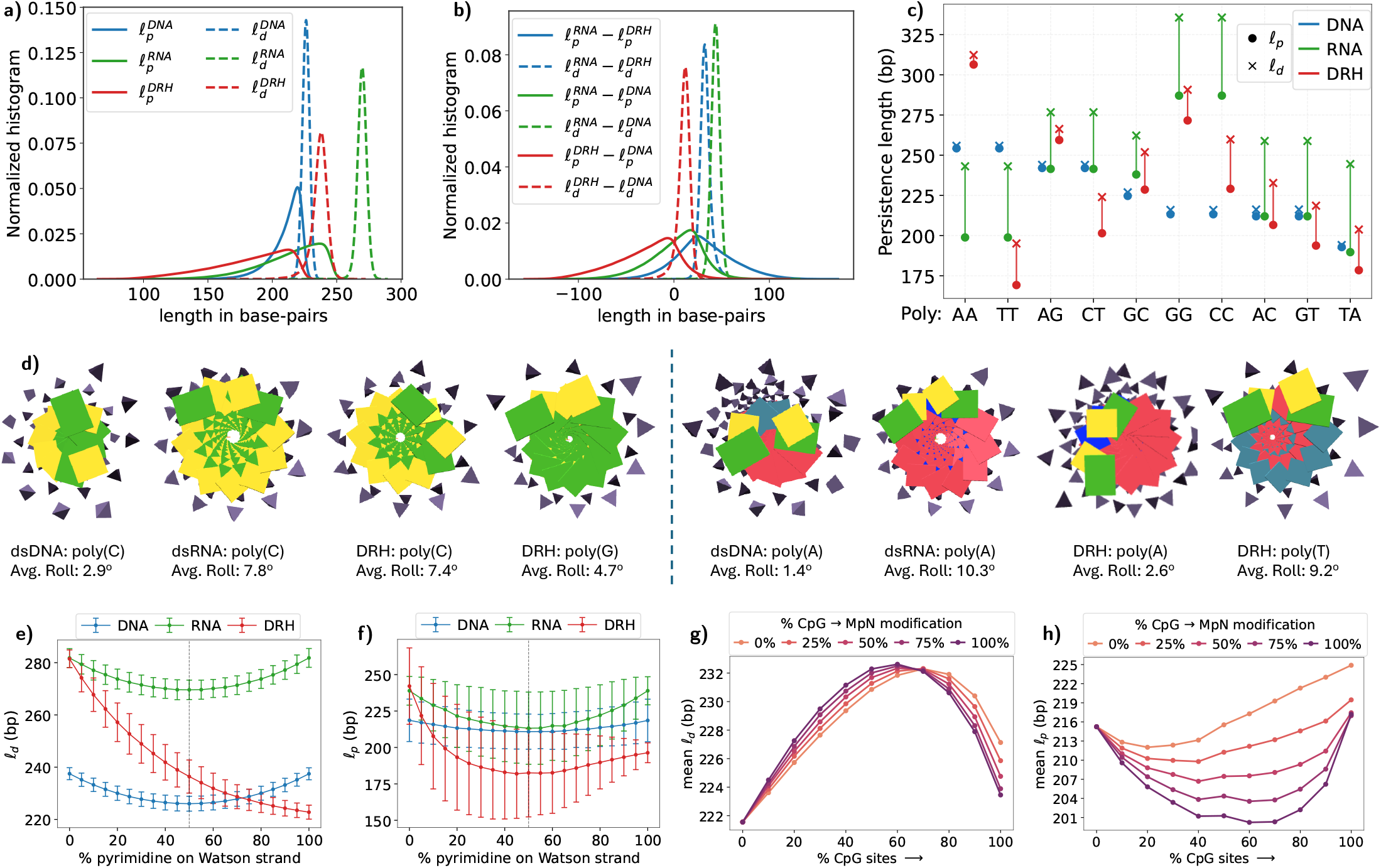
Sequence-dependent persistence lengths across dsNA. **(a)** Histograms of apparent (*l*_p_) and dynamic (*l*_d_) persistence lengths for two million random sequences (of length 220 bp) in dsDNA, dsRNA, and DRH. The same sequence set was used across all dsNA types. **(b)** Sequence-wise differences in persistence lengths between dsRNA/DRH and dsDNA. **(c)** *l*_d_ and *l*_p_ for all independent poly-dimers ((XY)_110_ embedded in GpC ends) across dsDNA, dsRNA, and DRH. **(d)** 3D equilibrium shapes for C_220_ and A_220_ in its dsDNA, dsRNA, and DRH with GpC ends, together with G_220_ and T_220_ for DRH, taken from the https://cgdnaweb.epfl.ch website. **(e)** *l*_*d*_ and **(f)** *l*_*p*_ as a function of Watson strand pyrimidine content for dsDNA, dsRNA, and DRH. Statistics (mean and standard deviation) are obtained from 105,000 random sequences of length 220 bp with varying pyrimidine content in increments of 5% (5000 sequences for each). The vertical dashed line marks 50% Watson-strand pyrimidine content, at which pyrimidine and purine contents are equal on both strands. Sequence compositions on either side of this line are reverse complement mirror and therefore correspond to identical sequence compositions for dsDNA and dsRNA, but to distinct chemical compositions for DRH. Persistence lengths, *l*_*d*_ **(g)** and *l*_*p*_ **(h)** as a function of C_p_G content and impact of symmetric C_p_G methylation. Data points represent mean across random sequences of length 220 bp with varying C_p_G content in increments of 10% (10000 sequences for each). Within each set, 0%, 25%, 50%, 75%, or 100% of the C_p_G steps were randomly methylated. Corresponding plots for symmetric hydroxymethylation are provided in Figure S26.

Figure 10b presents the same data differently by plotting, sequence by sequence, the differences between all three dsNA types of both apparent and dynamic persistence lengths. The range of differences of apparent persistence lengths, solid lines, has no specific sign, suggesting that there is no simple ordering between the apparent persistence lengths for the analogous sequences in the three types of dsNA. Most likely there is no simple rule how the groundstate changes when the dsNA type changes. In contrast the distributions of differences in dynamic persistence lengths are strongly weighted to positive values. There are only 18 exceptional sequences with a negative difference between dsRNA and dsDNA dynamic persistence length (out of two million sequences), and 26 exceptions for the difference between dsRNA and DRH, but many more for DRH and dsDNA. This data suggests a strong, but not strict, ordering between dynamic persistence length values *l*_*d*_ for the analogous sequences

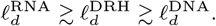

In the precursor cgDNA model [60], and with parameters trained on different protocol MD simulations, it was observed for dsDNA that the single highest outlier, and the single lowest outlier in the dynamic persistence length spectra were the homopolymer sequence dsDNA poly(A) and dsDNA alternating poly(AT) [66]. That remains the case for spectra computed here over two million random sequences. The dynamic and apparent persistence lengths for all independent poly(XY) fragments are presented in Figure 10c. Specific homopolymers or polydimers are often the stiffest and softest sequences across dsNA (compared to 2 million random sequences). For dsDNA, the stiffest sequence is poly(A)/poly(T), with *l*_*d*_ ≈ 256 bp, whereas poly(AT) is the softest, with *l*_*d*_ ≈ 194 bp. Notably, both sequences have straight groundstates, despite exhibiting a large difference in rigidity. For dsRNA, the stiffest sequence is poly(C)/poly(G), with *l*_*d*_ ≈ 336 bp, whereas the softest poly-dimer In the precursor cgDNA model [60], and with parameters trained on different protocol MD simulations, it was observed for dsDNA that the single highest outlier, and the single lowest outlier in the dynamic persistence length spectra were the homopolymer sequence dsDNA poly(A) and dsDNA alternating poly(AT) [66]. That remains the case for spectra computed here over two million random sequences. The dynamic and apparent persistence lengths for all independent poly(XY) fragments are presented in Figure 10c. Specific homopolymers or polydimers are often the stiffest and softest sequences across dsNA (compared to 2 million random sequences). For dsDNA, the stiffest sequence is poly(A)/poly(T), with *l*_*d*_ ≈ 256 bp, whereas poly(AT) is the softest, with *l*_*d*_ ≈ 194 bp. Notably, both sequences have straight groundstates, despite exhibiting a large difference in rigidity. For dsRNA, the stiffest sequence is poly(C)/poly(G), with *l*_*d*_ ≈ 336 bp, whereas the softest poly-dimer (not the overall softest sequence) is poly(A), with *l*_*d*_ ≈ 242 bp. For comparison, the lowest 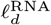observed among random sequences is 213 bp. For DRH, the trend is particularly striking: the stiffest sequence is poly(A), with *l*_*d*_ ≈ 312 bp, whereas poly(T) is the softest poly-dimer, with *l*_*d*_ ≈ 195 bp. The lowest 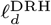 observed among random sequences is 166 bp. A similar asymmetry in complementary poly-dimers is also observed for other poly-dimer sequence pairs in DRH, such as poly(C) and poly(G), poly(TG) and poly(AC), and poly(TC) and poly(AG). These pairs correspond to the same physical dsDNA and dsRNA, but represent distinct molecules in DRH (Figures 10d and S25). In DRH, both *l*_*d*_ and *l*_*p*_ can differ substantially within each pair. This behaviour highlights that the flexibility and rigidity of DRH cannot be understood as a simple average of dsDNA and dsRNA properties; rather, DRH exhibits a distinct and more complex sequence-dependent mechanical pattern.

With only negligible exceptions, just 18 sequences out of two million, for which 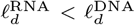, poly(A) stands out as the most extreme case, with *l*_*d*_ being 13 bp larger in dsDNA than in dsRNA. Moreover, poly(A) is also the stiffest sequence in DRH and, notably, the only homopolymer for which DRH stiffness exceeds that of both dsDNA and dsRNA. Together, these observations identify poly(A) as a striking outlier.

Figure 10e plots *l*_*d*_ across dsNA as a function of pyrimidine content on the Watson strand, with statistics computed from 5000 sequences at each pyrimidine fraction from 0 to 100% in increments of 5%. Relative to DRH, *l*_*d*_ in dsDNA and dsRNA varies only weakly and is symmetric about 50% Watson-strand pyrimidine content. This reflects the fact that sequence compositions on either side of 50% Watson-strand pyrimidine content are reverse-complement mirrors and therefore chemically equivalent in dsDNA and dsRNA. In contrast, 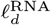 changes much more strongly: at low pyrimidine content it is closer to 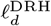, with *l*_*d*_ ≈ 282 bp, whereas with increasing pyrimidine content it monotonically decreases toward 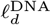, reaching approximately 223 bp. This trend is consistent with the configurational volume reported in Figure S16, where in DRH purine–purine steps are highly rigid and complementary pyrimidine–pyrimidine steps are highly flexible. *l*_*p*_ shows a similar trend (Figure 10f), although there is a much larger variation over sequence space because *l*_*p*_ also has intrinsic shape contribution. For example, roll varies strongly with sequence (Figure S8) and directly affects *l*_*p*_.

We emphasize that the persistence lengths predicted by the cgNA+ model are considerably higher than the experimental consensus value of approximately 150 bp for dsDNA. Nevertheless, the sequence-dependent trends are broadly consistent with those observed experimentally [66]. One possible explanation for the discrepancy is the difference in ionic conditions: experimental measurements are often performed at higher salt concentrations and frequently in the presence of divalent counterions, whereas the underlying MD simulations typically use monovalent ions at near-physiological concentrations. Since ionic conditions strongly affect dsNA flexibility, this can substantially influence the measured persistence length [35, 101, 102]. In addition, DNA parameterization in commonly used MD force fields may itself be somewhat too stiff [73, 103, 104]. Specifically, Roldán-Piñero et al. [104] reported a systematic overestimation of the persistence length of polydimer sequences across various atomistic force fields, including parmbsc1. More importantly, tangent–tangent correlation in samples drawn from the MD and cgNA+ inferred Gaussian distributions are extremely close (Figure 9), suggesting that the discrepancy is not intrinsic to the cgNA+ model itself.

The sequence space explored experimentally for dsRNA and DRH is quite limited. dsRNA is generally considered stiffer than dsDNA, with experimentally reported mean persistence lengths of 62–64 nm (∼205–211 bp) under near-physiological salt conditions [33]. For DRH, Zhang et al. [35] reported a persistence length of approximately 63 nm (∼210 bp) at 1 mM NaCl and showed that the persistence length of DRH generally lies between that of dsDNA and dsRNA. These observations are broadly consistent with our findings. At the same time, cgNA+ reveals a strong sequence dependence of persistence length, demonstrating variability that remains difficult to resolve using current experimental approaches.

#### Persistence length increases upon C_p_G modification but decreases upon hypermodification

Next, we quantified the effect of C_p_G modifications on the persistence length of dsDNA by analysing 55 ensembles of 10K sequences, all of length 220 bp. First, for each prescribed C_p_G content from 0% to 100%, in increments of 10%, we generated 11 ensembles of 10K sequences. Then within each of the sets of 10K, five different versions were created by randomly symmetrically methylating 0%, 25%, 50%, 75%, or 100% of the C_p_G steps. Other ensembles were created by similarly hydroxymethylating. Finally for each of the resulting 55 ensembles of 10K sequences, we computed an average dynamic persistence length *l*_*d*_ over the ensemble, to arrive at the data presented in Figure 10g. It can be seen that at each fixed methylation concentration as the fraction of C_p_G steps increases, the average *l*_*d*_ rises until the C_p_G content reaches 60-70%, after which it begins to decline. This trend likely reflects an initial increase in C/G content, accompanied by a reduction in highly flexible steps such as T_p_A (Figure S16d). However, at higher C/G content, the increasing number of flexible C_p_G steps itself reduces *l*_*d*_. For all the sequence ensembles at up to 60 percent fixed C_p_G content, increasing the percentage of C_p_G methylation (C_p_G to M_p_N) consistently increases the ensemble average *l*_*d*_. But at higher C_p_G content the trend reverses and increasing the methylation percentage decreases the average *l*_*d*_. Experimental studies have generally suggested that methylation increases the persistence length of dsDNA [19, 47]. However, more recent work has reported that hypermethylation can instead enhance dsDNA flexibility [32, 34]. In particular, Shon et al. observed that persistence length increases with C/G content and also rises upon C_p_G methylation up to approximately 50% C_p_G density, but methylation at higher C_p_G density leads to a drastic decrease in persistence length. Figure 10g reveals that cgNA+ predicts a similar biphasic trend for dynamic persistence length.

Figure 10h shows the analogous data for the apparent persistence length *l*_*p*_, which, we recall, is affected by both stiffness and intrinsic shape. For any fixed methylation percentage, the ensemble average of *l*_*p*_ first decreases and then increases with the fraction of C_p_G steps. At low epigenetic modification densities, more C_p_G steps increase the likelihood of forming A-tract-like motifs that can induce localized bends in the sequence. This is also reflected in the growing difference between *l*_*p*_ and *l*_*d*_ as the fraction of modified C_p_G steps increases, indicating that these modifications enhance intrinsic bending in the groundstate. At higher C_p_G densities, however, *l*_*p*_ increases even though *l*_*d*_ continues to decrease. By the time all C_p_G steps are modified, the difference between *l*_*p*_ and *l*_*d*_ becomes nearly negligible, consistent with an almost straight groundstate.

C_p_G methylation reduces *l*_*p*_ and hydroxymethylation leads to an even larger reduction (Figure S26). The minimum average *l*_*p*_ is approximately 201 bp for sequences with 60% C_p_G steps (all methylated), compared with about 187 bp for the corresponding hydroxymethylated sequences. This trend can be explained by the increase in roll at modified C_p_G steps (Figure S13), which bends the dsDNA groundstate more strongly. Figure S25e shows the tangent-tangent correlation decay for (*CG*)_110_, (*MN* )_110_, and (*HK*)_110_, all with GpC ends. The roll magnitude follows the order HK (10.3^°^) *>* MN (10.0^°^) *>* CG (5.7^°^), and the extent of deviation from the intrinsically straight groundstate follows the same trend, directly accounting for the observed ordering of persistence lengths.

## Conclusions and discussion

In this work, we described cgNA+, a sequence-dependent coarse-grained model for dsNA, and used it to construct a first large-scale comparative map of sequence-dependent mechanical landscapes across dsDNA (including epigenetic C_p_G modifications), dsRNA, and DRH. A primary result is that cgNA+ predicts non-local sequence-dependent equilibrium shape and stiffness matrices with errors an order of magnitude below the sequence-variability. Critically, the model captures non-local conformational changes arising from a single point mutation or from epigenetic modification at an isolated C_p_G step that propagate several base-pair positions in both directions from the site of chemical change. Although the cgNA+ model is parameterised locally at the level of dimer-steps, non-local effects emerge naturally through mechanical coupling between neighbouring structural units, which redistributes deformation across the duplex and gives rise to sequence effects beyond nearest-neighbour interactions.

The equilibrium shape of a base-pair step is strongly modulated by flanking sequence context, to such an extent that, for certain dimer steps, the effect of neighbouring base pairs is comparable to, or even greater than, that of changing the base-pair step itself. This flanking context-induced deformation at a given dimer step correlates strongly with the flexibility of the dimer step. The physical interpretation is intuitive: a mechanically soft dimer can be more easily deformed by the structural preferences of its neighbours. Pyrimidine–purine steps, and TA in particular, emerge as the most sensitive to flanking context in dsDNA, consistent with prior observations [10, 39, 49]. The same qualitative trend holds for dsRNA, but the absolute deviations are smaller, reflecting the greater conformational rigidity of the A-form helix. These findings reinforce the importance of going beyond dimer-level descriptions as also reported earlier [5, 6, 37].

Furthermore, C_p_G modifications alter the dimer’s groundstate comparable in magnitude to those caused by SNPs, and are therefore non-trivial. Given that SNPs are the most common form of genetic variation in the human genome and that their structural effects are increasingly recognised as contributors to differential protein-binding affinity and disease susceptibility through indirect readout mechanisms [17], this finding supports the view that epigenetic modifications can act as mechanical signals, rather than purely chemical ones [4, 91, 105].

The groove width analysis provides a structurally intuitive and biologically consequential perspective on sequence-dependent mechanics. The decamer-scale groove width spectra confirm and quantify the characteristic groove geometries of the three dsNA classes: dsDNA exhibits a wide major groove and narrow minor groove consistent with B-form geometry; dsRNA shows an inverted pattern with a narrow major groove and wide minor groove characteristic of A-form; and DRH occupies an intermediate position in both dimensions [43, 44]. Importantly, the groove width *distributions* are wide, demonstrating that groove geometry is not a fixed property of the helix type but is strongly modulated by sequence.

The sequence determinants of extreme groove widths qualitatively agree with crystallographic and NMR observations [17, 93]. For dsDNA, A/T-rich sequences and in particular A-tract motifs lacking TA steps are associated with narrow minor grooves, whereas G/C- rich sequences favour wider minor grooves. The narrow minor groove of A-tracts is associated with a strong negative electrostatic potential that facilitates recognition by positively charged arginine side chains [17], underpinning a widely used mode of protein–DNA interaction. The application to CTCF binding sites provides a direct biological illustration. We find that predicted CTCF binding sites in the human genome exhibit a statistically distinct groove width profile relative to GC-matched controls, with differences confined to the core 19-bp motif.

A large-scale analysis encompassing millions of sequences and lengths exceeding 200 base pairs reveals that the three nucleic-acid classes examined here occupy distinct mechanical landscapes. In general, dsRNA is mechanically stiffer than dsDNA, while DNA:RNA hybrids display intermediate but non-trivial behaviour. In several observables, hybrids do not behave like an average of DNA and RNA. Instead, they exhibit their own sequence-dependent signatures, reflecting the asymmetry between the DNA and RNA strands and the different structural preferences they impose. This is evident both in equilibrium geometry and in persistence-length behaviour, where hybrids can display markedly asymmetric responses depending on which strand carries a given base identity.

The persistence-length analysis further emphasizes the importance of sequence in determining dsNA mechanics. Both the apparent persistence length and the dynamic persistence length vary substantially across sequence space for all three nucleic-acid classes. While dsRNA is, on average, the stiffest class, and dsDNA the most flexible in dynamic terms, the spread within each class is large enough that sequence-specific effects are clearly biologically relevant. In DRH especially, the broad distribution of persistence lengths indicates a mechanically rich landscape that cannot be reduced to simple interpolation between dsRNA and dsDNA behaviour. These results support the view that sequence influences not just local geometry, but mesoscopic mechanical properties that are likely to matter for packaging and function.

Finally, we note that a full atomistic representation can be readily reconstructed from any cgNA+ configuration by fitting the ideal atomistic coordinates onto the coarse-grained structure. Sugar moieties can likewise be reconstructed, either through energy minimisation [63] or using deep-learning models trained on atomistic MD data [64]. This provides a convenient route to obtaining an equilibrium atomistic representation for any sequence, which can be particularly useful for initiating MD simulations of complex DNA structures such as minicircles, for which the equilibrium shape can be readily obtained using cgNA+min [72].

At the same time, several limitations of the cgNA+ framework should be kept in mind. First, cgNA+ is trained on only first and second moments observed from MD simulations. This does not imply that the underlying equilibrium distribution need be Gaussian, just that the distribution needs to have finite first and second moments, which is a very weak restriction. Certainly large conformational transitions, strand separation, base flipping, melting, or kinking are all multi-well and therefore non-Gaussian phenomena. On the other hand it is a general and powerful principle of maximum entropy estimation that if the only input information for modelling are first and second moments, then the maximum entropy, or least biased, estimator will be a Gaussian model. Another view point is to say that cgNA+ is a predictive model of the sequence-dependent first and second moments of an underlying atomistic distribution that itself is non-Gaussian, and in that interpretation the maximum entropy description should be Gaussian. Thus if it was wished to generalize to a non-Gaussian coarse grain description, then higher moments than second of the atomistic simulations would have to be observed and passed as inputs to the model fitting.

Second, although the model accurately reproduces the atomistic MD observations (via first and second moments), its predictions reflect the underlying atomistic MD force fields and solvent conditions; discrepancies between MD and experiment may therefore propagate into the cgNA+ model. One specific restriction of the MD simulations that have so far been used to train cgNA+ parameter sets is that all of the protocols have involved only monovalent counter ions. Finally, the current model focuses on linear double-stranded fragments in solution and does not by itself represent protein binding, self-avoidance, or other higher-order interactions unless these are introduced in an additional modelling layer. One step in this direction is the recent modelling of covalently closed minicircles using the cgNA+ model energy. In particular it was shown how to enforce closure of the two ends of a dsNA on itself in a computationally efficient way starting from the cgNA+ internal coordinates [72].

These limitations also point to several important directions for future work. Extending the framework to broader chemical alphabets, additional epigenetic marks, and base mismatches would widen its applicability and allow a more comprehensive description of chemically diverse nucleic-acid systems. At the same time, incorporating non-Gaussianity, for example through a multi-harmonic formulation [106], could provide a more realistic treatment of the inherent bimodality observed in some of the structural parameters [5, 37, 64] (although those models should be constrained by the fact that their first and second moments should match those of cgNA+). A further natural step is to couple cgNA+ with higher-level models of nucleosomes, chromatin fibres, and DNA minicircles, which would make it possible to investigate how sequence-encoded mechanical properties propagate to the larger scales relevant to genome organisation. Encouragingly, progress in this direction has already begun to emerge [67–69, 72, 105].

More broadly, the study highlights the utility of coarse-grained statistical mechanics models trained on atomistic data. Direct MD remains the most detailed route to sequence-dependent nucleic-acid mechanics, but it is prohibitively expensive for systematic exploration of large sequence ensembles. cgNA+ provides a practical solution: it accurately captures non-local sequence-dependent mechanics while enabling rapid inference over millions of sequences. By enabling large-scale access to sequence-dependent structural and mechanical landscapes, cgNA+ bridges detailed molecular modelling with genome-scale analysis, supporting statistical and quantitative understanding of how sequence and chemical modification encode nucleic-acid mechanics relevant to biological function.

## Code and data availability

The cgNA+ software packages in MATLAB and Python are available at https://github.com/rahul2512/cgNA_plus_Matlab and https://github.com/rahul2512/cgNA_plus. The cgNA+mc (C++ scripts) package for the Monte Carlo simulation is available at https://github.com/rahul2512/cgNA_plus_mc. The code for estimating cgNA+ parameters, starting from oligomer-level statistics for a set of sequences, is provided at rahul2512/cgNA_plus_training. The analysis code and data are available at 10.5281/zenodo.21892342, and the results can be reproduced by running Sequence_space.ipynb.

## Author contributions

R.Sharma: Conceptualization, Data curation, Formal analysis, Investigation, Methodology, Project administration, Software, Validation, Visualization, and Writing (original draft). A.S.P.: Conceptualization, Methodology, Software, and Writing (review & editing). R.Singh: Validation and Writing (original draft). D.P.G.: Validation and Writing (review & editing). O.G.: Conceptualization, Methodology, Validation, and Writing (review & editing). J.H.M.: Conceptualization, Funding acquisition, Investigation, Methodology, Validation, Resources, Supervision, and Writing (original draft).

## Footnotes

1 The importance of the dimension 42 has been pointed out previously by D. Adams.

