## Supplementary Information for "Sequence-dependent conformational and mechanical landscapes of double-stranded nucleic acids"

Rahul Sharma*[1], Alessandro S. Patelli[1], Raushan Singh[2], Daiva Petkevičiūtė-Gerlach[3], Oscar Gonzalez[4], and John H. Maddocks*[1]

[1]Institute of Mathematics, École Polytechnique Fédérale de Lausanne, Lausanne, Switzerland
[2]Department of Mechanical Engineering, IIT Madras, Chennai, India
[3]Department of Applied Mathematics, Kaunas University of Technology, Kaunas, Lithuania
[4]Department of Mathematics, University of Texas, Austin, USA

**4 Supplementary tables** **18**

**5 Supplementary figures** **19**

**References** **42**

### 1 Details of molecular dynamics simulations

cgNA+ parameter sets were independently obtained for dsDNA, dsRNA, DNA:RNA hybrids (DRHs), and epigenetically modified dsDNA using all-atom molecular dynamics (MD) simulations of carefully designed sequence libraries. The training libraries were constructed to provide broad coverage of local sequence environments while keeping the total number of MD simulations computationally tractable. Independent test libraries were used to assess model accuracy by comparing cgNA+ predictions against the corresponding MD simulations. Complete lists of the training and test sequences are provided in Sections 1.1–1.3 and Tables S1–S3. None of the test sequences were used during parameter estimation.

Initial dsDNA and dsRNA structures were generated using NAB in AMBERTOOLS [1]. DNA duplexes were constructed using Arnott B-form fibre parameters, whereas RNA duplexes were constructed using Arnott A-form fibre parameters. DRH structures were generated from A-form RNA duplexes by replacing uracil with thymine on the DNA strand and converting the corresponding ribose sugars to deoxyribose. Methylated and hydroxymethylated dsDNA sequences were generated by modifying the corresponding cytosine residues using the LEaP module of AMBERTOOLS [1]. The dsDNA molecules were described using the parmbsc1 corrections [2] along with parmbsc0 and parm99 parameters [3, 4], whereas the dsRNA molecules were described using the OL3 corrections [5]. For DRH, the DNA and RNA strands were described using the corresponding DNA and RNA force-field parameters, respectively. Additional force-field parameters were included for methylated and hydroxymethylated cytosine residues [6, 7]. All molecules were solvated in explicit TIP3P water [8] using a truncated octahedral box with a minimum solvent buffer of 10 Å. The solvated molecules were neutralised with $K^+$ ions, and additional KCl was added to obtain an ionic strength of approximately 150 mM. Potassium and chloride ions were described using the Joung–Cheatham parameters [9].

Following energy minimisation, the systems were heated to 300 K and equilibrated for 50 ps. Production simulations were subsequently performed in the NPT ensemble at 300 K and 1 atm using AMBER18 [10]. Simulations employed a 2 femtoseconds integration time step, periodic boundary conditions, particle-mesh Ewald electrostatics, and a 9 Å real-space cutoff. Trajectory snapshots were saved every 2 picoseconds.

Finally, we applied a filtering procedure to remove any MD snapshot containing one or more broken hydrogen bonds. Consistent with previous studies [11, 12, 13, 14], a hydrogen bond was classified as broken when either

1. the distance between the donor and acceptor heavy atoms exceeded 4 Å, or
2. the donor–hydrogen–acceptor angle was smaller than 120°.

The retained snapshots were subsequently converted into large ensembles of sets of cgNA+ internal configuration coordinates, and then used to estimate the MD-derived first and second moments employed for cgNA+ parameter estimation.

#### 1.1 Sequence library for interior parameter blocks

For dsDNA and dsRNA, we used the palindromic sequence library introduced in Ref. [15] (Table S1). The 16 training sequences were designed to provide balanced statistics for all interior dimer junctions and collectively contain all 256 possible tetramers on the reading strand. GC terminal base pairs were used throughout to minimise end fraying. The palindromic sequences (reading strand invariant) improve statistical efficiency as equivalent sequence environments are sampled on both strands which allow quantifying for convergence of MD trajectories. The same library was also used for DRH, with the DNA strand designated as the reading strand. Although a palindromic library cannot be constructed for DRHs because the two strands have different chemistries.

Independent test sequences were used to assess model transferability and accuracy. These include random palindromes, A-tracts, poly(A) and poly(AT) tracts, CpG-rich sequences, single-base substitutions, and longer random duplexes. None of the test sequences were used during parameter estimation.

Table S1: Palindromic sequence library used for cgNA+ parameterisation and validation. dsDNA sequences are represented using the standard A, T, C, and G alphabet. The corresponding dsRNA library is obtained by replacing T with U. The same sequence set was also used for DRHs, with the DNA strand designated as the reading strand.

| Index | MD Sequence |
|---|---|
| | Training sequences |
| 1 | GCTTAGTTCAAATTTGAACTAAGC |
| 2 | GCTCTCTGTATTAATACAGAGAGC |
| 3 | GCCCTTGGCGATATCGCCAAGGGC |
| 4 | GCTAAAGCCTTATAAGGCTTTAGC |
| 5 | GCGGTAGAAAACGTTTTCTACCGC |
| 6 | GCCAAGACATTGCAATGTCTTGGC |
| 7 | GCAGATGGTCAGCTGACCATCTGC |
| 8 | GCCTCACCGCTCGAGCGGTGAGGC |
| 9 | GCAGTGGAATCATGATTCCACTGC |
| 10 | GCTTTACTTCGTACGAAGTAAAGC |
| 11 | GCTACCTATGCTAGCATAGGTAGC |
| 12 | GCGCACTGGGGATCCCCAGTGCGC |
| 13 | GCTGAGGAGTCCGGACTCCTCAGC |
| 14 | GCTGCCGTCGGGCCCGACGGCAGC |
| 15 | GCGCACAACACGCGTGTTGTGCGC |
| 16 | GCCTAACCCTGCGCAGGGTTAGGC |
| | Test sequences |
| 1 | GCATTACGCTCCGGAGCGTAATGC |
| 2 | GCAAAAAAAAAAAAAAGC |
| 3 | GCATATATATATATATGC |
| 4 | GCGGATTACGCAGGC |
| 5 | GCGGATTCCGCAGGC |
| 6 | GCGCGAAAATTTTCGAAAATTTTCGCGC |
| 7 | GCGCGTTTTAAAACGTTTTAAAACGCGC |
| 8 | GCGCGCGCGCGCGCGCGCGC |
| 9 | CGGCGCACGTGACCGCG |
| 10 | GCATCGCCACTGAAGTTGGTTATAACCAACTTCAGTGGCGATGC |

#### 1.2 Sequence library for modified-base interior parameter blocks

cgNA+ includes parameters for CpG methylation and hydroxymethylation, covering both symmetric modifications, where cytosines on both DNA strands are modified, and asymmetric modifications, where only one cytosine is modified. We denote 5-methylcytosine by M and the guanine paired with methylated cytosine on the complementary strand by N. Similarly, H denotes 5-hydroxymethylcytosine and K denotes the guanine paired with hydroxymethylcytosine.

The training and test sequences used to determine the modified interior dimer parameter blocks are listed in Table S2. The sequences were designed to sample isolated and adjacent modified CpG steps in diverse flanking sequence contexts. The same sequence set was used for both methylated and hydroxymethylated DNA, with the symbols M, N replaced by H, K for the hydroxymethylated sequences.

Table S2: Libraries of methylated and hydroxymethylated dsDNA sequences used for cgNA+ parameterisation and validation. Here, M denotes 5-methylcytosine and N the guanine paired with methylated cytosine on the complementary strand. The same sequence set was also used for hydroxymethylated DNA, with M and N replaced by H and K, respectively, where H denotes 5-hydroxymethylcytosine and K its complementary guanine.

| Index | Sequence | | |
|---|---|---|---|
| | Training library | | Test library |
| 1 | GCTAMNTGTAMNMNTACAMNTAGC | 1 | GCTAMGTGTCMNMNGACACNTAGC |
| 2 | GCATMNACGAMNMNTCGTMNATGC | 2 | GCATMGACGTMNMNACGTCNATGC |
| 3 | GCGCMNGGAGMNMNCTCCMNGCGC | 3 | GCTGMGTTCGMNMNCGAACNCAGC |
| 4 | GCTCMNCTAAMNMNTTAGMNGAGC | 4 | GCCTMGCGTTMNMNAACGCNAGGC |
| 5 | GCTGMNTTCCMNMNGGAAMNCAGC | 5 | GCCTGAGTAMGMNCNTACTCAGGC |
| 6 | GCCTMNCGTGMNMNCACGMNAGGC | 6 | GCGGATTAMNCAGGC |
| 7 | GCGCMGGGATMNMNATCCCNGCGC | 7 | GCGCGCGMNMNMNCGCGCGC |
| 8 | GCTCMGCTACMNMNGTAGCNGAGC | 8 | GCGCGCGMGMGMGCGCGCGC |
| 9 | GCTAMGTGTCCNMGGACACNTAGC | 9 | GCGCGMNCGCGCGMGCGCGC |
| 10 | GCATMGACGTMGCNACGTCNATGC | | |
| 11 | GCAGMGMGATAATTATCNCNCTGC | | |
| 12 | GCCACAAGTCNMNMGACTTGTGGC | | |

#### 1.3 Sequence library for terminal parameter blocks

The palindromic and modified-base libraries described above contain GC as terminal base pairs (to minimize fraying) and were therefore sufficient to determine the interior parameter blocks together with the GC terminal parameter block. For the remaining 15 non-GC terminal dimer steps, additional simulations were required to train parameters. To this end, we constructed the sequence library shown in Table S3. For each non-GC terminal dimer, four 12-bp sequences of the form

XYUV-(hexamer)-GC

were generated, where $XY$ denotes the terminal dimer of interest, $UV$ was selected from the pyrimidine-pyrimidine, pyrimidine-purine, purine-purine, and purine-pyrimidine dimer classes to provide diverse local sequence environments, and the remaining six bases were chosen randomly. The opposite end of each sequence was fixed to GC (to minimize fraying).

Table S3: Sequence library used to parameterise non-GC terminal dimer blocks. For each of the 15 non-GC terminal dimers, four 12-bp sequences were simulated with the terminal dimer placed at one end of the duplex and a GC terminal dimer at the opposite end.

| Index | Sequence | Index | Sequence | Index | Sequence |
|---|---|---|---|---|---|
| 1 | AAGACCACTTGC | 21 | TGAGGCCACCGC | 41 | GTAAGATTACGC |
| 2 | AAGTTTAGGGGC | 22 | TGATCAAGTAGC | 42 | GTGCGACGCTGC |
| 3 | AATGCGTATCGC | 23 | TGTGCCGAGAGC | 43 | GTCAGGATAAGC |
| 4 | AATCACTTAGGC | 24 | TGCTTGATTTGC | 44 | GTTTCTAATAGC |
| 5 | ATAGACCCAAGC | 25 | TCAATTCGACGC | 45 | CGGACTACTCGC |
| 6 | ATGTATCACAGC | 26 | TCACAGCCATGC | 46 | CGGTGCTGCTGC |
| 7 | ATCAGGATAGGC | 27 | TCTGTGCAAAGC | 47 | CGTGGTGGAGGC |
| 8 | ATTTCTAGTGGC | 28 | TCTTGCGTTGGC | 48 | CGTCCTATTGGC |
| 9 | AGAAACTCGTGC | 29 | TTGATACCGCGC | 49 | CCGGCCCGCCGC |
| 10 | AGATAACACTGC | 30 | TTATCATGCAGC | 50 | CCACCCCGTCGC |
| 11 | AGCGCTCGTCGC | 31 | TTTGAATTATGC | 51 | CCTAAGTCTAGC |
| 12 | AGCCATGAAAGC | 32 | TTCTGGTTACGC | 52 | CCTTGCCTACGC |
| 13 | ACGGACGAATGC | 33 | GGGGCTCTTCGC | 53 | CTAGAGCGTGGC |
| 14 | ACGTTCAGTGGC | 34 | GGGTCGGACCGC | 54 | CTGCAACCCAGC |
| 15 | ACCGCGGTGAGC | 35 | GGTATCGACGGC | 55 | CTCATCCAACGC |
| 16 | ACCCAAAGCTGC | 36 | GGCCTATTATGC | 56 | CTCTGAGGTGGC |
| 17 | TAGACACTGTGC | 37 | GAGAGATGTCGC | 57 | CAAAGTCGACGC |
| 18 | TAATCCTCGCGC | 38 | GAATTATTACGC | 58 | CAACCCATTCGC |
| 19 | TATAGTGAGCGC | 39 | GACAGATCACGC | 59 | CACGGAAAGCGC |
| 20 | TATCGGGAATGC | 40 | GACTATGGTAGC | 60 | CATTAACGCCGC |

#### 2 cgNA+ coordinates

This section describes how atomistic MD configurations retained after hydrogen-bond filtering (Section 1) are mapped to the cgNA+ coarse-grained representation. For a duplex of length $N$, one strand is chosen as the reading, or Watson, strand and is written in the $5'$ to $3'$ direction as

$$\mathcal{S} = X_1 X_2 \cdots X_N,$$

where $X_i$ denotes the base at position $i$. The complementary sequence (reading from the Crick strand) is then written in the opposite direction as

$$\bar{\mathcal{S}} = \bar{X}_N \bar{X}_{N-1} \cdots \bar{X}_1.$$

where $\bar{X}_i$ denotes the complementary base of $X_i$. A $\leftrightarrow$ T, A $\leftrightarrow$ U (for dsRNA), and C $\leftrightarrow$ G are complementary base-pairs.

In cgNA+, nucleobases and phosphate groups are treated as rigid bodies, while sugar and solvent are represented implicitly. Each rigid body is described by a position vector $\mathbf{r} \in \mathbb{R}^3$ and an orientation matrix $R \in SO(3)$, where $SO(3)$ is the three-dimensional rotation group, consisting of all orthogonal $3 \times 3$ matrices with determinant one. Base and phosphate reference frames are fitted to atomistic

MD snapshots using a CURVES+ implementation of the weighted least-squares rigid-body fitting procedure described in Ref. [16]. The ideal base-frame geometries were taken from the Tsukuba convention definition [17], and an analogous ideal phosphate geometry was defined in [15].

Table S4: Weighted least-squares rigid-body fitting algorithm used to assign reference frames to nucleobases and phosphate groups. Given two sets of corresponding points $X = \{\mathbf{x}_i\}_{i=1}^n$ and $Y = \{\mathbf{y}_i\}_{i=1}^n$, representing the ideal reference atom coordinates and the corresponding atomistic coordinates in an MD snapshot, respectively, the algorithm computes the rigid-body transformation $(R, \mathbf{r})$ that minimises the weighted fitting error. In this work, all atoms are assigned equal weights, i.e., $w_i^{\text{fit}} = 1$ for $i = 1, \ldots, n$.

| Step | Operation | Formula |
| --- | --- | --- |
| 1 | Compute weighted centroids | $\bar{\mathbf{x}} = \frac{\sum_{i=1}^n w_i^{\text{fit}} \mathbf{x}_i}{\sum_{i=1}^n w_i^{\text{fit}}}, \qquad \bar{\mathbf{y}} = \frac{\sum_{i=1}^n w_i^{\text{fit}} \mathbf{y}_i}{\sum_{i=1}^n w_i^{\text{fit}}}.$ |
| 2 | Translate coordinates to the centroid frame | $\mathbf{p}_i = \mathbf{x}_i - \bar{\mathbf{x}}, \qquad \mathbf{q}_i = \mathbf{y}_i - \bar{\mathbf{y}}.$ |
| 3 | Construct weighted covariance matrix | Define $P = [\mathbf{p}_1, \ldots, \mathbf{p}_n], \qquad Q = [\mathbf{q}_1, \ldots, \mathbf{q}_n],$ and $W = \text{diag}\left(w_1^{\text{fit}}, \ldots, w_n^{\text{fit}}\right).$ The weighted covariance matrix is $S = PWQ^T.$ |
| 4 | Compute singular value decomposition | $S = U\Sigma V^T.$ |
| 5 | Determine optimal proper rotation | $R = V\,\text{diag}\big(1, 1, \det\big(VU^T\big)\big)\,U^T.$ The determinant correction ensures that $\det(R) = 1, \qquad R \in SO(3).$ |
| 6 | Determine optimal translation | $\mathbf{r} = \bar{\mathbf{y}} - R\bar{\mathbf{x}}.$ |

For each rigid body, the fitted transformation $(R, \mathbf{r})$ is obtained by minimising

$$\sum_{i=1}^{n} w_i^{\text{fit}} \left\| \mathbf{r} + R\mathbf{x}_i - \mathbf{y}_i \right\|^2 , \tag{1}$$

where $\{\mathbf{x}_i\}_{i=1}^n$ are the ideal reference atom coordinates, $\{\mathbf{y}_i\}_{i=1}^n$ are the corresponding atomistic coordinates in the MD snapshot, and $w_i^{\text{fit}}$ are atom weights. In this work, equal weights were assigned to all atoms. The resulting optimisation problem is solved using the weighted singular-value-decomposition algorithm summarised in Table S4.

Applying the rigid-body fitting procedure to each retained MD snapshot yields a coarse-grained time-series of base and phosphate frames. For a blunt-ended duplex of length $N$, the two terminal 5′-phosphate groups are absent, so each configuration contains $4N - 2$ rigid bodies. These coarse-grained trajectories provide the absolute-coordinate representation of the duplex and form the starting point for constructing the cgNA+ internal coordinates.

Frame fitting assigns each nucleobase a reference point $\mathbf{r} \in \mathbb{R}^3$ and a right-handed orthonormal frame

$$D = [d_1 \ d_2 \ d_3] \in SO(3).$$

In the Tsukuba convention [17], $d_1$ points approximately towards the major groove, $d_2$ points along the strand direction away from the complementary base, and $d_3 = d_1 \times d_2$ points approximately along the local helical axis.

Because the two strands are antiparallel, the fitted frames associated with complementary bases are naturally not aligned. For modelling, it is convenient to adopt a common orientation convention on both strands as shown in Figure S1a. We therefore transform Crick-strand base frames using

$$P_{\text{flip}} = \begin{bmatrix} 1 & 0 & 0 \\ 0 & -1 & 0 \\ 0 & 0 & -1 \end{bmatrix}, \tag{2}$$

which reverses the $d_2$ and $d_3$ directions while preserving $d_1$. The Watson- and Crick-strand base frames are then represented as

$$D^+ = D, \qquad D^- = DP_{\text{flip}}.$$

After this transformation, the frame axes $d_1^\pm$, $d_2^\pm$, and $d_3^\pm$ have the same geometric interpretation on both strands (see Figure S1a). Throughout this work, superscripts $+$ and $-$ denote rigid bodies associated with the Watson and Crick strands, respectively.

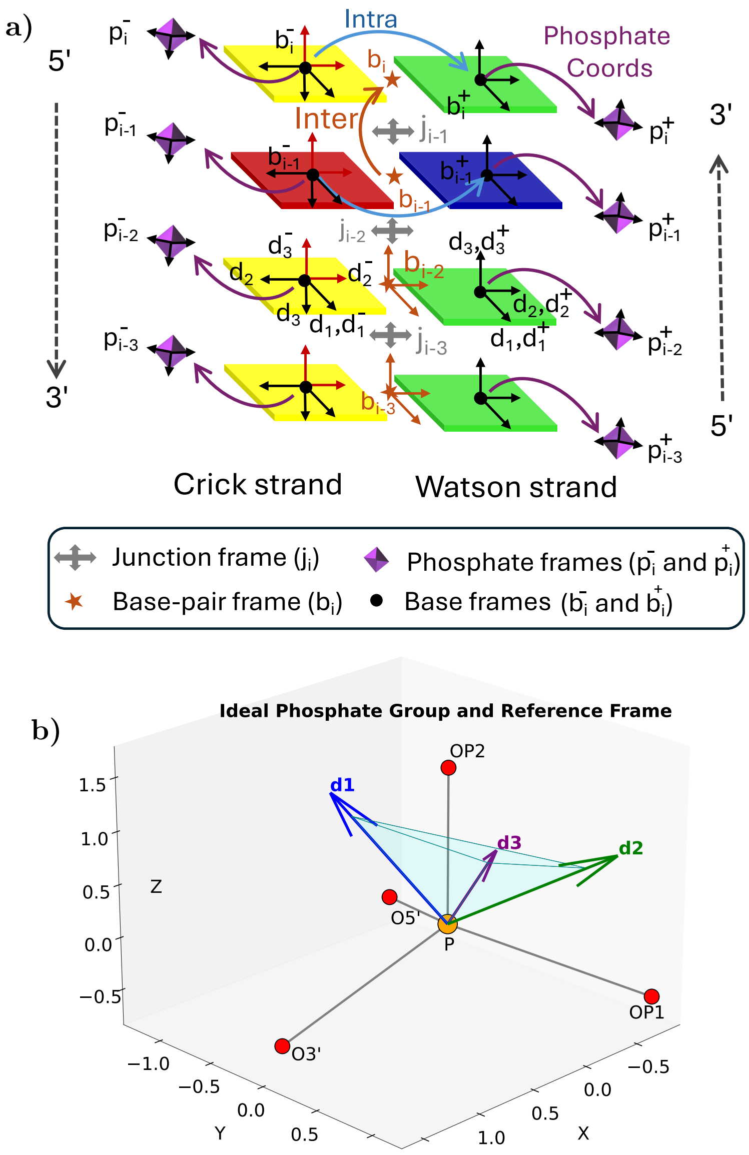

Figure S1: **a)** Schematic representation of an internal fragment of a cgNA+ coarse-grained snapshot. Nucleobases and phosphates are treated as rigid bodies, while sugar moieties are represented implicitly. A rigid frame with three rotational and three translational degrees of freedom is fitted to each base and phosphate by least-squares alignment to reference coordinates, as described in Table S4. Base reference frames follow the Tsukuba convention [17], and analogous phosphate reference coordinates are given in Table S5. Base frames are denoted by $\mathrm{b}_n^{\pm}$ and phosphate frames $\mathrm{p}_n^{\pm}$ for $\mathrm{n}^{\mathrm{th}}$ base-pair level and + denotes the Watson strand (chosen as reading strand) and − denotes the Crick strand. For each base, the frame orientation (shown with black arrows) is described by $D = [d_1\ d_2\ d_3] \in SO(3)$, where $d_1$ points approximately toward the major groove, $d_2$ points along the strand direction away from the complementary base, and $d_3 = d_1 \times d_2$ points approximately along the local helical axis. Because the Watson and Crick strands are antiparallel, the fitted frames associated with complementary bases are not aligned. For modelling purposes, Crick-strand frames are transformed using the flip operation (Eq. 2) so that Watson- and Crick-strand base frames share a common orientation convention; the flipped Crick base frame is shown in red. Base-pair frame $(b_i)$ and junction frame $(j_i)$ are also shown schematically. **b)** Ideal phosphate coordinates and frame. Frame axes are given as $d_3 = \frac{\mathbf{O}_{5'}-\mathbf{O}_{3'}}{\|\mathbf{O}_{5'}-\mathbf{O}_{3'}\|}$, $d_2 = \frac{\mathbf{P}-(\mathbf{O}_{5'}+\mathbf{O}_{3'})}{\|\mathbf{P}-(\mathbf{O}_{5'}+\mathbf{O}_{3'})\|}$, and $d_1 = d_2 \times d_3$.

#### 2.1 Ideal atom coordinates

The ideal coordinates for the standard nucleobases (A, G, T, C, and U) follow the Tsukuba convention [17]. For the modified bases 5-methylcytosine (M) and 5-hydroxymethylcytosine (H), the cytosine reference geometry is used when fitting the corresponding base frames.

The frame axes for ideal phosphate frame (Figure S1b; [15]) are defined by

$$d_3 = \frac{\mathbf{O}_{5'} - \mathbf{O}_{3'}}{\|\mathbf{O}_{5'} - \mathbf{O}_{3'}\|}, \qquad d_2 = \frac{\mathbf{P} - (\mathbf{O}_{5'} + \mathbf{O}_{3'})}{\|\mathbf{P} - (\mathbf{O}_{5'} + \mathbf{O}_{3'})\|}, \qquad d_1 = d_2 \times d_3,$$

where ideal phosphate atom coordinates are listed in Table S5. The corresponding ideal phosphate frame $D^p$ and phosphate reference point $r^p$ are given as

$$D^p = [d_1\ d_2\ d_3] \in SO(3) \text{ and } \mathbf{r}^p = \mathbf{P}.$$

Table S5: Cartesian coordinates (Å) of the non-hydrogen atoms defining the ideal nucleobase and phosphate reference geometries used for rigid-body frame fitting. Coordinates for the nucleobases follow the Tsukuba convention [17], together with analogous ideal phosphate coordinates.

| Atom | Adenine | | | Guanine | | | Phosphate | | |
|---|---|---|---|---|---|---|---|---|---|
| | x (Å) | y (Å) | z (Å) | x (Å) | y (Å) | z (Å) | x (Å) | y (Å) | z (Å) |
| C1′ | -2.479 | 5.346 | 0.000 | -2.477 | 5.399 | 0.000 | — | — | — |
| N9 | -1.291 | 4.498 | 0.000 | -1.289 | 4.551 | 0.000 | — | — | — |
| C8 | 0.024 | 4.897 | 0.000 | 0.023 | 4.962 | 0.000 | — | — | — |
| N7 | 0.877 | 3.902 | 0.000 | 0.870 | 3.969 | 0.000 | — | — | — |
| C5 | 0.071 | 2.771 | 0.000 | 0.071 | 2.883 | 0.000 | — | — | — |
| C6 | 0.369 | 1.398 | 0.000 | 0.424 | 1.460 | 0.000 | — | — | — |
| N6 | 1.611 | 0.909 | 0.000 | — | — | — | — | — | — |
| O6 | — | — | — | 1.554 | 0.955 | 0.000 | — | — | — |
| N1 | -0.668 | 0.532 | 0.000 | -0.700 | 0.641 | 0.000 | — | — | — |
| C2 | -1.912 | 1.023 | 0.000 | -1.999 | 1.087 | 0.000 | — | — | — |
| N2 | — | — | — | -2.949 | 0.139 | -0.001 | — | — | — |
| N3 | -2.320 | 2.290 | 0.000 | -2.342 | 2.364 | 0.001 | — | — | — |
| C4 | -1.267 | 3.124 | 0.000 | -1.265 | 3.177 | 0.000 | — | — | — |
| P | — | — | — | — | — | — | 0.000 | 0.000 | 0.000 |
| O3′ | — | — | — | — | — | — | 1.518 | 0.000 | -0.537 |
| O5′ | — | — | — | — | — | — | -0.759 | -1.315 | -0.537 |
| OP1 | — | — | — | — | — | — | -0.698 | 1.208 | -0.493 |
| OP2 | — | — | — | — | — | — | 0.000 | 0.000 | 1.480 |
| **Atom** | **Thymine** | | | **Cytosine** | | | **Uracil** | | |
| C1′ | -2.481 | 5.354 | 0.000 | -2.477 | 5.402 | 0.000 | -2.481 | 5.354 | 0.000 |
| N1 | -1.284 | 4.500 | 0.000 | -1.285 | 4.542 | 0.000 | -1.284 | 4.500 | 0.000 |
| C2 | -1.462 | 3.135 | 0.000 | -1.472 | 3.158 | 0.000 | -1.462 | 3.131 | 0.000 |
| O2 | -2.562 | 2.608 | 0.000 | -2.628 | 2.709 | 0.001 | -2.563 | 2.608 | 0.000 |
| N3 | -0.298 | 2.407 | 0.000 | -0.391 | 2.344 | 0.000 | -0.302 | 2.397 | 0.000 |
| C4 | 0.994 | 2.897 | 0.000 | 0.837 | 2.868 | 0.000 | 0.989 | 2.884 | 0.000 |
| O4 | 1.944 | 2.119 | 0.000 | — | — | — | 1.935 | 2.094 | -0.001 |
| N4 | — | — | — | 1.875 | 2.027 | 0.001 | — | — | — |
| C5 | 1.106 | 4.338 | 0.000 | 1.056 | 4.275 | 0.000 | 1.089 | 4.311 | 0.000 |
| C5M | 2.466 | 4.961 | 0.001 | — | — | — | — | — | — |
| C6 | -0.024 | 5.057 | 0.000 | -0.023 | 5.068 | 0.000 | -0.024 | 5.053 | 0.000 |

##### 2.2 Frames to cgNA+ internal coordinates

The absolute base and phosphate frames obtained from atomistic snapshots are transformed into internal coordinates that are invariant under rigid-body translations and rotations of the duplex. We first define base-pair and junction frames. For base pair $i$, the base-pair frame $(B_i, g_i)$ is defined by

$$\{B_i, g_i\} = \left\{ D_i^- \sqrt{\Lambda_i}, \frac{1}{2}\left(r_i^+ + r_i^-\right) \right\}, \qquad \Lambda_i = (D_i^-)^T D_i^+, \tag{3}$$

where $\Lambda_i$ is the relative rotation between the Crick- and Watson-strand base frames within base pair $i$. The frame $B_i$ represents the geometric midpoint orientation of the two complementary bases in a base pair, while $g_i$ is their midpoint position.

The junction frame between base pairs $i$ and $i+1$ is defined by

$$\{J_i, t_i\} = \left\{ B_i \sqrt{\Gamma_i}, \frac{1}{2}\left(g_{i+1} + g_i\right) \right\}, \qquad \Gamma_i = B_i^T B_{i+1}, \tag{4}$$

where $\Gamma_i$ is the relative rotation between neighbouring base-pair frames. Thus, $(J_i, t_i)$ represents the geometric midpoint between adjacent base pairs.

To parameterise rotational degrees of freedom, cgNA+ employs the scaled Cayley transformation [13, 18],

$$\mathrm{cay}_\alpha : \mathbb{R}^3 \to SO(3),$$

defined by

$$\mathrm{cay}_\alpha(\rho) = I + \frac{4\alpha\, \rho^\times + 2(\rho^\times)^2}{4\alpha^2 + \|\rho\|^2}, \tag{5}$$

where $\alpha \in \mathbb{R}$ is a scaling factor and $\rho^\times$ denotes the skew-symmetric matrix associated with $\rho = (\rho_1, \rho_2, \rho_3)^T \in \mathbb{R}^3$,

$$\rho^\times = \begin{bmatrix} 0 & -\rho_3 & \rho_2 \\ \rho_3 & 0 & -\rho_1 \\ -\rho_2 & \rho_1 & 0 \end{bmatrix},$$

so that

$$\rho^\times x = \rho \times x \qquad \forall\, x \in \mathbb{R}^3.$$

Equivalently, if

$$u = \frac{\rho}{\|\rho\|}, \qquad \theta = 2 \arctan\left(\frac{\|\rho\|}{2\alpha}\right),$$

then $\mathrm{cay}_\alpha(\rho)$ represents a rotation through angle $\theta$ about the axis $u$. The inverse transformation

$$\mathrm{cay}_\alpha^{-1} : SO(3) \to \mathbb{R}^3$$

is given by

$$\mathrm{cay}_\alpha^{-1}(R) = \frac{2\alpha}{1 + \mathrm{tr}(R)}\ \mathrm{vec}(R - R^T), \tag{6}$$

where vec(·) denotes the inverse of the cross-product matrix operator, i.e., the axial vector associated with a skew-symmetric matrix. Throughout this work, rotational coordinates are represented using scaled Cayley parameters with scaling factor $\alpha = 5$, while translational coordinates are measured in Å. This scaling places rotational and translational fluctuations on comparable numerical scales and follows the convention adopted throughout the cgNA family of models [11, 12, 18, 15].

The base-pair coordinates are divided into intra-base-pair and inter-base-pair coordinate blocks. The intra-base-pair coordinate block is

$$x_i = (\tau_i, \xi_i) = \left\{ \mathrm{cay}_\alpha^{-1}(\Lambda_i), B_i^T(r_i^+ - r_i^-) \right\} \in \mathbb{R}^6, \tag{7}$$

where $\tau_i \in \mathbb{R}^3$ and $\xi_i \in \mathbb{R}^3$ denote the rotational and translational intra-base-pair coordinates, respectively. These correspond to the Curves+ intra-base-pair parameters buckle, propeller, opening, shear, stretch, and stagger.

The inter-base-pair coordinate block associated with the junction between base pairs $i$ and $i+1$ is

$$y_i = (u_i, v_i) = \left\{ \mathrm{cay}_\alpha^{-1}(\Gamma_i), J_i^T(g_{i+1} - g_i) \right\} \in \mathbb{R}^6, \tag{8}$$

where $u_i \in \mathbb{R}^3$ and $v_i \in \mathbb{R}^3$ denote the rotational and translational inter-base-pair coordinates, respectively. These correspond to the Curves+ inter-base-pair parameters tilt, roll, twist, shift, slide, and rise.

Phosphate coordinates are defined relative to the nucleobase, i.e., base-to-5′-phosphate. The phosphate coordinate block is

$$z_i^\pm = (\eta_i^\pm, w_i^\pm) = \left\{ \mathrm{cay}_\alpha^{-1}\left( (D_i^\pm)^T D_i^{p\pm} \right), (D_i^\pm)^T (r_i^{p\pm} - r_i^\pm) \right\} \in \mathbb{R}^6, \tag{9}$$

where $\eta_i^\pm \in \mathbb{R}^3$ and $w_i^\pm \in \mathbb{R}^3$ denote the rotational and translational phosphate coordinates, respectively, and the superscripts $+$ and $-$ refer to Watson- and Crick-strands.

For a blunt-ended duplex of length $N$, the complete cgNA+ internal-coordinate vector is

$$\Omega = \left( x_1, z_1^-, y_1, z_2^+, x_2, z_2^-, \ldots, y_{N-1}, z_N^+, x_N \right) \in \mathbb{R}^{24N-18}. \tag{10}$$

The vector $\Omega$ contains $6N$ intra-base-pair coordinates, $6N-6$ inter-base-pair coordinates, and $12N-12$ phosphate coordinates.

##### 2.3 Change of reading strand

The cgNA+ internal-coordinate representation depends on the choice of reading strand. A given duplex can be represented either by reading the Watson strand,

$$\mathcal{S} = X_1 X_2 \cdots X_N,$$

or, equivalently, by reading the complementary Crick strand,

$$\bar{\mathcal{S}} = \bar{X}_N \bar{X}_{N-1} \cdots \bar{X}_1.$$

These two descriptions correspond to the same physical duplex but yield different internal-coordinate vectors. They are related by a linear involutive transformation [15, 18],

$$\Omega(\bar{\mathcal{S}}) = E_N\, \Omega(\mathcal{S}), \qquad \mathcal{K}(\bar{\mathcal{S}}) = E_N\, \mathcal{K}(\mathcal{S})\, E_N, \tag{11}$$

where $\Omega$ denotes the cgNA+ internal-coordinate vector, $\mathcal{K}$ the corresponding stiffness matrix, and $E_N$ a length-dependent signed permutation matrix.

$$\mathbb{R}^{24N-18\times 24N-18} \ni E_N = \begin{bmatrix} & & & & E^{5'} \\ & & & E^{\text{int}} & \\ & & \iddots & & \\ & E^{\text{int}} & & & \\ E^{3'} & & & & \end{bmatrix}, \tag{12}$$

where $E^{5'}$, $E^{3'}$, and $E^{\text{int}}$ describe the transformations acting on the terminal 5′-end, terminal 3′-end, and interior coordinate blocks, respectively.

$$\mathbb{R}^{36\times 36} \ni E^{5'} = \begin{bmatrix} & & & & & E \\ & & & & I_6 & \\ & & & E & & \\ & & I_6 & & & \\ & E & & & & \\ I_6 & & & & & \end{bmatrix}, \tag{13}$$

$$E^{3'} = (E^{5'})^{-1} = (E^{5'})^T, \tag{14}$$

$$\mathbb{R}^{42\times 42} \ni E^{\text{int}} = \begin{bmatrix} & & & & & & I_6 \\ & & & & & E & \\ & & & & I_6 & & \\ & & & E & & & \\ & & I_6 & & & & \\ & E & & & & & \\ I_6 & & & & & & \end{bmatrix}, \tag{15}$$

where

$$E = \begin{bmatrix} -I_3 & 0 \\ 0 & I_3 \end{bmatrix},$$

changes the sign of the rotational coordinates while leaving the translational coordinates unchanged, and $I_3$ and $I_6$ denote the $3\times 3$ and $6\times 6$ identity matrices, respectively.

The matrix $E_N$ reverses the ordering of coordinate blocks and changes the sign of those coordinates whose orientation depends on the reading direction. It satisfies

$$E_N^{-1} = E_N^T = E_N.$$

For chemically symmetric duplexes such as dsDNA and dsRNA, the change-of-reading-strand transformation is equivalent to Crick–Watson strand exchange and can be used to reduce the number of independent sequence-dependent parameter blocks. This symmetry does not apply to DRH, since exchanging the strands changes the underlying molecular chemistry.

#### 2.4 cgNA+ internal coordinates to frames

The transformation from rigid-body frames to internal coordinates described in Section 2.2 is invertible. Given a cgNA+ configuration

$$\Omega = \big(x_1, z_1^-, y_1, z_2^+, x_2, \ldots, y_{N-1}, z_N^+, x_N\big) \in \mathbb{R}^{24N-18},$$

the corresponding base and phosphate frames can be reconstructed uniquely up to an arbitrary global rigid-body motion. Reconstruction proceeds in three stages. First, the base-pair frames $(B_i, g_i)$ are recovered recursively from the inter-base-pair coordinates $y_i = (u_i, v_i)$. Since the internal-coordinate representation is invariant under global rigid-body motions, an arbitrary reference frame may be chosen. Throughout this work, the reconstruction is initialised with

$$B_1 = I, \qquad g_1 = \mathbf{0}.$$

The remaining base-pair frames are then obtained by successively applying the relative rigid-body transformations encoded by the inter-base-pair coordinates.

Second, the Watson- and Crick-strand base frames are reconstructed from the intra-base-pair coordinates $x_i = (\eta_i, \xi_i)$, which specify the relative orientation and displacement of the two bases within each base pair.

Finally, the phosphate frames are recovered from the phosphate coordinates $z_i^{\pm} = (\tau_i^{\pm}, \zeta_i^{\pm})$, which define the orientation and position of each phosphate group relative to its associated nucleobase.

The reconstructed base and phosphate frames provide a complete coarse-grained representation of the duplex. When required, atomistic structures can be regenerated by embedding the ideal nucleobase and phosphate geometries described in Section 2.1 into the corresponding rigid-body frames. This procedure produces a full three-dimensional atomistic structure that is consistent with the cgNA+ configuration.

Consequently, the sequence-dependent equilibrium probability distribution predicted by cgNA+ can be mapped directly to an ensemble of atomistic structures. This enables the generation of representative conformations, structural ensembles, and derived observables directly from the coarse-grained model. The same reconstruction procedure is used by the cgNA+web server, which provides downloadable three-dimensional structures in standard atomistic PDB format generated from cgNA+ predictions.

### 3 cgNA+ model

#### 3.1 Estimation of oligomer-level statistics

For a given sequence, once each snapshot of the MD time-series (post H-bond filtering) has been transformed into internal coordinates, the oligomer-level statistics can be estimated by fitting a multivariate Gaussian distribution, $\rho_{\text{MD}}(\Omega; \mathcal{S})$ characterized by the groundstate $\hat{\Omega}(\mathcal{S})$ and inverse covariance $\mathcal{C}(\mathcal{S})$ or stiffness matrix $\mathcal{K}(\mathcal{S})$. For an MD trajectory $[\Omega^m(\mathcal{S})]_{m=1}^{M}$, where $m$ indexes the snapshots and $M \sim 10^6$ is the total number of configurations, the groundstate and covariance matrix are estimated as

$$\begin{aligned} \hat{\Omega}(\mathcal{S}) &= \frac{1}{M} \sum_{m=1}^{M} \Omega^m(\mathcal{S}), \\ C(\mathcal{S}) &= \frac{1}{M} \sum_{m=1}^{M} \left(\Omega^m(\mathcal{S}) - \hat{\Omega}(\mathcal{S})\right) \left(\Omega^m(\mathcal{S}) - \hat{\Omega}(\mathcal{S})\right)^T. \end{aligned} \tag{16}$$

For palindromic dsDNA and dsRNA sequences ($\mathcal{S} = \bar{\mathcal{S}}$), the complementary Crick–Watson symmetry is exploited to improve the statistical estimates by defining the symmetrized estimators

$$
\begin{aligned}
\hat{\Omega}_p(\mathcal{S}) &= \frac{1}{2}\left(\hat{\Omega} + E_N\hat{\Omega}\right), \\
\mathcal{H}(\mathcal{S}) &= \mathcal{C} + \hat{\Omega}\hat{\Omega}^T, \\
\mathcal{H}_p(\mathcal{S}) &= \frac{1}{2}\left(\mathcal{H} + E_N\mathcal{H}E_N\right), \\
\mathcal{C}_p(\mathcal{S}) &= \mathcal{H}_p - \hat{\Omega}_p\hat{\Omega}_p^T,
\end{aligned}
\tag{17}
$$

where $\mathcal{H}$ is the second moment and $(E_N)$ is the linear operator corresponding to the reading-strand transformation (see Section 2.3). Note that, unlike dsDNA, no complementary Watson–Crick symmetry can be exploited for DRH, even when $\mathcal{S} = \bar{\mathcal{S}}$, because $\mathcal{S}$ and $\bar{\mathcal{S}}$ represent chemically distinct molecules.

##### 3.2 cgNA+ inference

Given a dsNA sequence

$$\mathcal{S} = X_1 X_2 \cdots X_N$$

and a cgNA+ parameter set $\mathcal{P}$, the corresponding sequence-dependent stress-like vector $\sigma(\mathcal{P}, \mathcal{S})$ and stiffness matrix $\mathcal{K}(\mathcal{P}, \mathcal{S})$ are assembled from the local dimer-dependent parameter blocks. Because neighbouring dimer steps share internal-coordinate degrees of freedom, the local parameter blocks overlap. Their contributions are combined through the sparse Boolean reconstruction matrix

$$R_d \in \{0, 1\}^{(42N-12)\times(24N-18)},$$

yielding the oligomer-level

$$\mathcal{K}(\mathcal{P}, \mathcal{S}) = R_d^T \mathcal{K}_d R_d \qquad \text{and} \qquad \sigma(\mathcal{P}, \mathcal{S}) = R_d^T \sigma_d, \tag{18}$$

where

$$\mathcal{K}_d = \mathrm{diag}\left(\mathcal{K}^{5'X_1X_2}, \ldots, \mathcal{K}^{X_iX_{i+1}}, \ldots, \mathcal{K}^{X_{N-1}X_N3'}\right)$$

is the block-diagonal matrix of local dimer stiffness matrices, and

$$\sigma_d = \left(\sigma^{5'X_1X_2}, \ldots, \sigma^{X_iX_{i+1}}, \ldots, \sigma^{X_{N-1}X_N3'}\right)$$

is the corresponding concatenated stress-like vector.

The reconstruction matrix $R_d$ maps local dimer parameters onto the oligomer-level internal-coordinate vector. Its block structure is

$$
R_d = \begin{bmatrix}
I_{18} & & & & & & \cdots & \\
& I_{18} & & & & & & \\
& I_{18} & & & & & & \\
& & I_6 & & & & & \\
& & & I_{18} & & & & \\
& & & I_{18} & & & & \\
& & & & I_6 & & & \\
\vdots & & & & & & \ddots & \vdots \\
& & & & & & & I_{18}
\end{bmatrix},
\tag{19}
$$

where $I_k$ denotes the $k \times k$ identity matrix.

Because neighbouring dimer parameter blocks overlap through shared internal coordinates, this assembly process is not invertible. A schematic illustration of the reconstruction procedure is shown below in Figure S2.

The above mapping rule, which given a parameter set $\mathcal{P}$ and $\mathcal{S}$ returns banded matrix $\mathcal{K}(\mathcal{P}, \mathcal{S})$ and $\sigma(\mathcal{P}, \mathcal{S})$ can be succinctly written as:

$$\mathcal{R}(\mathcal{P}, \mathcal{S}) = (\sigma(\mathcal{P}, \mathcal{S}), \mathcal{K}(\mathcal{P}, \mathcal{S})). \tag{20}$$

For computation purposes, we have used the vectorised form of the parameter set denoted as $\mathcal{P}_{\text{vec}} \in \mathbb{R}^L$, where $L$ is number of independent entries in parameter set. An analogous reconstruction rule in form of $\mathcal{P}_{\text{vec}}$ can be written as

$$\mathcal{R}_{\text{vec}}(\mathcal{P}, \mathcal{S}) = \mathcal{R}_{\mathcal{P}}(\mathcal{S})\mathcal{P}_{\text{vec}} = (\sigma(\mathcal{P}, \mathcal{S}), \mathcal{K}(\mathcal{P}, \mathcal{S})). \tag{21}$$

where $\mathcal{R}_{\mathcal{P}} \in N + N^2 \times L$ is parameter reconstruction matrix which maps $\mathcal{P}_{\text{vec}}$ to the entries of $\mathcal{K}(\mathcal{P}, \mathcal{S})$ and $\sigma(\mathcal{P}, \mathcal{S})$.

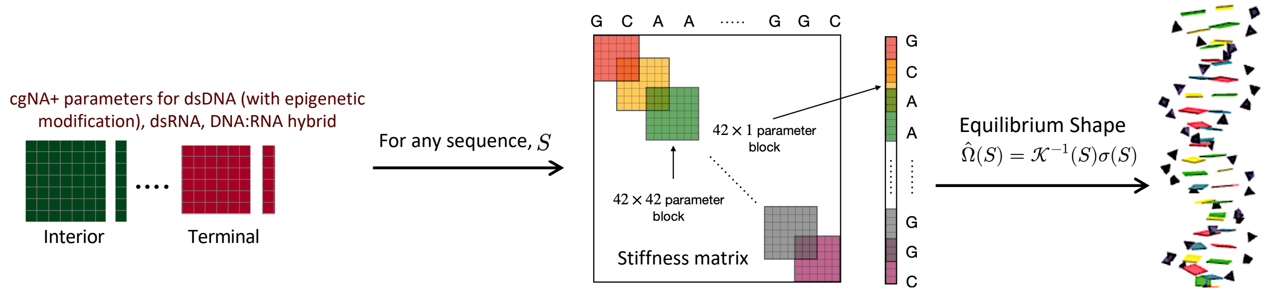

Figure S2: Assembly of sequence-dependent stress-like vector $\sigma$ and banded stiffness matrix $\mathcal{K}$ from local dimer parameter blocks. Terminal 5′-, interior-, and terminal 3′-dimer blocks are shown in different colours. Each matrix cell has dimension $6 \times 6$, while each vector cell has dimension $6 \times 1$.

##### 3.3 Quadratic free-energy approximation

The cgNA+ model approximates the conformational free-energy landscape of a dsNA oligomer by a quadratic function of the internal-coordinate vector $\Omega \in \mathbb{R}^{24N-18}$. The oligomer free energy is assumed to be the sum of local contributions associated with successive dimer steps [18, 15, 11],

$$U(\Omega; \mathcal{S}) \approx \sum_{i=1}^{N-1} U_i(\Omega_i) = \sum_{i=1}^{N-1} \left[ \frac{1}{2}(\Omega_i - \widehat{\Omega}_i)^T \mathcal{K}_i (\Omega_i - \widehat{\Omega}_i) + c_i \right], \tag{22}$$

where $\Omega_i$ denotes the local internal-coordinate vector associated with dimer step $i$, $\widehat{\Omega}_i$ and $\mathcal{K}_i$ are the corresponding equilibrium configuration and stiffness matrix, respectively, and $c_i$ is an additive constant independent of $\Omega_i$. Expanding the local quadratic forms and introducing the local equilibrium-shape (force) vectors

$$\sigma_i = \mathcal{K}_i \widehat{\Omega}_i,$$

the reconstruction procedure described in the previous subsection yields the global free-energy function

$$U(\Omega; \mathcal{S}) = \frac{1}{2}\Omega^T \mathcal{K}(\mathcal{S})\Omega - \sigma(\mathcal{S})^T \Omega + c, \tag{23}$$

where

$$\mathcal{K}(\mathcal{S}) = R_d^T \mathcal{K}_d R_d, \qquad \sigma(\mathcal{S}) = R_d^T \sigma_d.$$

The equilibrium (groundstate) conformation is obtained by minimizing $U(\Omega; \mathcal{S})$. Since $\mathcal{K}(\mathcal{S})$ is symmetric positive definite, the minimizer is unique and satisfies

$$\nabla_\Omega U = \mathcal{K}(\mathcal{S})\Omega - \sigma(\mathcal{S}) = 0.$$

Hence,

$$\widehat{\Omega}(\mathcal{S}) = \mathcal{K}(\mathcal{S})^{-1}\sigma(\mathcal{S}). \tag{24}$$

Thus, the equilibrium-shape vector $\sigma$ and stiffness matrix $\mathcal{K}$ completely determine the sequence-dependent equilibrium conformation and the associated quadratic free-energy landscape.

##### 3.4 Computation of new cgNA+ parameter sets

Parameter sets for dsDNA, dsRNA, and DRH are currently available. A detailed description of parameter estimation is beyond the scope of this presentation, which instead focuses on how the model can be used with existing parameter sets [19]. Nevertheless, in our experience, new parameter sets corresponding to different training libraries and MD protocols can be obtained efficiently by using an existing parameter set as the initial guess for the gradient-flow optimisation described in the main text. In contrast, finding an initial guess for the computation of the first parameter set was quite delicate [15].

The iterative optimisation in Eq. 7 (main text) is equivalent to a quasi-Newton scheme in which the Jacobian matrix is held fixed over all iterations. In this context, the Jacobian is precisely the Fisher information matrix, and is evaluated at the theoretically ideal convergence point where the model and MD distributions agree. As the iterations approach this theoretical point, the difference between the fixed and actual local Jacobian vanishes, and the scheme effectively behaves like an exact Newton scheme, with a much more rapid rate of convergence than standard gradient descent. We note that our scheme employs a pseudo-inverse instead of a usual inverse of the Jacobian due to the fact that it has some zero modes, which arise from symmetries in the model. We note also that analytical expressions are available for the Jacobian, due to the Gaussian form of the distributions involved. A detailed analysis of this scheme will be given elsewhere [15].

The Fisher information [20] can be defined as the second centred moment of $\log \rho(x; \theta)$ conditional to parameter $\mathbb{R}^N \ni \theta = \{\mu, \mathcal{K}\},\ N > 1$,

$$[\mathbb{F}(\theta)]_{ij} = -\mathbb{E}\left[\frac{\partial^2}{\partial\theta_i \partial\theta_j}(\log \rho(x;\theta))\bigg|\theta\right] \tag{25}$$

where $\rho(x;\theta)$ is a pdf conditional on parameter $\theta \in \mathbb{R}^N$, and $\mu$ and $\mathcal{K}$ are the mean and inverse covariance matrix. Furthermore, for two parametric pdfs that are close in parametric space, say $\rho(x;\theta')$ and $\rho(x;\theta)$ where $\theta = \theta' + \delta\theta,\ \theta,\ \theta',\ \delta\theta \in \mathbb{R}^N$ and $\delta\theta << 1$, there exists a relation between KL divergence (defined in Eq. 8 in the main paper) and Fisher information,

$$\mathbb{F}(\theta) = -\int_\Omega \rho(x,\theta') \frac{\partial^2}{\partial\theta'^2} \log\big(\rho(x,\theta')\big) = \frac{\partial^2}{\partial\theta^2} D_{\mathrm{KL}}(\rho(x,\theta'), \rho(x,\theta))\big|_{\theta=\theta'}. \tag{26}$$

Now, Taylor expansion at $\theta = \theta'$ gives,

$$D_{\mathrm{KL}}(\rho(x;\theta'),\rho(x;\theta)) = \frac{1}{2}\delta\theta.\mathbb{F}(\theta)\delta\theta + \mathcal{O}(|\delta\theta^3|) = D_{\mathrm{KL}}(\rho(x;\theta),\rho(x;\theta')). \tag{27}$$

The training of cgNA+ model requires computation of the KL divergence between the observed Gaussian for each oligomer in the training library, $\rho(w;\theta_{\mathrm{MD}}(\mathcal{S}))$ and banded Gaussian predicted by cgNA+ reconstruction, $\rho(w;\theta_{\mathcal{P}_{\mathrm{vec}}}(S,\mathcal{P}))$. Now, Eq. 27 can be approximated as,

$$D_{\mathrm{KL}}(\rho(w;\theta_{\mathcal{P}_{\mathrm{vec}}}),\rho(w;\theta_{\mathrm{MD}})) \approx \frac{1}{2}\theta_{\mathcal{P}_{\mathrm{vec}}}.\mathbb{F}(\theta_{\mathrm{MD}})\theta_{\mathcal{P}_{\mathrm{vec}}} - \theta_{\mathcal{P}_{\mathrm{vec}}}.\mathbb{F}(\theta_{\mathrm{MD}})\theta_{\mathrm{MD}} + \frac{1}{2}\theta_{\mathrm{MD}}.\mathbb{F}(\theta_{\mathrm{MD}})\theta_{\mathrm{MD}}. \tag{28}$$

Using the relation $\theta_{\mathcal{P}_{\mathrm{vec}}}(\mathcal{P},\mathcal{S}) = R_{\mathcal{P}}(\mathcal{S})\mathcal{P}_{\mathrm{vec}}$ (Eq. 21) in Eq. 28 gives,

$$\begin{aligned} D_{\mathrm{KL}}(\rho(w;\theta_{\mathcal{P}_{\mathrm{vec}}}),\rho(w;\theta_{\mathrm{MD}})) \approx &\frac{1}{2}\mathcal{P}_{\mathrm{vec}}.R^T_{\mathcal{P}}(\mathcal{S})\mathbb{F}(\theta_{\mathrm{MD}})R_{\mathcal{P}}(\mathcal{S})\mathcal{P}_{\mathrm{vec}} \\ &- \frac{1}{2}\mathcal{P}_{\mathrm{vec}}.R^T_{\mathcal{P}}(\mathcal{S})\mathbb{F}(\theta_{\mathrm{MD}})\theta_{\mathrm{MD}} + \frac{1}{2}\theta_{\mathrm{MD}}.\mathbb{F}(\theta_{\mathrm{MD}})\theta_{\mathrm{MD}}. \end{aligned} \tag{29}$$

Now, using linear change of variable $\mathbb{F}_{(\mathcal{P},\mathcal{S})}(\theta_{\mathrm{MD}}) = R^T_{\mathcal{P}}(\mathcal{S})\mathbb{F}(\theta_{\mathrm{MD}})R_{\mathcal{P}}(\mathcal{S})$ in above equation gives

$$\begin{aligned} D_{\mathrm{KL}}(\rho(w;\theta_{\mathcal{P}_{\mathrm{vec}}}),\rho(w;\theta_{\mathrm{MD}})) \approx &\frac{1}{2}\mathcal{P}_{\mathrm{vec}}.\mathbb{F}_{(\mathcal{P},\mathcal{S})}(\theta_{\mathrm{MD}})\mathcal{P}_{\mathrm{vec}} \\ &- \frac{1}{2}\mathcal{P}_{\mathrm{vec}}.R^T_{\mathcal{P}}(\mathcal{S})\mathbb{F}(\theta_{\mathrm{MD}})\theta_{\mathrm{MD}} + \frac{1}{2}\theta_{\mathrm{MD}}.\mathbb{F}(\theta_{\mathrm{MD}})\theta_{\mathrm{MD}}. \end{aligned} \tag{30}$$

Now, in order to find the best-fit parameter set, the following equation needs to be minimized,

$$\begin{aligned} \sum_{i=1}^{L} D_{\mathrm{KL}}(\rho(w;\theta_{\mathcal{P}_{\mathrm{vec}}}(\mathcal{S}_i)),\rho(w;\theta_{\mathrm{MD}}(\mathcal{S}_i))) \approx &\sum_{i=1}^{L}\frac{1}{2}\mathcal{P}_{\mathrm{vec}}.\mathbb{F}_{(\mathcal{P},\mathcal{S}_i)}(\theta_{\mathrm{MD}})\mathcal{P}_{\mathrm{vec}} \\ &- \sum_{i=1}^{L}\frac{1}{2}\mathcal{P}_{\mathrm{vec}}.R^T_{\mathcal{P}}(\mathcal{S}_i)\mathbb{F}(\theta_{\mathrm{MD}})\theta_{\mathrm{MD}} + \sum_{i=1}^{L}\frac{1}{2}\theta_{\mathrm{MD}}.\mathbb{F}(\theta_{\mathrm{MD}})\theta_{\mathrm{MD}}. \end{aligned} \tag{31}$$

Thus, differentiating Eq. 31 with respect to $\mathcal{P}_{\mathrm{vec}}$ gives,

$$\sum_{i=1}^{L}\mathbb{F}_{(\mathcal{P},\mathcal{S}_i)}(\theta_{\mathrm{MD}})\mathcal{P}_{\mathrm{vec}} - \sum_{i=1}^{L} R^T_{\mathcal{P}}(\mathcal{S}_i)\mathbb{F}(\theta_{\mathrm{MD}})\theta_{\mathrm{MD}} = 0, \tag{32}$$

which can be solved using the least squares method. Eq. 32 can be rewritten as

$$\begin{aligned} &\mathbb{F}_{(\mathcal{P},\mathrm{Lb})}\mathcal{P}_{\mathrm{vec}} = B \\ &\text{where } \mathbb{F}_{(\mathcal{P},\mathrm{Lb})} = \sum_{i=1}^{L}\mathbb{F}_{(\mathcal{P},\mathcal{S}_i)}(\theta_{\mathrm{MD}}),\quad B = \sum_{i=1}^{L} R^T_{\mathcal{P}}(\mathcal{S}_i)\mathbb{F}(\theta_{\mathrm{MD}})\theta_{\mathrm{MD}}. \end{aligned} \tag{33}$$

However, the matrix $\mathbb{F}_{(\mathcal{P},\mathrm{Lb})}$ is not invertible due to non-injectivity of the reconstruction rule (section 3.2). Therefore, to find the least squares solution for Eq. 33, Moore-Penrose pseudo-inverse has been used to obtain the initial guess as

$$\mathcal{P}^{\mathrm{lsq}}_{\mathrm{vec}} = \mathbb{F}^{\dagger}_{(\mathcal{P},\mathrm{Lb})}B, \tag{34}$$

where $\mathbb{F}^{\dagger}_{(\mathcal{P},\mathrm{Lb})}$ is pseudo-inverse of $\mathbb{F}_{(\mathcal{P},\mathrm{Lb})}$.

### 4 Supplementary tables

Table S6: The cgNA+ prediction error in terms of Mahalanobis distance and KL divergence per degree of freedom for training and test sequences. The sequence-variability (which quantifies variation over sequence) is obtained by computing the average pair-wise symmetric Mahalanobis distance (Mahal) and KL divergence (KLD) between all the training sequences across dsNAs. Corresponding sequences are provided in Tables S1 and S2. MDNA and HDNA denote the sequences containing methylated and hydroxymethylated $C_pG$ steps, respectively.

| Training sequences | | | | | | | | | | |
|---|---|---|---|---|---|---|---|---|---|---|
| | dsDNA | | dsRNA | | DRH | | MDNA | | HDNA | |
| Index | Mahal | KLD | Mahal | KLD | Mahal | KLD | Mahal | KLD | Mahal | KLD |
| 1 | 0.0018 | 0.0240 | 0.0010 | 0.0058 | 0.0019 | 0.0239 | 0.0021 | 0.0329 | 0.0020 | 0.0322 |
| 2 | 0.0025 | 0.0439 | 0.0011 | 0.0064 | 0.0019 | 0.0255 | 0.0020 | 0.0266 | 0.0019 | 0.0268 |
| 3 | 0.0020 | 0.0302 | 0.0013 | 0.0070 | 0.0017 | 0.0171 | 0.0021 | 0.0261 | 0.0020 | 0.0245 |
| 4 | 0.0016 | 0.0267 | 0.0011 | 0.0083 | 0.0015 | 0.0166 | 0.0016 | 0.0211 | 0.0017 | 0.0357 |
| 5 | 0.0021 | 0.0289 | 0.0012 | 0.0081 | 0.0017 | 0.0226 | 0.0019 | 0.0260 | 0.0019 | 0.0256 |
| 6 | 0.0025 | 0.0368 | 0.0013 | 0.0063 | 0.0018 | 0.0213 | 0.0017 | 0.0290 | 0.0018 | 0.0291 |
| 7 | 0.0021 | 0.0353 | 0.0012 | 0.0070 | 0.0018 | 0.0210 | 0.0028 | 0.0329 | 0.0028 | 0.0322 |
| 8 | 0.0017 | 0.0266 | 0.0011 | 0.0071 | 0.0016 | 0.0162 | 0.0026 | 0.0281 | 0.0027 | 0.0304 |
| 9 | 0.0022 | 0.0328 | 0.0013 | 0.0080 | 0.0019 | 0.0216 | 0.0024 | 0.0342 | 0.0025 | 0.0352 |
| 10 | 0.0020 | 0.0276 | 0.0011 | 0.0074 | 0.0017 | 0.0209 | 0.0033 | 0.0402 | 0.0031 | 0.0387 |
| 11 | 0.0020 | 0.0342 | 0.0013 | 0.0095 | 0.0015 | 0.0168 | 0.0026 | 0.0329 | 0.0025 | 0.0307 |
| 12 | 0.0020 | 0.0322 | 0.0013 | 0.0067 | 0.0018 | 0.0220 | 0.0028 | 0.0350 | 0.0029 | 0.0360 |
| 13 | 0.0018 | 0.0297 | 0.0014 | 0.0101 | 0.0017 | 0.0215 | | | | |
| 14 | 0.0016 | 0.0282 | 0.0014 | 0.0092 | 0.0019 | 0.0219 | | | | |
| 15 | 0.0023 | 0.0344 | 0.0014 | 0.0101 | 0.0028 | 0.0395 | | | | |
| 16 | 0.0017 | 0.0296 | 0.0013 | 0.0076 | 0.0047 | 0.0532 | | | | |
| **Average** | 0.0020 | 0.0313 | 0.0012 | 0.0078 | 0.0020 | 0.0239 | 0.0023 | 0.0304 | 0.0023 | 0.0314 |
| Test sequences | | | | | | | | | | |
| Index | Mahal | KLD | Mahal | KLD | Mahal | KLD | Mahal | KLD | Mahal | KLD |
| 1 | 0.0026 | 0.0357 | 0.0015 | 0.0087 | 0.0032 | 0.0555 | 0.0025 | 0.0339 | 0.0026 | 0.0349 |
| 2 | 0.0037 | 0.0291 | 0.0014 | 0.0095 | 0.0021 | 0.0198 | 0.0033 | 0.0389 | 0.0032 | 0.0377 |
| 3 | 0.0034 | 0.0483 | 0.0021 | 0.0100 | 0.0021 | 0.0213 | 0.0022 | 0.0295 | 0.0024 | 0.0297 |
| 4 | 0.0027 | 0.0306 | 0.0018 | 0.0101 | 0.0027 | 0.0250 | 0.0023 | 0.0306 | 0.0022 | 0.0299 |
| 5 | 0.0026 | 0.0291 | 0.0015 | 0.0070 | 0.0022 | 0.0202 | 0.0025 | 0.0356 | 0.0023 | 0.0333 |
| 6 | 0.0022 | 0.0285 | 0.0011 | 0.0092 | 0.0018 | 0.0192 | 0.0027 | 0.0324 | 0.0027 | 0.0319 |
| 7 | 0.0019 | 0.0283 | 0.0010 | 0.0076 | 0.0019 | 0.0227 | 0.0019 | 0.0244 | 0.0018 | 0.0226 |
| 8 | 0.0024 | 0.0254 | 0.0016 | 0.0061 | | | 0.0025 | 0.0281 | 0.0028 | 0.0316 |
| 9 | 0.0027 | 0.0297 | | | | | 0.0024 | 0.0262 | 0.0023 | 0.0266 |
| 10 | 0.0016 | 0.1449 | | | | | | | | |
| **Average** | 0.0027 | 0.0316 | 0.0015 | 0.0085 | 0.0023 | 0.0262 | 0.0025 | 0.0311 | 0.0025 | 0.0309 |
| ***sequence-variability*** | 0.0245 | 0.4395 | 0.0177 | 0.2185 | 0.0209 | 0.3273 | 0.0211 | 0.3378 | 0.0214 | 0.3449 |

### 5 Supplementary figures

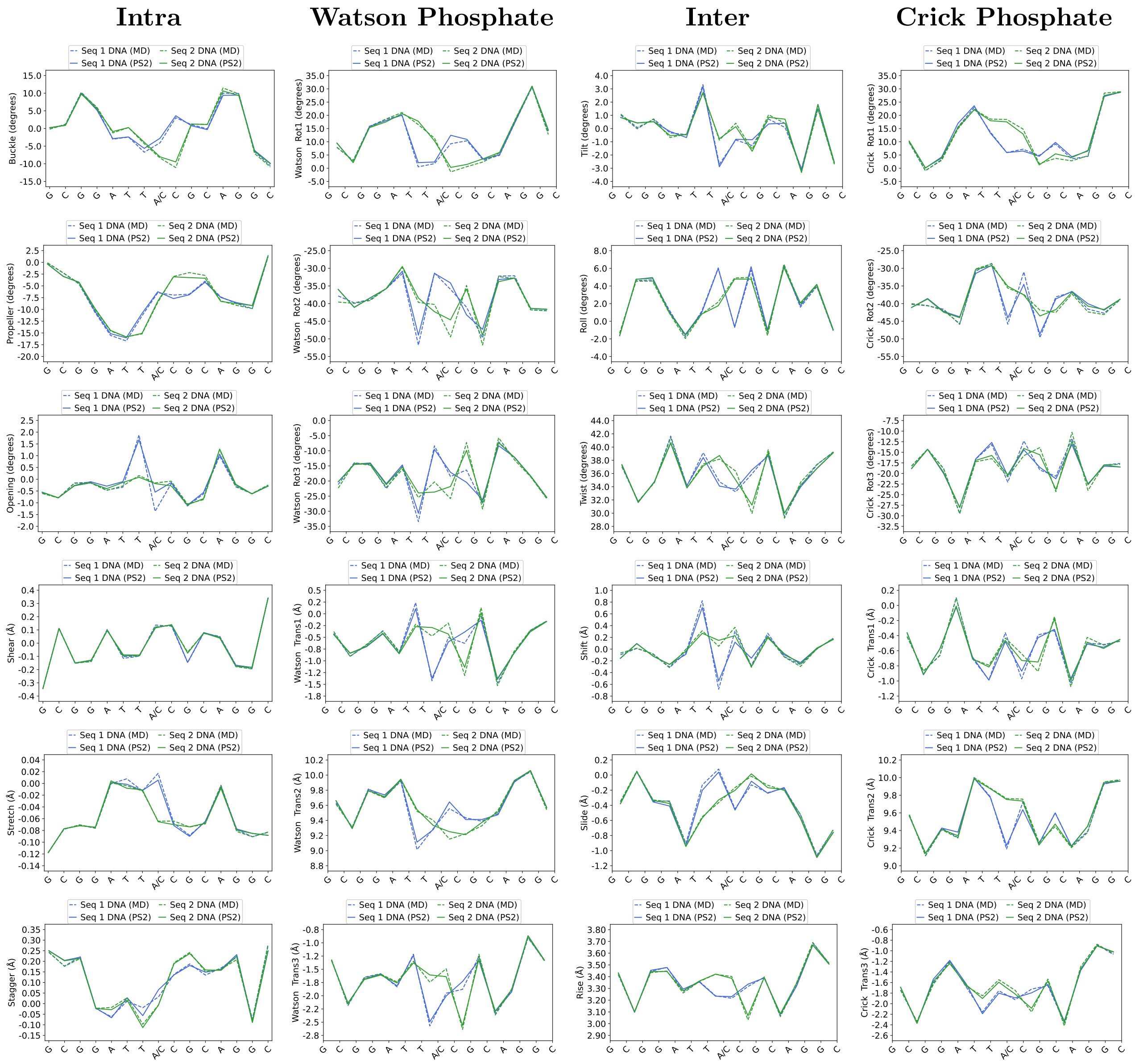

Figure S3: Comparison of cgNA+ predictions (solid line) with MD observations (dashed line) for two dsDNA sequences differing at only one position. cgNA+ model predictions are labelled as PS2 (the name of the corresponding parameter set), following the convention used in analogous data on the `https://cgdnaweb.epfl.ch` website.

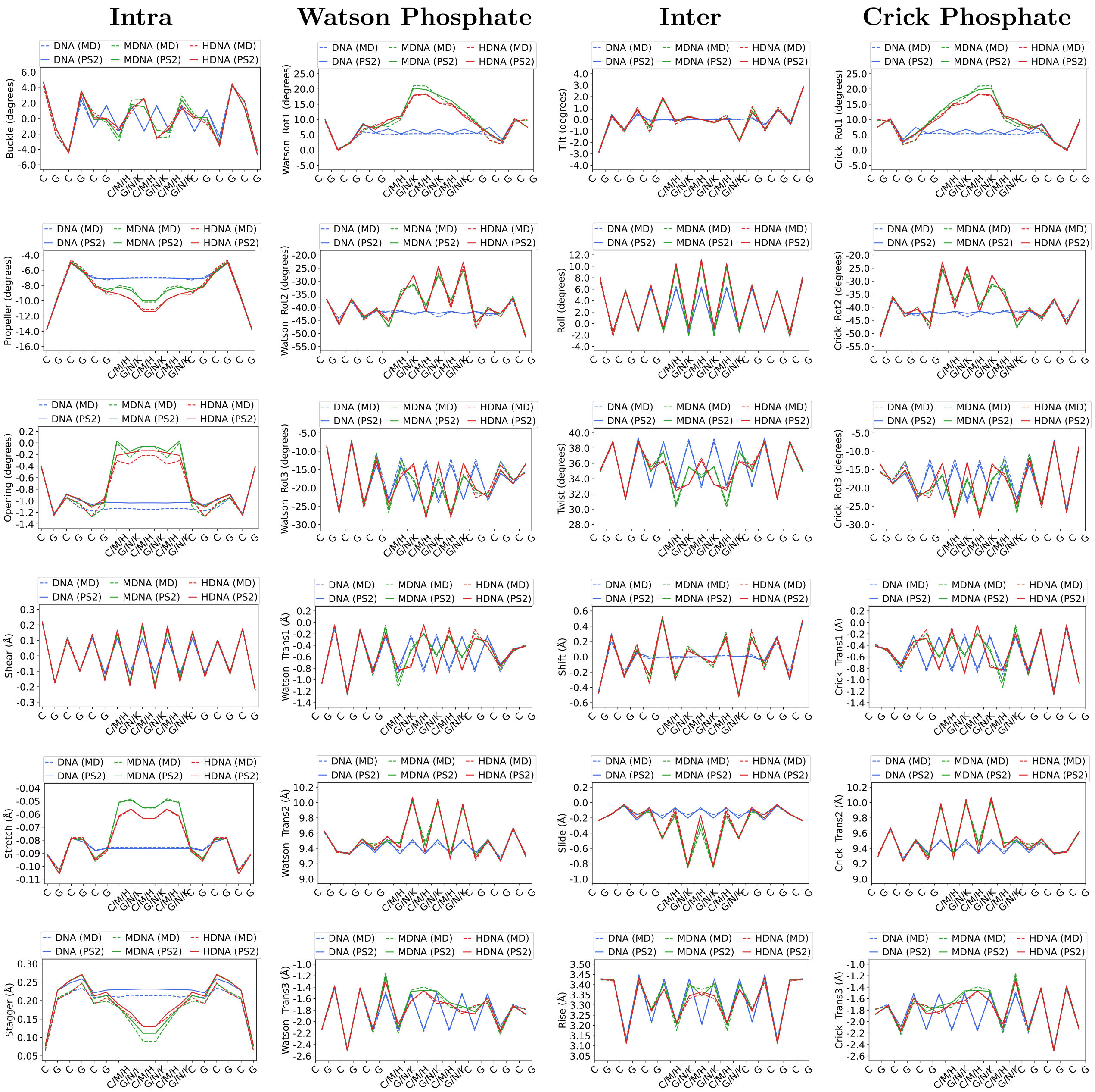

Figure S4: Comparison of cgNA+ predictions (solid line) with MD observations (dashed line) for three sequences $(CG)_9$ and its epigenetically methylated ($(CG)_3(MN)_3(CG)_3$) and hydroxymethylated version ($(CG)_3(HK)_3(CG)_3$). cgNA+ model predictions are labelled as PS2 (the name of the corresponding parameter set), following the convention used in analogous data on the `https://cgdnaweb.epfl.ch` website.

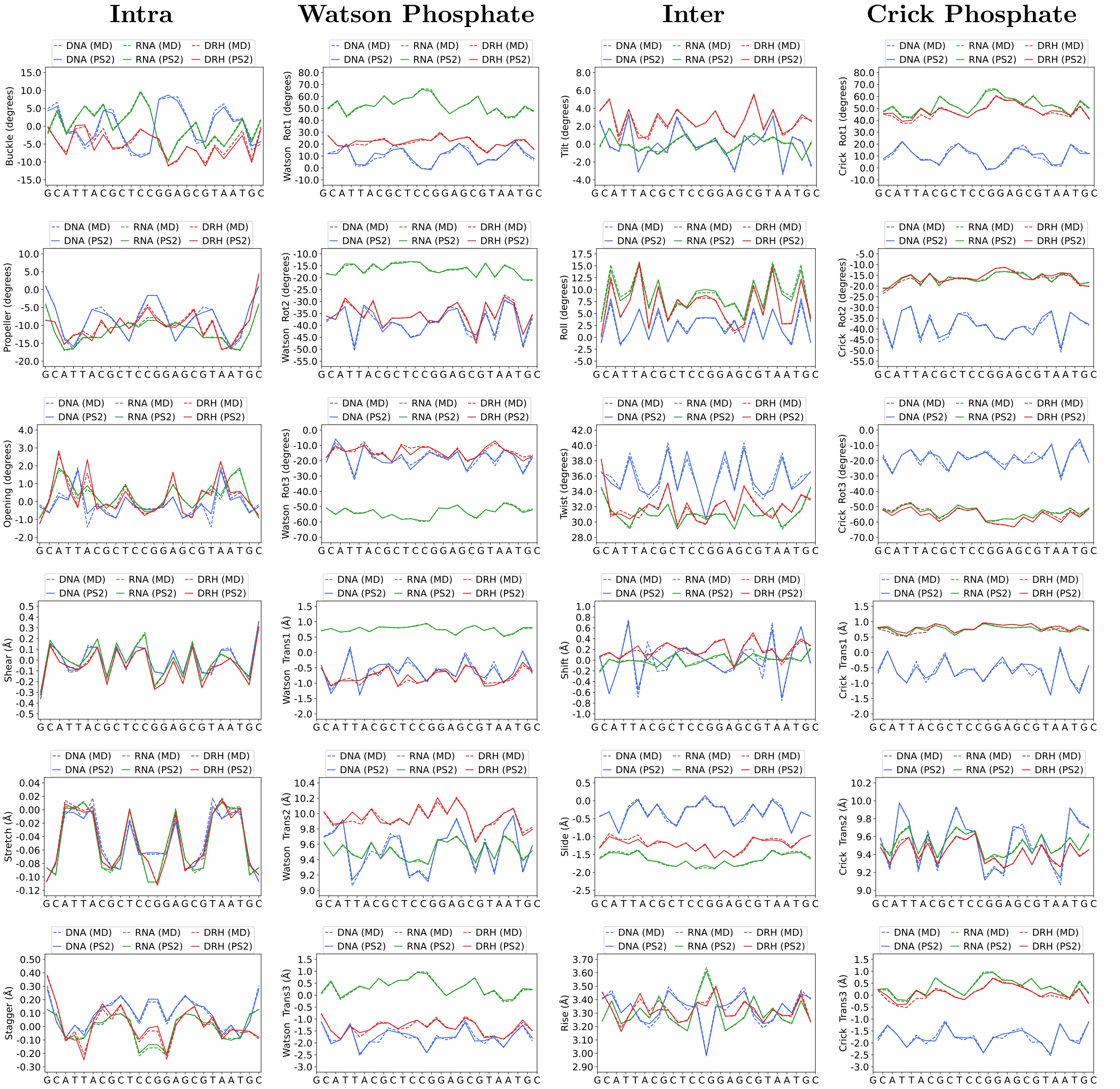

Figure S5: Comparison of cgNA+ predictions (solid line) with MD observations (dashed line) for a sequence in dsDNA, dsRNA and DNA:RNA hybrid sequences. cgNA+ model predictions are labelled as PS2 (the name of the corresponding parameter set), following the convention used in analogous data on the https://cgdnaweb.epfl.ch website.

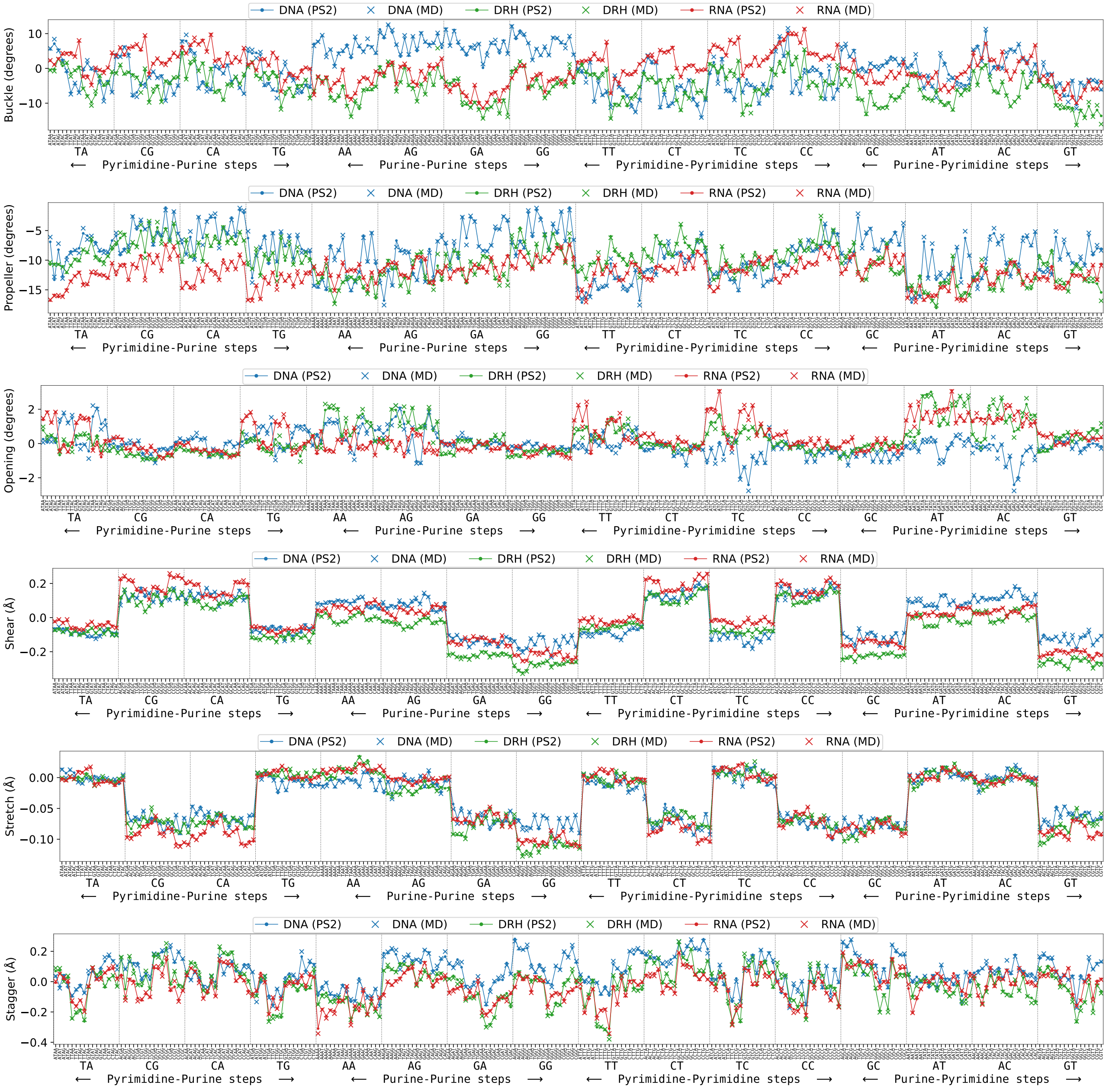

Figure S6: Comparison of intra base-pair (corresponding to first base-pair in dimer step) coordinates of the central dimer steps in all tetramer flanking contexts across dsNAs. Coordinates observed in MD simulations and cgNA+ predictions are plotted as × and •, respectively. For better visualization, a line plot is overlaid along the • symbols. cgNA+ predictions are referred to as PS2 as in the `https://cgdnaweb.epfl.ch` website.

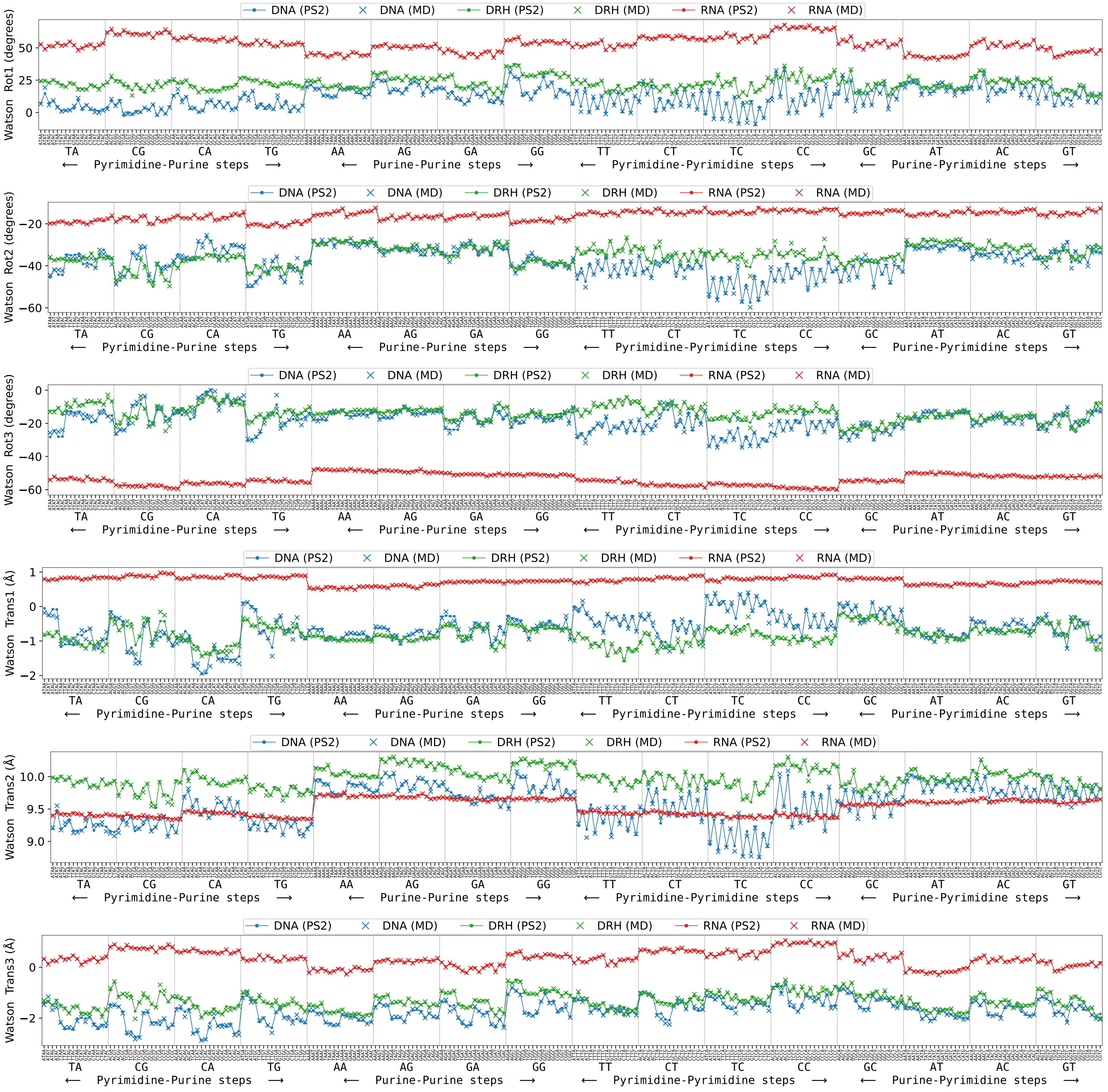

Figure S7: Comparison of Watson phosphate coordinates of the central dimer steps in all tetramer flanking contexts across dsNAs. Coordinates observed in MD simulations and cgNA+ predictions are plotted as × and •, respectively. For better visualization, a line plot is overlaid along the • symbols. cgNA+ predictions are referred to as PS2 as in the `https://cgdnaweb.epfl.ch` website.

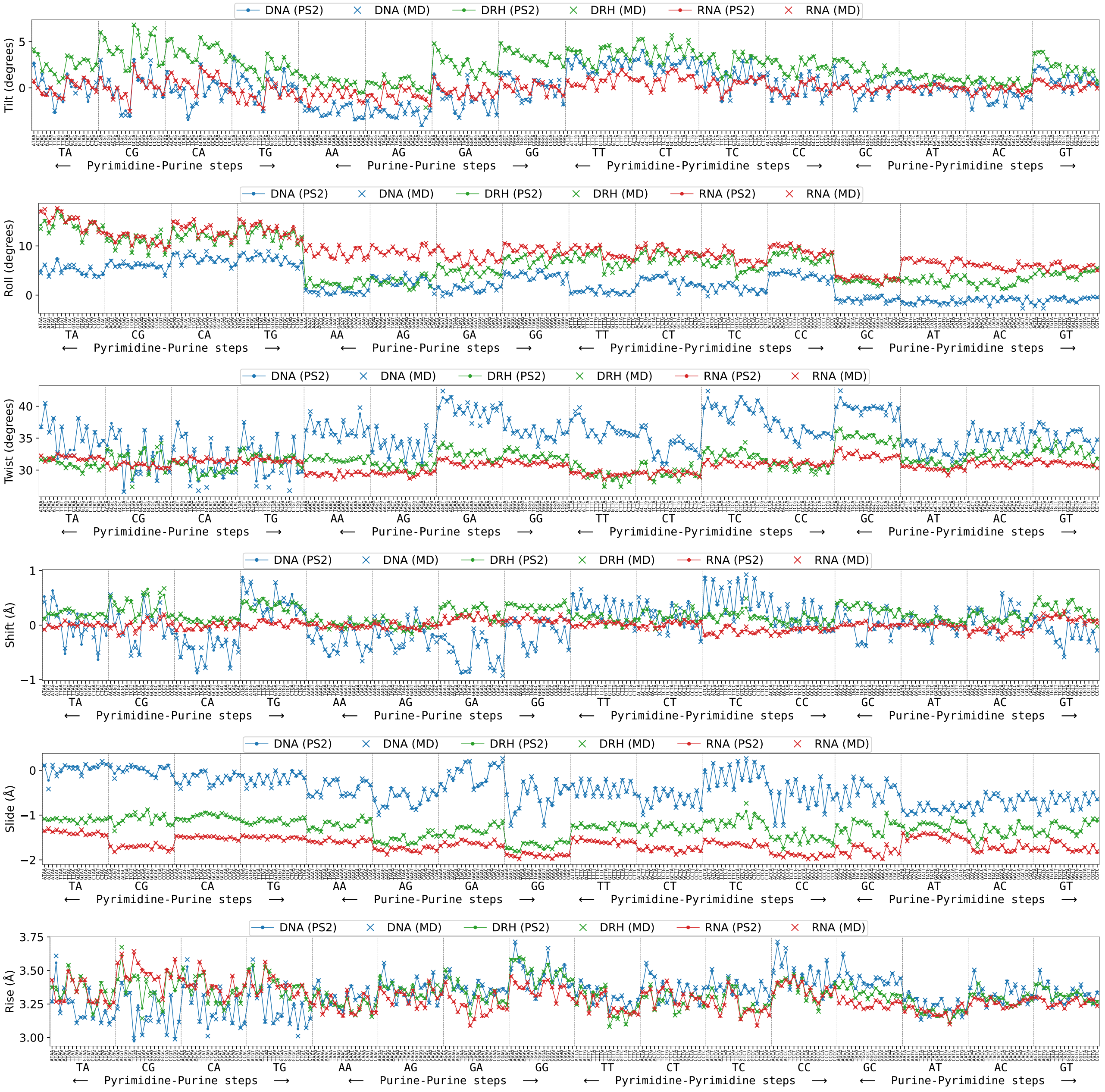

Figure S8: Comparison of inter base-pair step coordinates of the central dimer steps in all tetramer flanking contexts across dsNAs. Coordinates observed in MD simulations and cgNA+ predictions are plotted as × and •, respectively. For better visualization, a line plot is overlaid along the • symbols. cgNA+ predictions are referred to as PS2 as in the `https://cgdnaweb.epfl.ch` website.

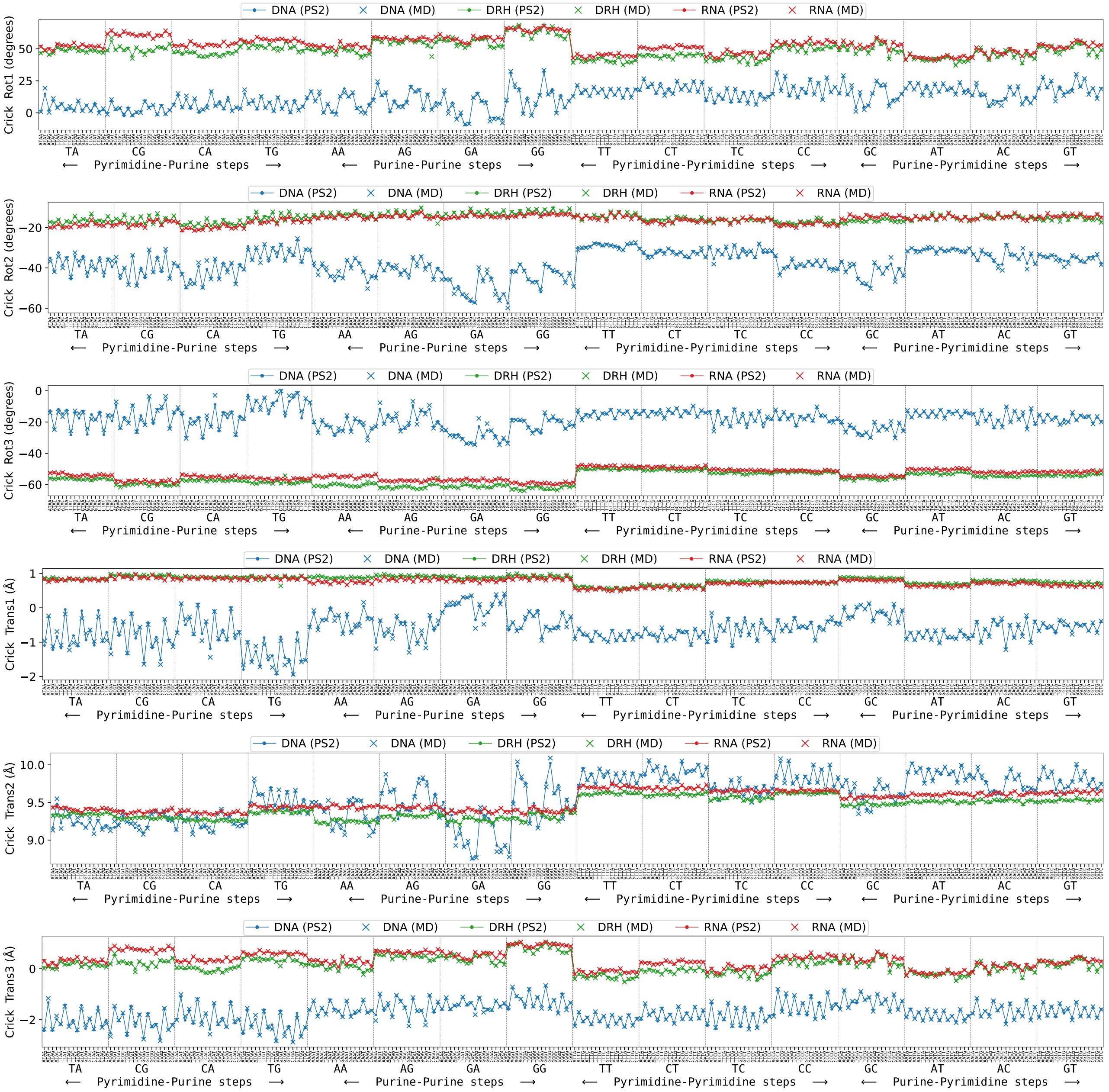

Figure S9: Comparison of Crick phosphate coordinates of the central dimer steps in all tetramer flanking contexts across dsNAs. Coordinates observed in MD simulations and cgNA+ predictions are plotted as × and •, respectively. For better visualization, a line plot is overlaid along the • symbols. cgNA+ predictions are referred to as PS2 as in the `https://cgdnaweb.epfl.ch` website.

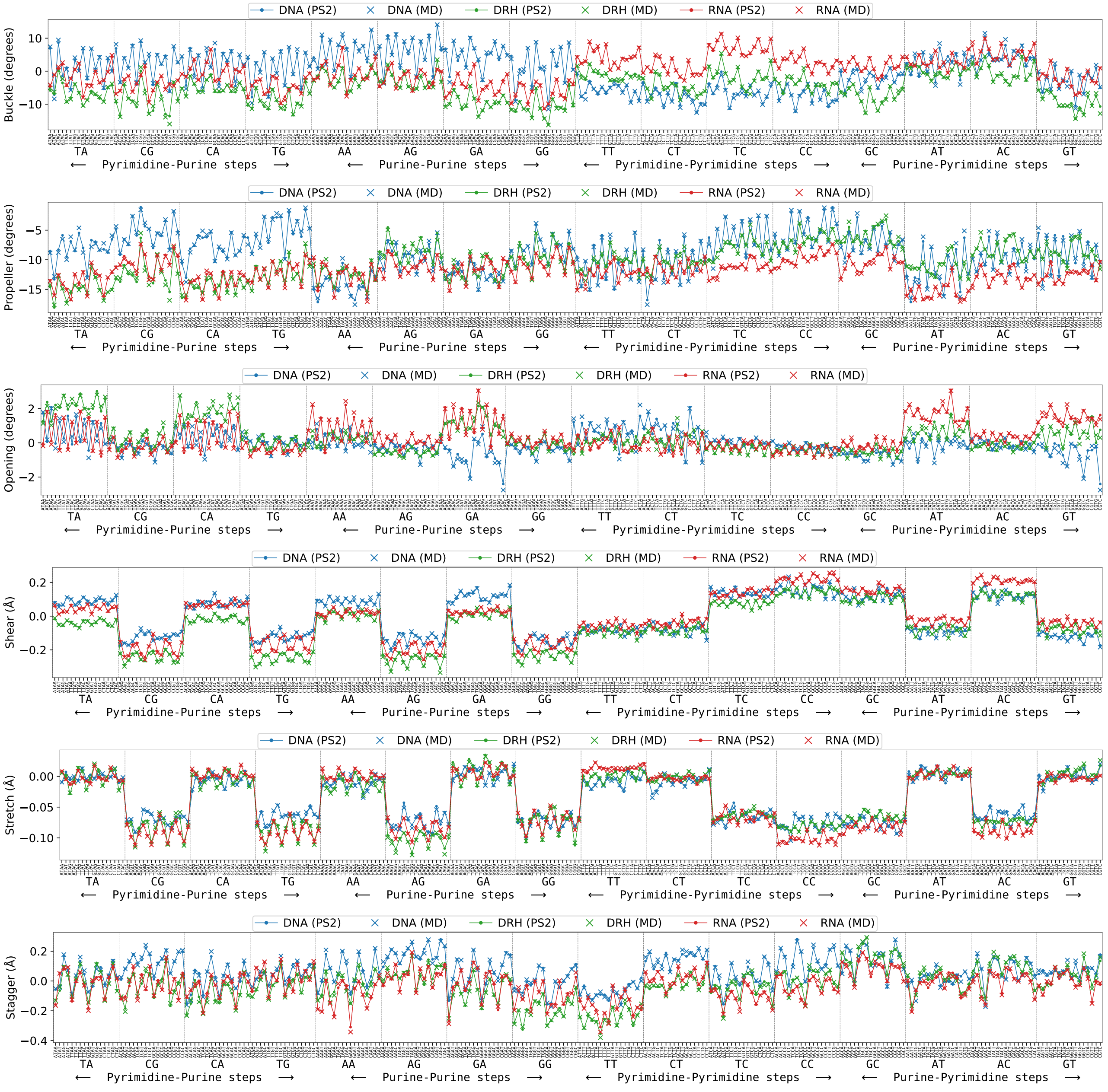

Figure S10: Comparison of intra base-pair (corresponding to second base-pair in dimer step) coordinates of the central dimer steps in all tetramer flanking contexts across dsNAs. Coordinates observed in MD simulations and cgNA+ predictions are plotted as × and •, respectively. For better visualization, a line plot is overlaid along the • symbols. cgNA+ predictions are referred to as PS2 as in the `https://cgdnaweb.epfl.ch` website.

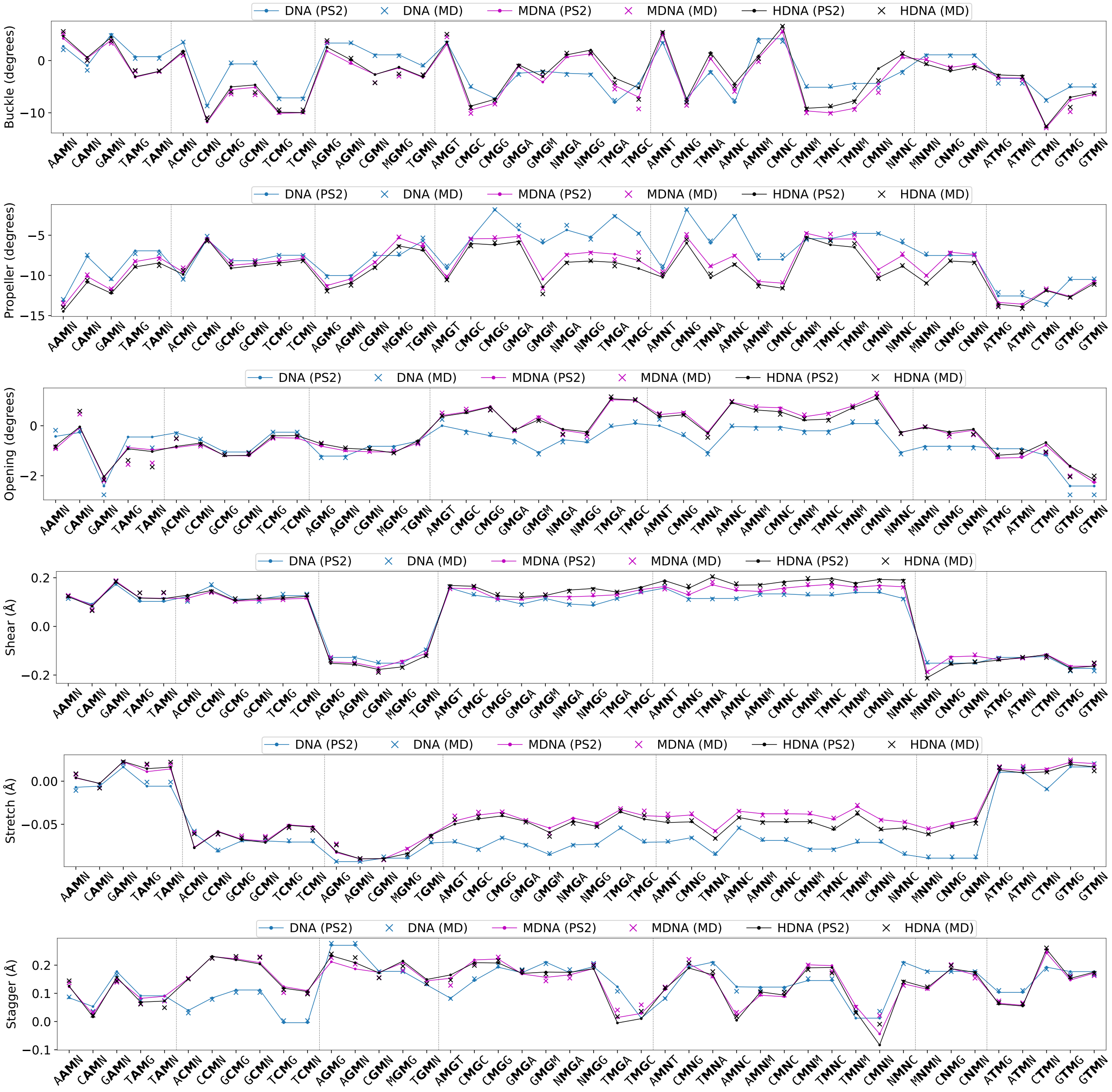

Figure S11: Comparison of intra base-pair (coordinates corresponding to the first base-pair in dimer step) coordinates of the central dimer steps in tetramer flanking contexts. All independent tetramers with methylated or hydroxymethylated Cytosine are considered. M and H denotes the methylated C and hydroxymethylated C while N and K denotes the G paired with methylated C and hydroxymethylated C. Coordinates observed in MD simulations and cgNA+ predictions are plotted as × and •, respectively. For better visualization, a line plot is overlaid along the • symbols. cgNA+ predictions are referred to as PS2 as in the `https://cgdnaweb.epfl.ch` website.

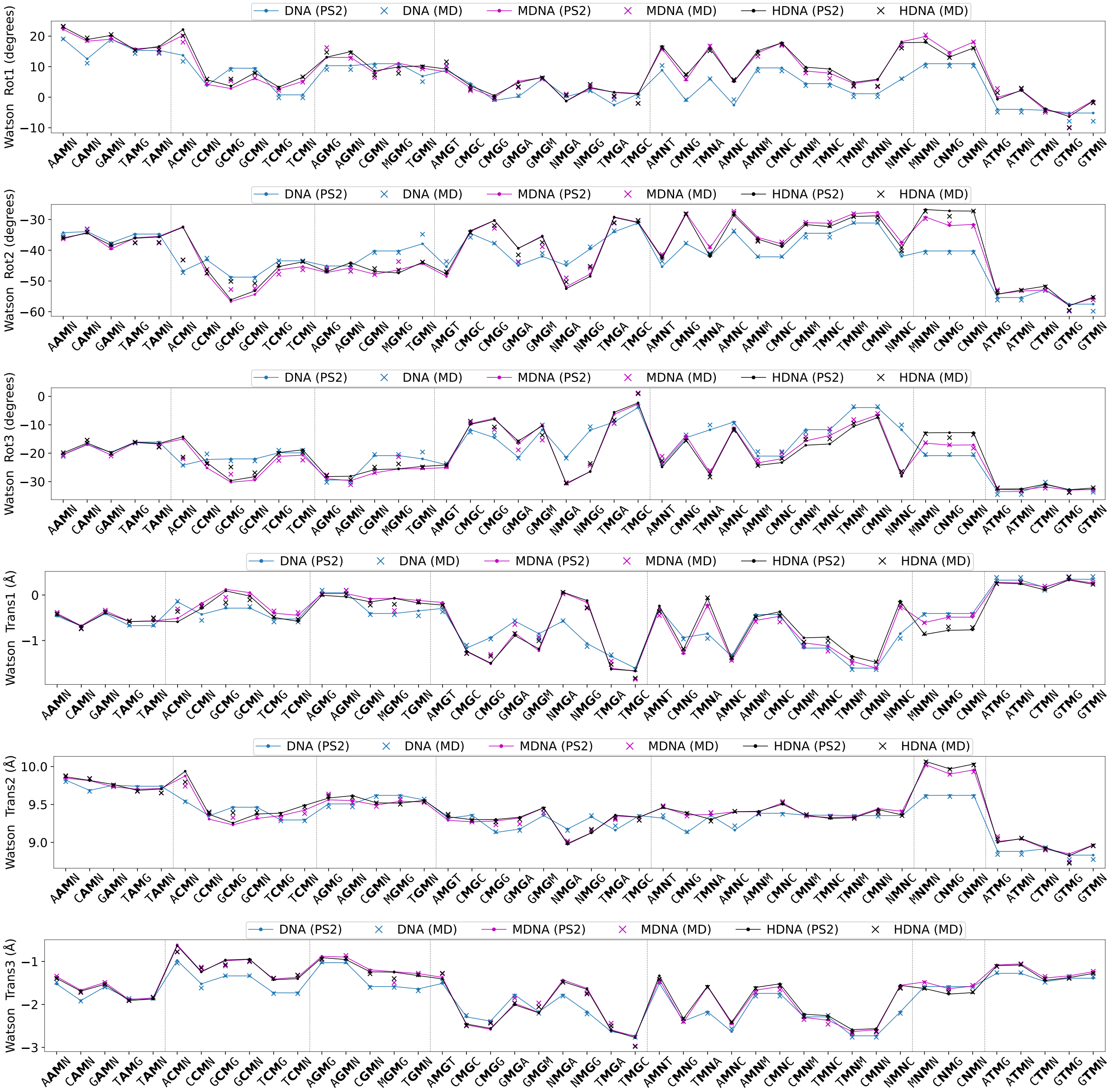

Figure S12: Comparison of Watson phosphate coordinates of the central dimer steps in tetramer flanking contexts. All independent tetramers with methylated or hydroxymethylated Cytosine are considered. M and H denotes the methylated C and hydroxymethylated C while N and K denotes the G paired with methylated C and hydroxymethylated C. Coordinates observed in MD simulations and cgNA+ predictions are plotted as × and •, respectively. For better visualization, a line plot is overlaid along the • symbols. cgNA+ predictions are referred to as PS2 as in the `https://cgdnaweb.epfl.ch` website.

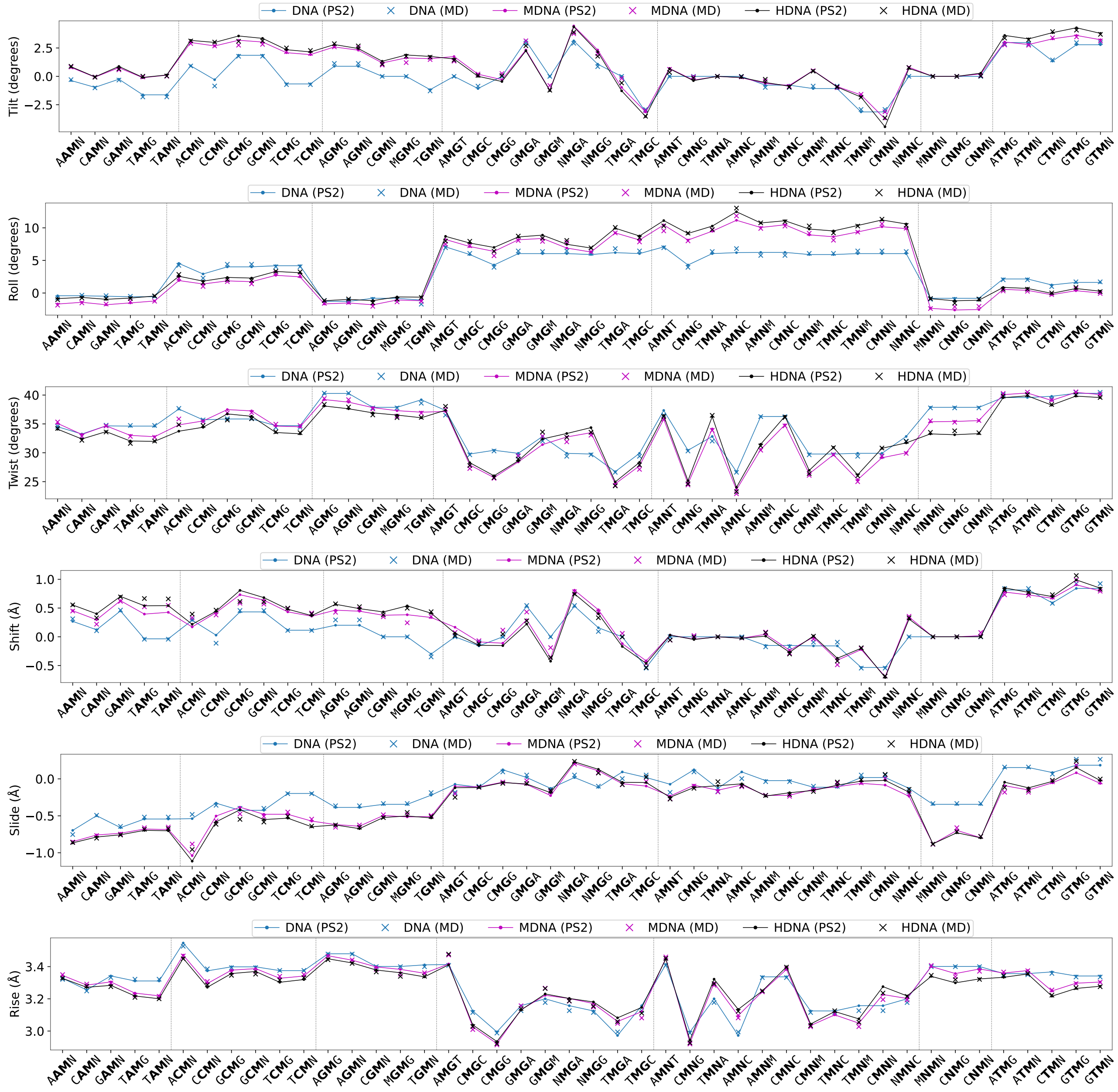

Figure S13: Comparison of inter base-pair step coordinates of the central dimer steps in tetramer flanking contexts. All independent tetramers with methylated or hydroxymethylated Cytosine are considered. M and H denotes the methylated C and hydroxymethylated C while N and K denotes the G paired with methylated C and hydroxymethylated C. Coordinates observed in MD simulations and cgNA+ predictions are plotted as × and •, respectively. For better visualization, a line plot is overlaid along the • symbols. cgNA+ predictions are referred to as PS2 as in the `https://cgdnaweb.epfl.ch` website.

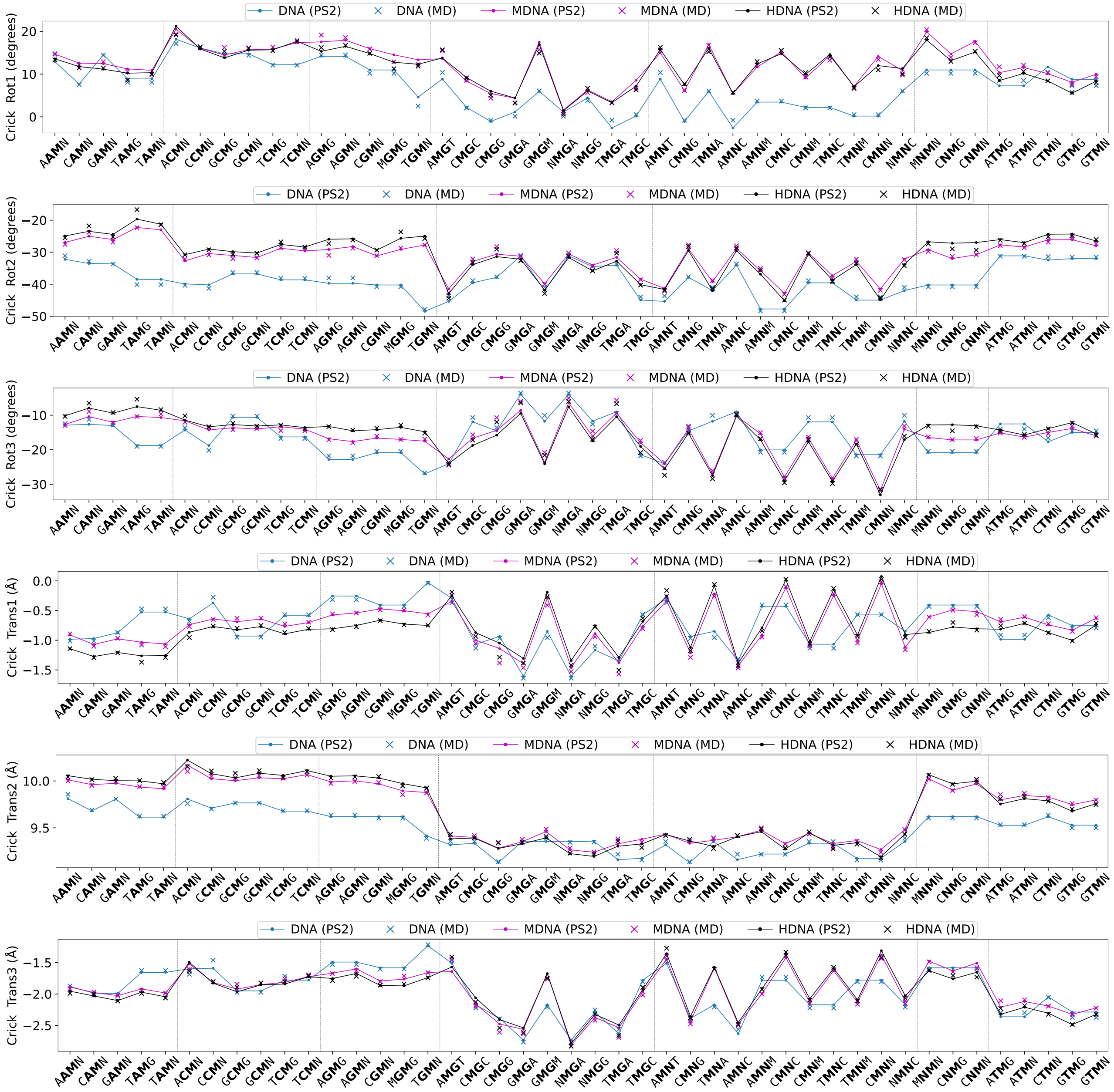

Figure S14: Comparison of Crick phosphate coordinates of the central dimer steps in tetramer flanking contexts. All independent tetramers with methylated or hydroxymethylated Cytosine are considered. M and H denotes the methylated C and hydroxymethylated C while N and K denotes the G paired with methylated C and hydroxymethylated C. Coordinates observed in MD simulations and cgNA+ predictions are plotted as × and •, respectively. For better visualization, a line plot is overlaid along the • symbols. cgNA+ predictions are referred to as PS2 as in the `https://cgdnaweb.epfl.ch` website.

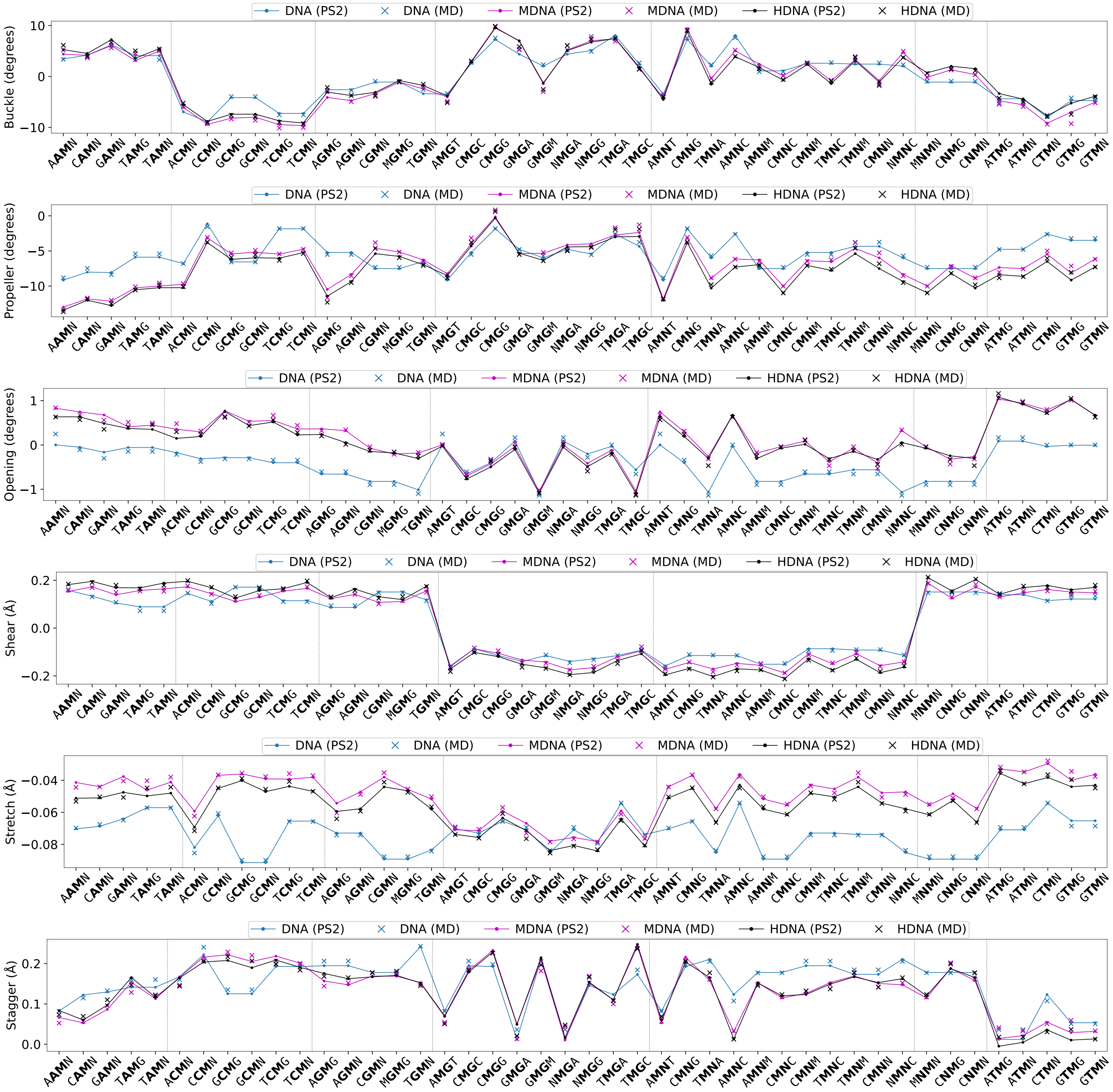

Figure S15: Comparison of intra base-pair (corresponding to second base-pair in dimer step) coordinates of the central dimer steps in tetramer flanking contexts. All independent tetramers with methylated or hydroxymethylated Cytosine are considered. M and H denotes the methylated C and hydroxymethylated C while N and K denotes the G paired with methylated C and hydroxymethylated C. Coordinates observed in MD simulations and cgNA+ predictions are plotted as × and •, respectively. For better visualization, a line plot is overlaid along the • symbols. cgNA+ predictions are referred to as PS2 as in the `https://cgdnaweb.epfl.ch` website.

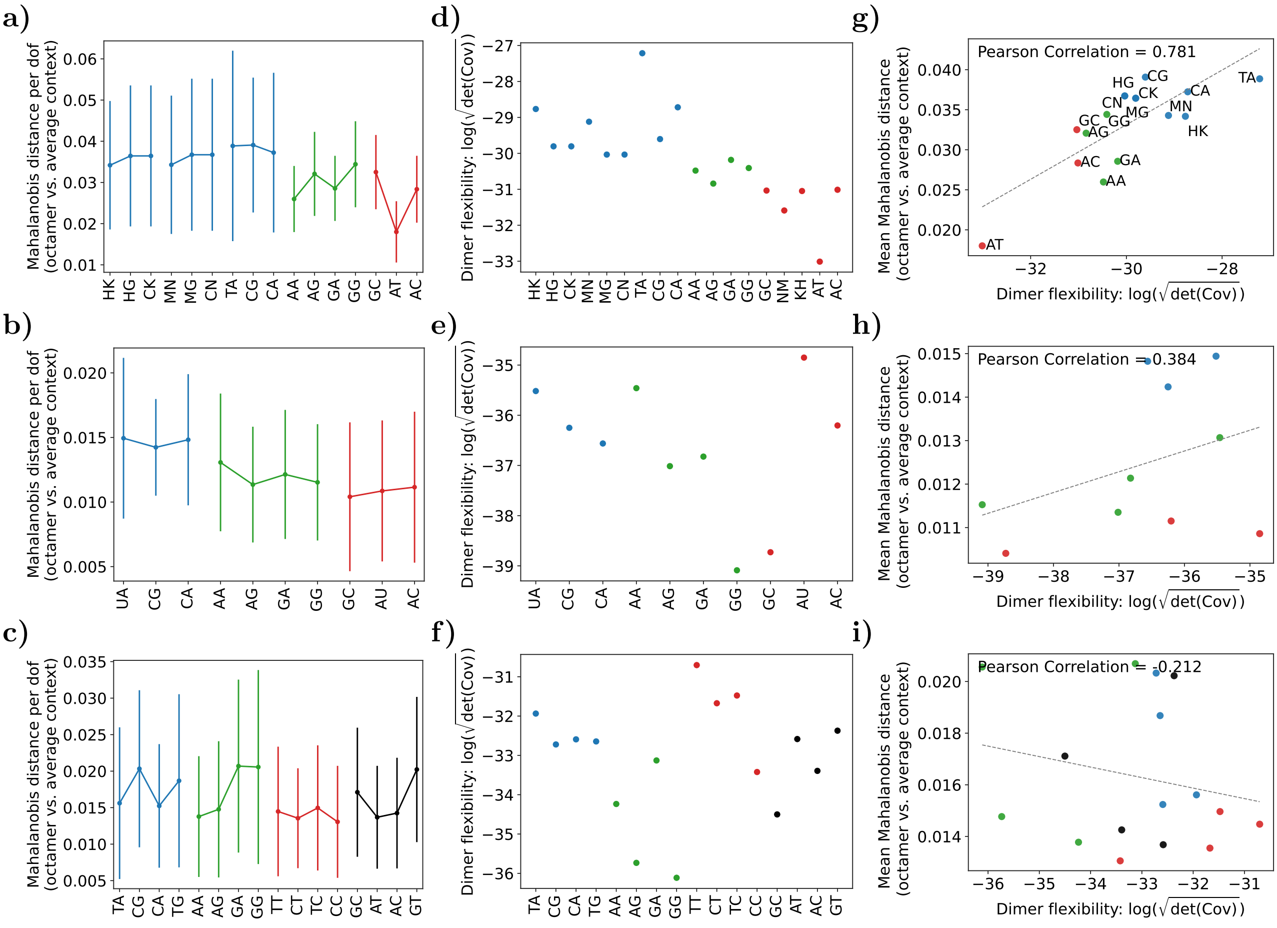

Figure S16: **(a–c)** Deviation of the dimer groundstate, or equilibrium shape, in a specific octamer context relative to its equilibrium shape in the average context. The difference is quantified using the symmetric Mahalanobis distance, and the error bars denote the standard deviation across all octamer contexts. **(d–f)** Configurational volume (in log scale) of the dimer in the average octamer context. For fluctuations in $\mathbb{R}^N$, the configurational volume is defined as $C_{vol} = \sqrt{\det(\mathrm{Cov})}$ [21]. Here the dimer junction fluctuations are in $\mathbb{R}^{30}$ and the unit of $C_{vol}$ is $\mathring{A}^{15} \cdot (\mathrm{rad}/5)^{15}$. **(g–i)** Pearson correlation between the deviation of the dimer groundstate, or equilibrium shape, in a specific octamer context relative to the average context, and the logarithm of the configurational volume.

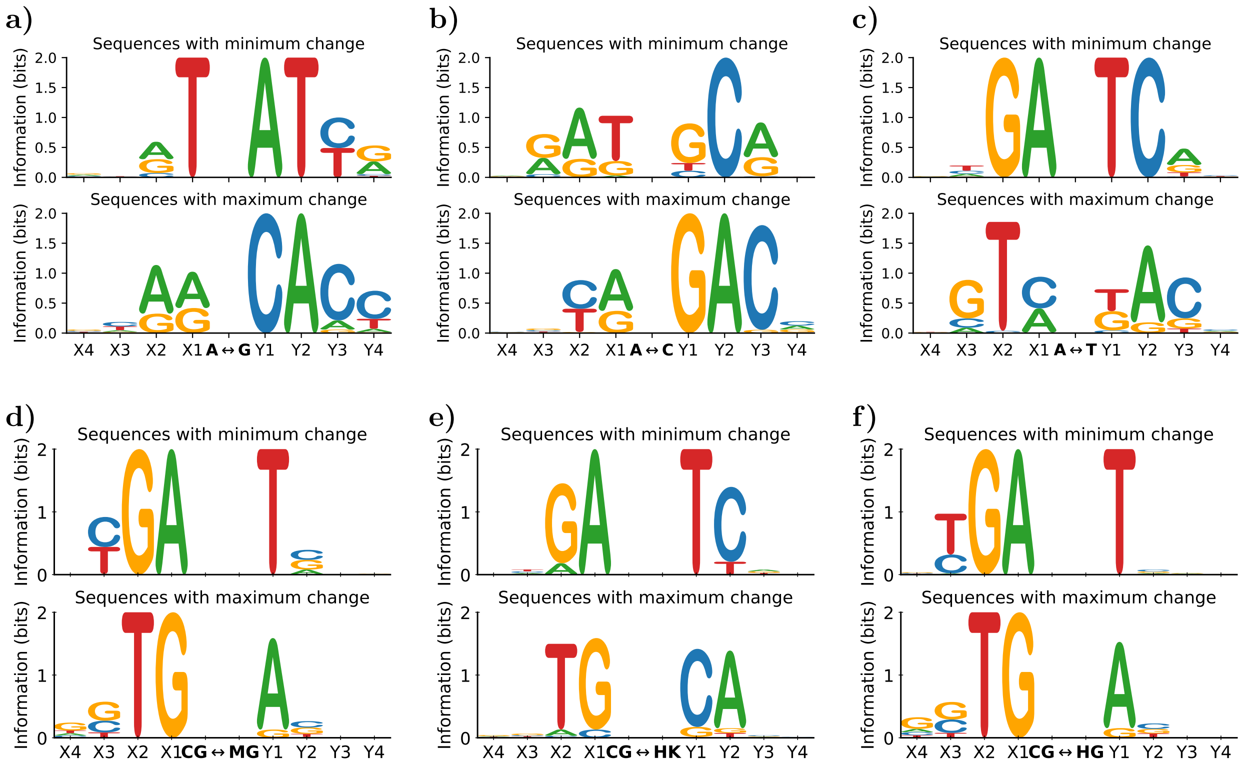

Figure S17: **(a)–(c)** Sequence logos for flanking contexts that least and most change the groundstate on various SNPs at $5^{\text{th}}$ position. **(d)–(f)** Sequence logos to highlight flanking contexts that least and most influence the change in groundstate upon epigenetic modification of central $C_pG$ step. Statistics are derived from all tetramer flanking contexts on both sides of the central SNP position or central $C_pG$ steps embedded in a 21/22mer: $GCGTCGX_4X_3X_2X_1$—central position —$Y_1Y_2Y_3Y_4GTCGGC$. The x-axis is base index with information content at that index on the y-axis. Information content in $X_j$ and $Y_j$ for the largest and smallest (beyond three standard deviations from the mean) changes in groundstate is plotted on the ordinate.

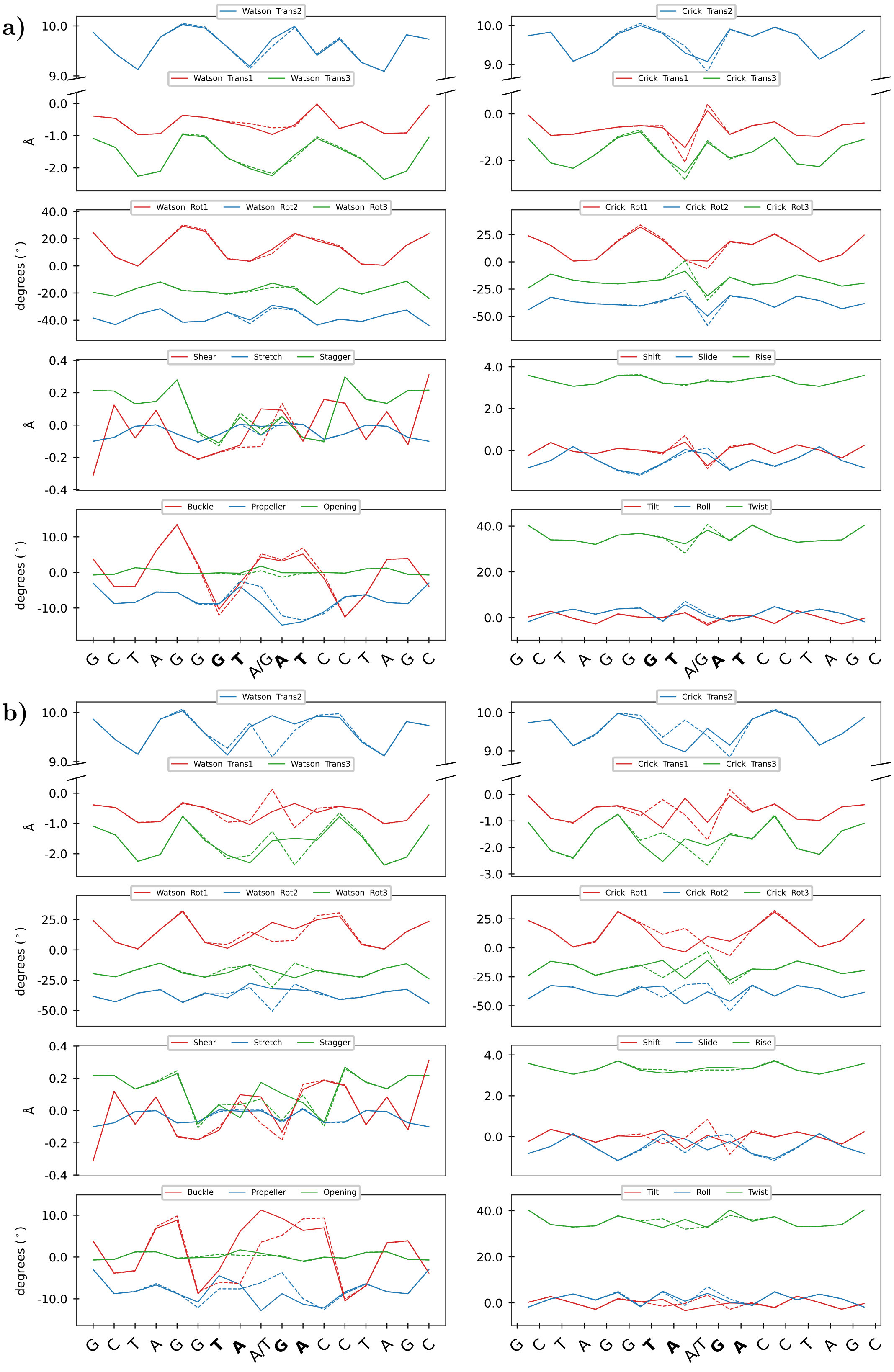

Figure S18: cgNA+ predicted equilibrium shape for two sets of sequences: **(a)** GC-TAGG**GT**A/G**AT**CCTAGC and **(b)** GCTAGG**TA**A/T**GA**CCTAGC each containing point mutations and differing only in the two base pairs flanking the mutation site on either side. These two examples visually illustrate the changes in the groundstate induced by point mutations and flanking sequence context.

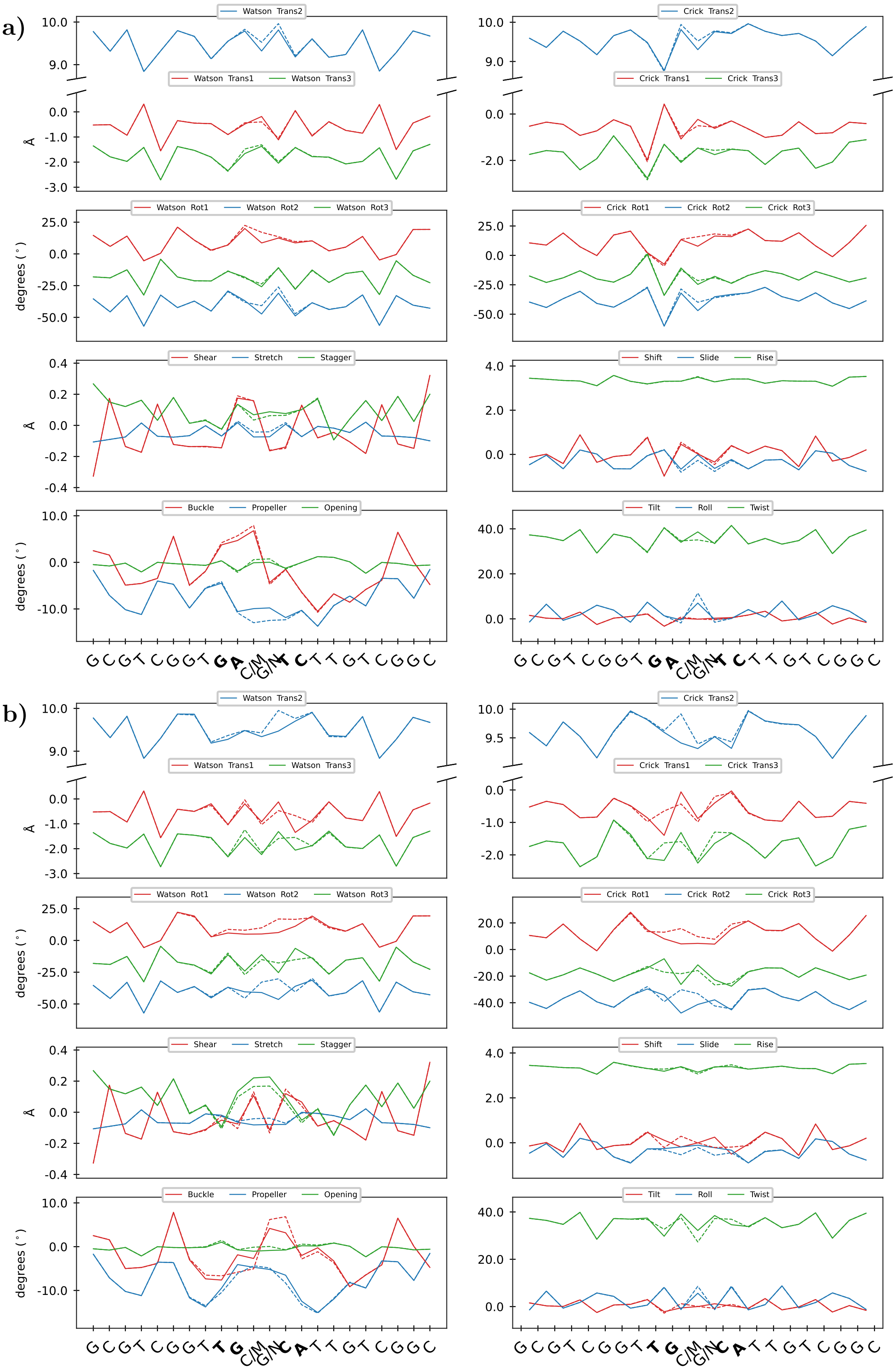

Figure S19: The groundstate for two sequences which only differ by immediate hexamer flanking context (underlined) to the central $C_pG$ step, **(a)** GCGTCGGTGA**CG**TCTTGTCGGC and **(b)** GCGTCGGTTG**CG**CATTGTCGGC to visualize the change in groundstate on the symmetric methylation of the central $C_pG$ step (in bold) to $M_pN$ step.

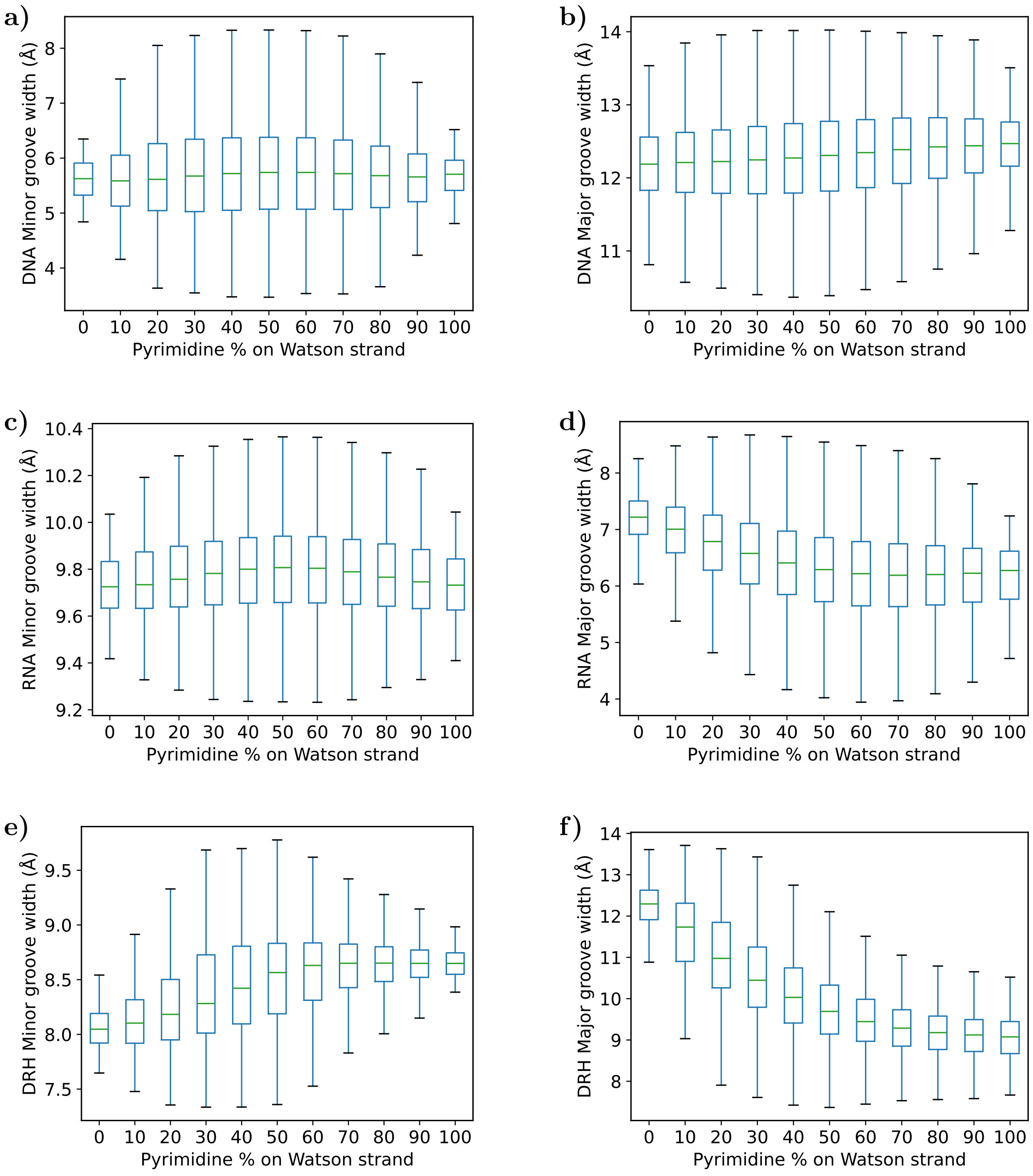

Figure S20: Major and minor groove widths as a function of pyrimidine content (on Watson or reference strand) in dsDNA, dsRNA, and DRH. **(a)** Minor groove widths in dsDNA, **(b)** Major groove widths in dsDNA, **(c)** Minor groove widths in dsRNA, **(d)** Major groove widths in dsRNA, **(e)** Minor groove widths in DRH, **(f)** Major groove widths in DRH. The statistics are obtained by exhaustively computing the groove widths for all decamers ($\approx$ one million sequences) embedded in fixed flanking contexts. The Watson phosphate between $5^{\text{th}}$ and $6^{\text{th}}$ is taken as the reference phosphate to compute the groove widths. The x-axis is the pyrimidine content in the decamer and the y-axis is the groove width in Å. The error bars show the standard deviation of groove widths for all sequences with a given pyrimidine content.

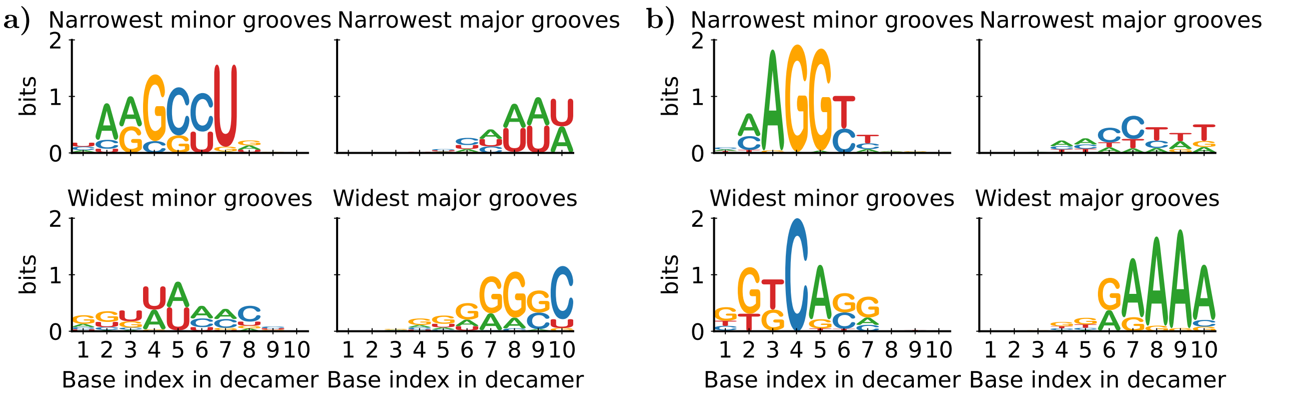

Figure S21: Sequence logos for sequences with extreme major and minor groove widths in **(a)** dsRNA and **(b)** DRH. The statistics are obtained from all decamers (≈ one million sequences) embedded in fixed flanking contexts. The x-axis is base index in the decamer with information content at that index on the y-axis. The Watson phosphate between $5^{th}$ and $6^{th}$ is taken as the reference phosphate.

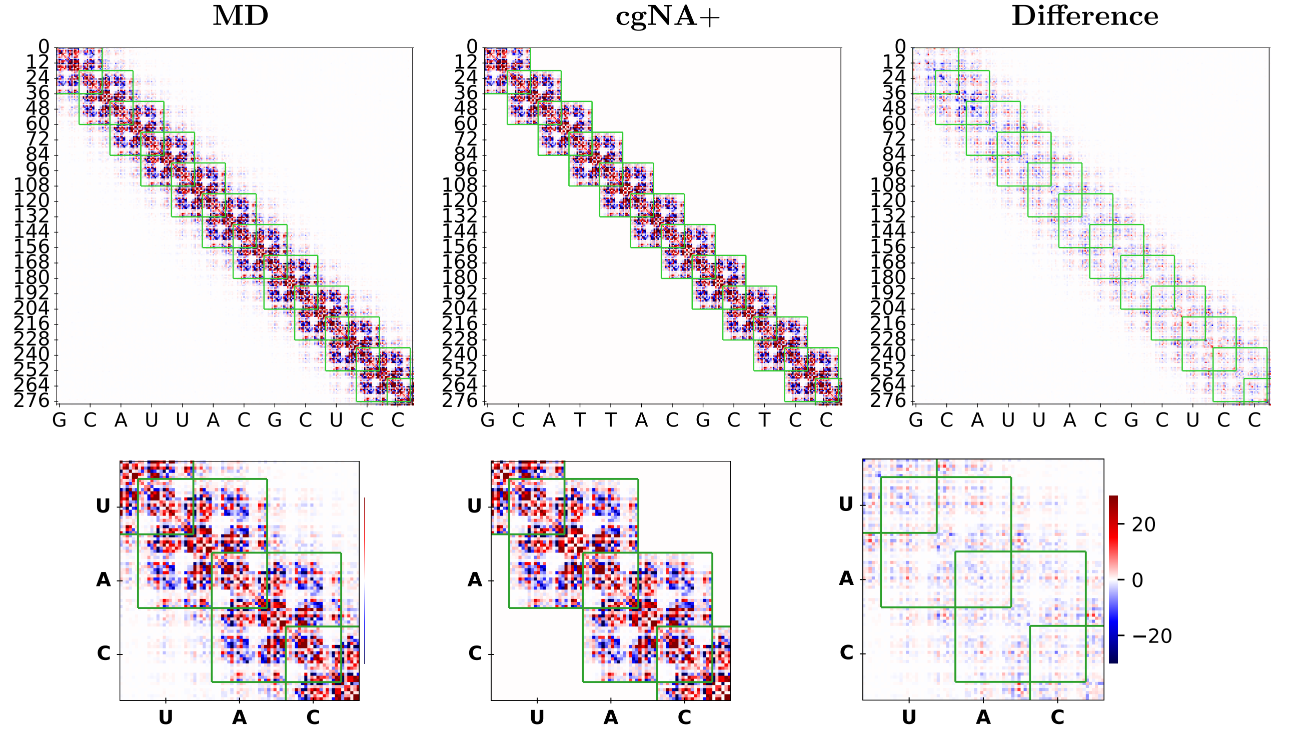

Figure S22: Comparison of stiffness matrices for a dsRNA sequence GCAUUACGCUCCG-GAGCGUAAUGC. The stiffness matrix estimated from MD simulations is shown on the left, the cgNA+ prediction in the center, and the element-wise difference between the MD-derived and model-predicted matrices on the right. Because the sequence is palindromic, only half of each matrix contains independent entries; accordingly, only this portion is displayed. The green outlines indicate the (42×42) blocks corresponding to the nearest-neighbour interaction approximation. The lower panels show magnified views of the corresponding matrices.A common color bar, ranging from -30 to 30, is used for all matrices displayed in the figure. The discrepancy between the MD-derived and cgNA+ matrices is quantified by the symmetrized Kullback–Leibler divergence per dof, which is 0.0081.

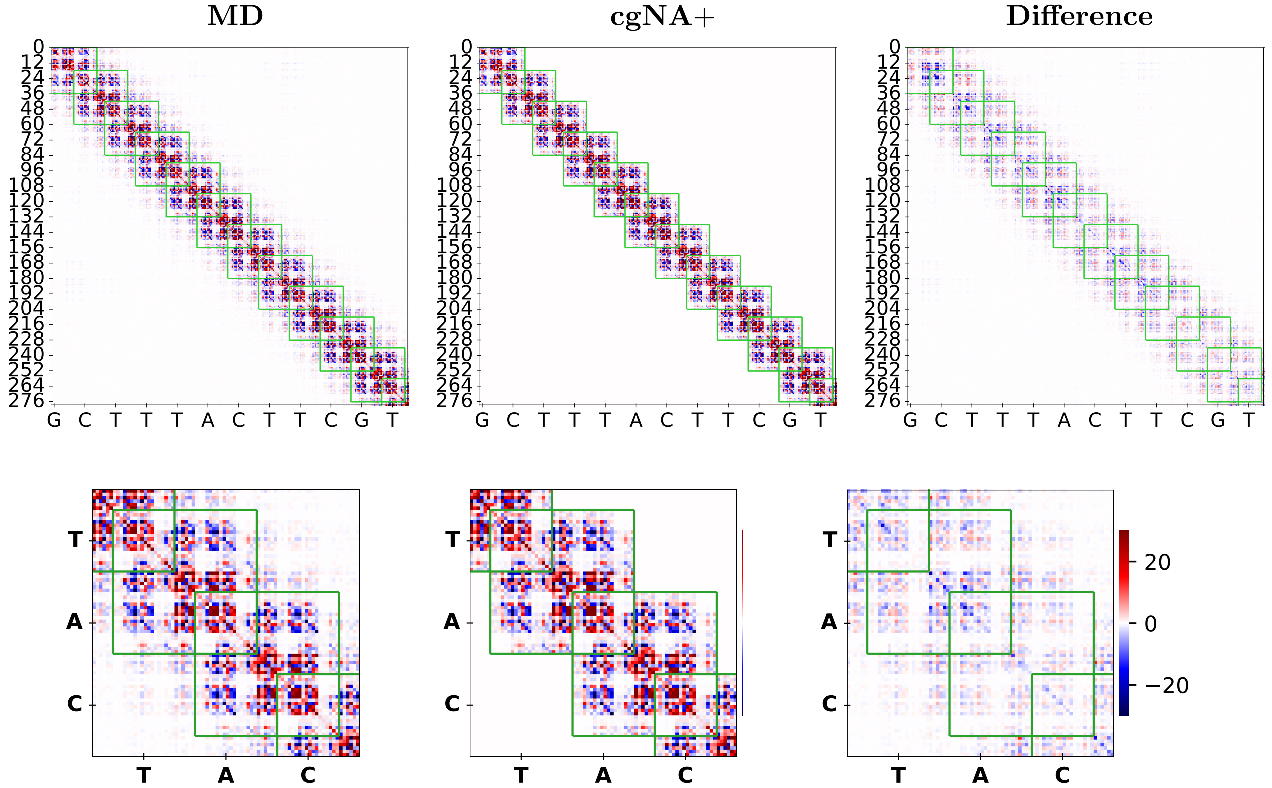

Figure S23: Comparison of stiffness matrices for a DRH sequence GCTTTACTTCGTACGAAG-TAAAGC. The stiffness matrix estimated from MD simulations is shown on the left, the cgNA+ prediction in the center, and the element-wise difference between the MD-derived and model-predicted matrices on the right. To be consistent with dsDNA and dsRNA versions of this plot, only half of the matrix is displayed. The green outlines indicate the (42×42) blocks corresponding to the nearest-neighbour interaction approximation. The lower panels show magnified views of the corresponding matrices. A common color bar, ranging from -30 to 30, is used for all matrices displayed in the figure. The discrepancy between the MD-derived and cgNA+ matrices is quantified by the symmetrized Kullback–Leibler divergence per dof, which is 0.0201.

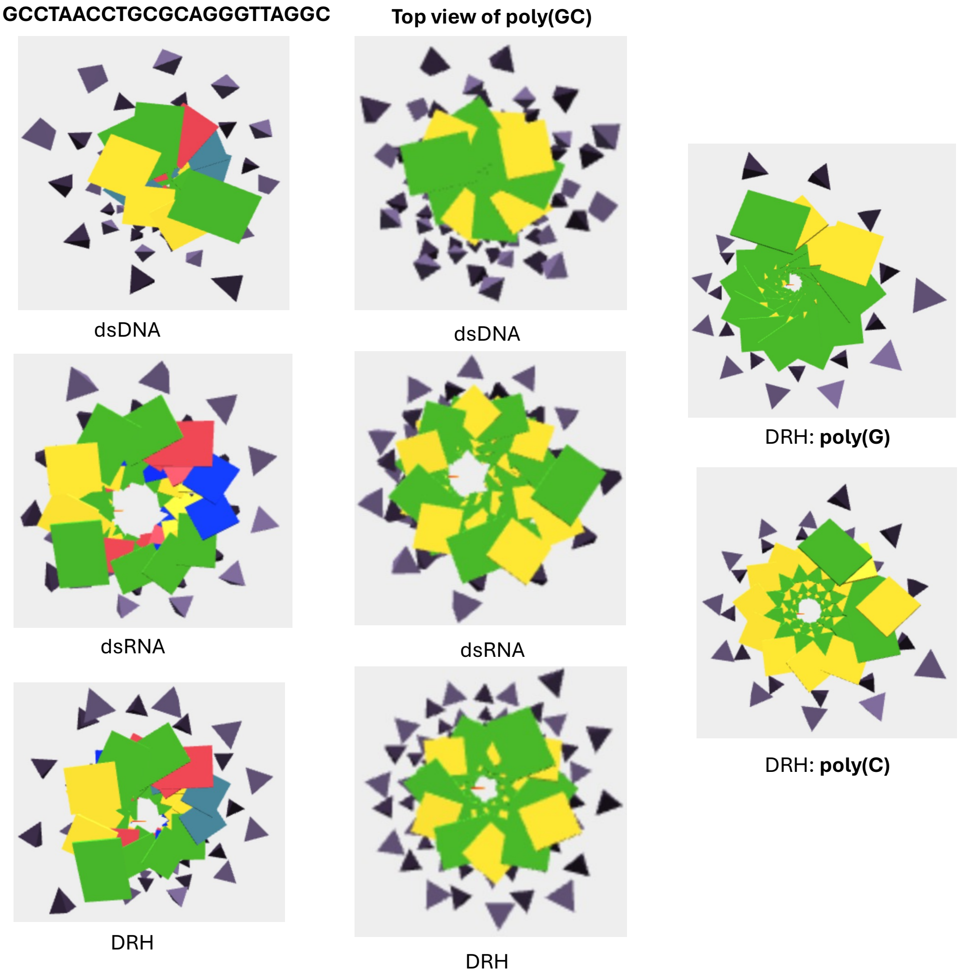

Figure S24: Top views of the 3D equilibrium shapes obtained from the `https://cgdnaweb.epfl.ch`. Left: GCCTAACCTGCGCAGGGTTAGGC in its dsDNA, dsRNA, and DRH versions. Center: $(GC)_{20}$ in its dsDNA, dsRNA, and DRH versions. The figure highlights the distinct helical diameters adopted by the sequence in the three duplex classes: dsDNA has the smallest diameter, consistent with B-form geometry; dsRNA has the largest diameter, consistent with A-form geometry; and DRH has an intermediate diameter, reflecting its mixed A/B-form character. Right: Compares the helical diameters of poly(C) and poly(G) for DRH, highlighting that the helical diameter is not simply the average of dsRNA and dsDNA, but is strongly modulated by sequence.

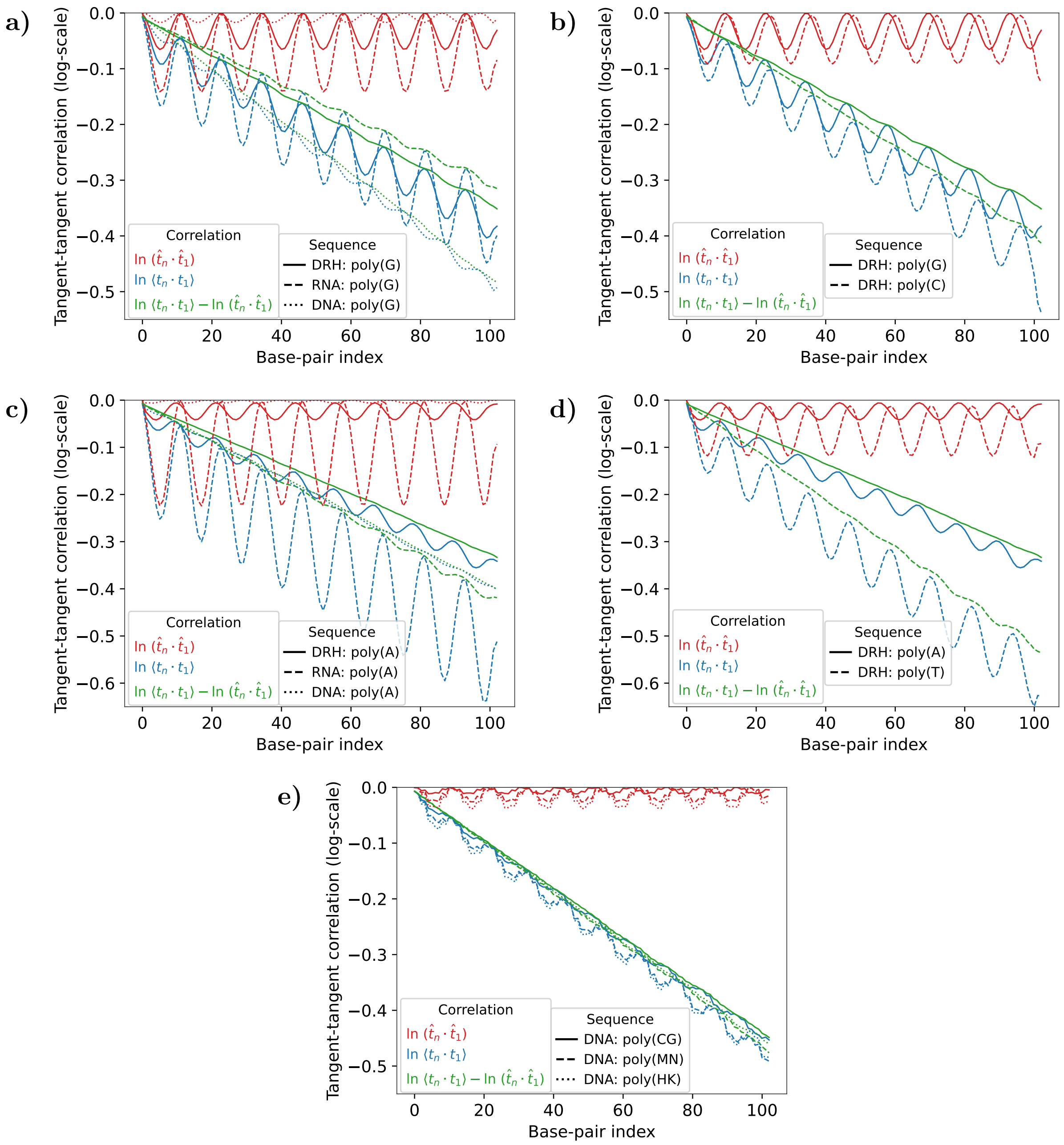

Figure S25: Comparison of the logarithm of the tangent–tangent correlation obtained from Monte Carlo sampling of the Gaussian pdfs inferred from cgNA+ for **(a)** poly(G) in dsDNA, DRH, and dsRNA versions, **(b)** poly(G) and poly(C) DRH version, **(c)** poly(A) in dsDNA, DRH, and dsRNA versions, **(d)** poly(A) and poly(T) DRH version, and **(e)** poly(CG) and its methylated poly(MN) and hydroxymethylated poly(HK) dsDNA versions. $\mathbf{t}_i$ denotes the unit tangent vector associated with base-pair frame $i$, $\hat{\mathbf{t}}_i$ denotes the unit tangent vector associated with base-pair frame $i$ in the groundstate, and $\langle \cdot \rangle$ denotes an average over the Monte Carlo ensemble. The oscillations in $\ln(\langle \mathbf{t}_i \cdot \mathbf{t}_1 \rangle)$ and $\ln\big(\hat{\mathbf{t}}_i \cdot \hat{\mathbf{t}}_1\big)$ reflect the helical diameter of the centerline in the equilibrium shape of dsNA. For dsDNA sequences, the centerline is close to straight, corresponding to a very small helical diameter characteristic of B-form geometry, whereas for dsRNA it adopts A-form geometry with a substantially larger helical diameter, and for DRH the behaviour is intermediate in a sequence-dependent manner. The helical diameter likely driven by roll, an inter-base-pair parameter (see Figure S8). The increase in roll upon CpG modification (see Figure S13) increases the helical diameter of poly(CG) as can be seen in **(e)**.

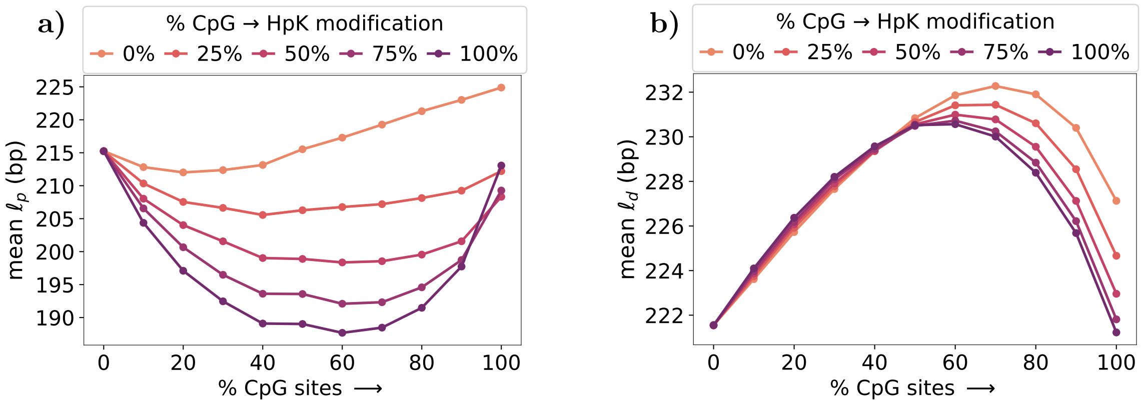

Figure S26: Persistence lengths, $\ell_p$ **(a)** and $\ell_d$ **(b)** as a function of $C_pG$ content and impact of symmetric $C_pG$ hydroxymethylation. Data points represent mean across random sequences of length 220 bps with varying $C_pG$ content in increments of 10% (10000 sequences for each). Within each set, 0%, 25%, 50%, 75%, or 100% of the $C_pG$ steps were randomly hydroxymethylated.

[18] Daiva Petkevičiūtė. A DNA coarse-grain rigid base model and parameter estimation from molecular dynamics simulations. *PhD thesis, No. 5520, EPFL, 10.5075/epfl-thesis-5520*, 2012.

[19] Rahul Sharma, Alessandro S Patelli, Lennart De Bruin, and John H Maddocks. cgNA+web: A visual interface to the cgNA+ sequence-dependent statistical mechanics model of double-stranded nucleic acids. *Journal of Molecular Biology*, page 167978, 2023.

[20] Harold Jeffreys. An invariant form for the prior probability in estimation problems. *Proceedings of the Royal Society of London. Series A. Mathematical and Physical Sciences*, 186(1007):453–461, 1946.

[21] W. K. Olson, A. A. Gorin, X. J. Lu, L. M. Hock, and V. B. Zhurkin. DNA sequence-dependent deformability deduced from protein–DNA crystal complexes. *Proceedings of the National Academy of Sciences*, 95(19):11163–11168, 1998.
